# Defects in translation-dependent quality control pathways lead to convergent molecular and neurodevelopmental pathology

**DOI:** 10.1101/2021.01.25.428167

**Authors:** Markus Terrey, Scott I. Adamson, Jeffrey H. Chuang, Susan L. Ackerman

## Abstract

Translation-dependent quality control pathways such as no-go decay (NGD), non-stop decay (NSD) and nonsense-mediated decay (NMD) govern protein synthesis and proteostasis by resolving non-translating ribosomes and preventing the production of potentially toxic peptides derived from faulty and aberrant mRNAs. However, how translation is altered and the *in vivo* defects that arise in the absence of these pathways are poorly understood. Here, we show that the NGD/NSD factors *Pelo* and *Hbs1l* are critical for cerebellar neurogenesis but expendable for survival of these neurons after development. Analysis of mutant embryonic fibroblasts revealed translational pauses, alteration of signaling pathways, and translational reprogramming. Similar effects on signaling pathways, the translatome and cerebellar development were observed upon deletion of the NMD factor *Upf2.* These data reveal that these quality control pathways that function to mitigate errors at distinct steps in translation can evoke similar cellular responses.

## Introduction

Regulation of gene expression is essential for cell growth and development. Although epigenetic mechanisms and transcriptional regulation are initial steps in gene expression, translation and its regulation have emerged as major hubs to control the production of functional proteins. In fact, translation is intricately coordinated with the degradation of faulty mRNAs and their resulting peptide products to balance gene expression and proteostasis (Collart and Weiss, 2019). Numerous studies have revealed that mRNA levels show limited correlation with protein levels, particularly when cells undergo dynamic transitions (Abreua et al., 2009; Kristensen et al., 2013; Liu et al., 2016; Maier et al., 2009). Indeed, post-transcriptional mechanisms, such as translation, play vital roles in rapidly altering gene expression and the signaling pathways that mediate cell identity and cell fate changes necessary for mammalian development (Blair et al., 2017; Blanco et al., 2016; Fujii et al., 2017; Gabut et al., 2020; Kong and Lasko, 2012; Rodrigues et al., 2020; Signer et al., 2014; Tahmasebi et al., 2019).

The process of translation is highly organized, and ribosomes need to accurately perform the steps of translation initiation, elongation and termination. However, multiple factors can perturb translation including secondary structures of mRNAs, amino acid limitations, tRNA deficiencies, rare codons, interactions of nascent peptides with the ribosome, chemically damaged mRNAs, and cleaved or aberrant mRNAs (Brandman and Hegde, 2016; Brule and Grayhack, 2017; Buhr et al., 2016; Drummond and Wilke, 2008; Hu et al., 2009; Simms et al., 2014; Spencer et al., 2012; Thommen et al., 2017; Wolf and Grayhack, 2015; Yu et al., 2015). In turn, these defects may trigger translation-dependent quality control pathways including non-stop decay (NSD), no-go decay (NGD) or nonsense-mediated mRNA decay (NMD) to release ribosomes, eliminate potentially toxic or faulty peptide products, and co-translationally decay problematic or aberrant mRNAs (Collart and Weiss, 2019).

Defects in translation and their translation-dependent quality control pathways impair cellular homeostasis and have been linked to proteotoxicity, changes in synaptic function, and neurodegeneration (Choe et al., 2016; Chu et al., 2009; Huang et al., 2017; Ishimura et al., 2014; Johnson et al., 2019; Kapur et al., 2020; Martin et al., 2020; Notaras et al., 2019; Terrey et al., 2020; Yonashiro et al., 2016). Here, we demonstrate that the ribosome rescue factors *Pelo* and *Hbs1l*, which are implicated in NSG and NGD, are critical for embryonic and brain development, but dispensable for neuronal survival in the adult brain. Our analysis of *Pelo*- and *Hbs1l*-deficient fibroblasts reveal translational reprogramming of multiple pathways. Inhibition of NMD via deletion of *Upf2* resulted in strikingly similar effects on the translatome, signaling pathways, and neurogenesis. Our data reveal that defects in translation-dependent quality control pathways, which mitigate errors in translation to prevent the production of defective peptide products from aberrant mRNAs, can trigger similar cellular responses and neurodevelopmental abnormalities.

## Results

### *Hbs1l* is required for embryogenesis

Multiple neurological abnormalities, including defects in motor control, were recently described in a patient with biallelic mutations in *Hbs1l* (O’Connell et al., 2019). Alternative splicing of *Hbs1l* produces transcripts that encode two distinct proteins (*Figure 1A*). Levels of full length *Hbs1l* (*Hbs1l*-V1 and Hbs1 in human and yeast, respectively) were dramatically decreased in *Hbs1l* patient fibroblasts (O’Connell et al., 2019). The levels of the shorter isoform II (*Hbs1l*-V3 in human), which is encoded by the first 4 exons of full length *Hbs1l* and a unique last exon (‘exon 5a’) located between exon 4 and exon 5 of the *Hbs1l* locus, were relatively unaffected in the *Hbs1l* patient fibroblasts (O’Connell et al., 2019). In contrast to the translation-dependent quality control function of *Hbs1l*, previous studies suggested that isoform II of *Hbs1l* is likely an ortholog of the Saccharomyces cerevisiae protein SKI7 (Brunkard and Baker, 2018; Kalisiak et al., 2017; Marshall et al., 2018), which is involved in global mRNA turnover (Kalisiak et al., 2017). Although an additional splice variant (*Hbs1l*-V2 in human) which lacks the third coding exon is annotated in the human transcriptome (Mills et al., 2016; O’Connell et al., 2019), this splice variant is not annotated in mice, and we were unable to detect it by RT-PCR.

**Figure 1.**
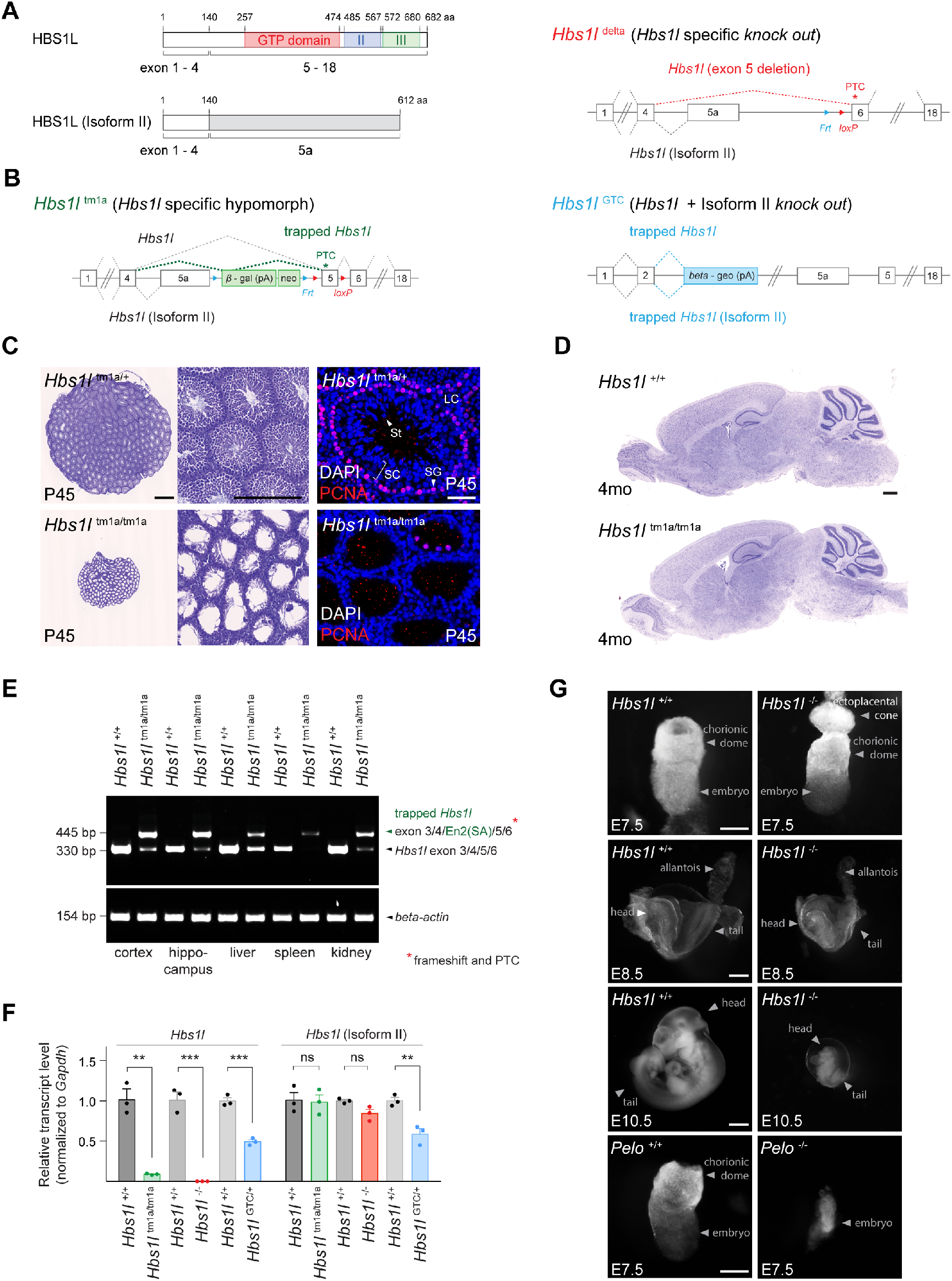
*Hbs1l* is required for embryogenesis. (A) Domain structure of HBS1L and Isoform II and the exons encoding the two splice variants. (B) Design of *Hbs1l* loss-of-function alleles that target *Hbs1l* and Isoform II. *Hbs1l*^tm1a^ allele, *Hbs1l* specific gene trap (*hypomorph*); *Hbs1l*^delta^ allele, *Hbs1l* specific deletion of exon 5 (*knock out*); and *Hbs1l*^GTC^ allele, *Hbs1l* gene trap to target *Hbs1l* and isoform II (*knock out*). (C) Cresyl violet-stained cross-sections of testis from P45 control (*Hbs1l*^tm1a/+^) and *Hbs1l*^tm1a/tm1a^ mice. Immunofluorescence was performed with antibodies to proliferating cell nuclear antigen (PCNA, red) and sections were counterstained with DAPI. (D) Cresyl violet-stained sagittal brain section from 4-month-old control (*Hbs1l*^+/+^) and *Hbs1l*^tm1a/tm1a^ mice. (E) Splicing analysis of correctly spliced *Hbs1l* and trapped *Hbs1l* transcripts in various tissues from 4-month-old control (*Hbs1l*^+/+^) and *Hbs1l*^tm1a/tm1a^ mice. *β-actin* was used as an input control. (F) Quantitative RT-PCR analysis of *Hbs1l* and isoform II using cDNA from E8.5 embryos. Data were normalized to *Gapdh* and the fold change in gene expression is relative to that of controls (*Hbs1l*^+/+^) from each cross. Data represent mean + SEM. (G) Bright field images of control (*Hbs1l*^+/+^ and *Pelo*^+/+^), *Hbs1l*^-/-^ and *Pelo*^-/-^ embryos at E7.5, E8.5 and E10.5. Scale bars: 500μm and 200μm (higher magnification), 50μm (immunofluorescence image) (C); 500μm (D); 100μm (E7.5), 200μm (E8.5), and 2mm (E10.5) (G). PTC, premature termination codon; Frt, flippase-mediated recombination site; loxP, *Cre* recombinase-mediated recombination site; En2(SA), splice acceptor of mouse *Engrailed-2* exon 2; SC, spermatocytes; SG, spermatogonia; St, spermatids; LC, Leydig cells. t-tests were corrected for multiple comparisons using Holm-Sidak method (E). ns, not significant; ** *P* ≤ 0.01; *** *P* ≤ 0.001.

To study the neurological function of *Hbs1l* in mice, we first examined an *Hbs1l* allele (*Hbs1l*^tm1a^) with a gene trap cassette inserted between ‘exon 5a’ and exon 5 (*Figure 1B*). As previously reported, homozygous *Hbs1l*^tm1a/tm1a^ mice were viable, but male mice were infertile (O’Connell et al., 2019). Histological analysis of the *Hbs1l*^tm1a/tm1a^ testis at postnatal day P45 revealed a dramatic loss of mitotically active (PCNA^+^) spermatogonia, as well as spermatocytes and spermatids that normally differentiate from these cells (*Figure 1C*). However, no overt defects were observed in the *Hbs1l*^tm1a/tm1a^ brain (*Figure 1D*).

Residual levels of *Hbs1l* were still present in various tissues from *Hbs1l*^tm1a/tm1a^ mice (O’Connell et al., 2019). In agreement, *Hbs1l* transcripts spliced into the gene trap cassette in all tested tissues; however, correctly spliced *Hbs1l* transcripts were still detected in several tissues (*Figure 1E*). Thus, to completely eliminate expression of *Hbs1l*, we ubiquitously deleted exon 5 in *Hbs1l*^tm1a^ mice to generate *Hbs1l*^-/-^ mice (*Figure 1B*). In contrast to embryonic day (E) 8.5 *Hbs1l*^tm1a/tm1a^ embryos, which still expressed 9% of the wild type levels of *Hbs1l* mRNA, expression of *Hbs1l* was not detected in E8.5 *Hbs1l*^-/-^ embryos (*Figure 1F*). Expression of *Hbsl1* isoform II was not significantly changed in homozygous embryos of either allele, as predicted (*Figure 1F*). In contrast to hypomorphic *Hbs1l*^tm1a/tm1a^ mice, *Hbs1l*^-/-^ embryos failed to develop after E8.5 and could not be recovered at E11.5 (*Figure 1G*, *Supplementary file 1*). These results demonstrate that *Hbs1l* is necessary for embryonic development, however embryos lacking *Hbs1l* develop longer than embryos deficient for *Pelo*, the binding partner of HBS1L, which die by E7.5 (Adham et al., 2003) (*Figure 1G*).

To determine if isoform II of *Hbs1l* is also necessary for embryonic viability, we utilized an additional allele (*Hbs1l*^GTC^) with a gene trap cassette located in intron 2 (*Figure 1B*). Expression of *Hbs1l* and isoform II transcripts in E8.5 heterozygous *Hbs1l*^GTC/+^ embryos was reduced by 51% and 41%, respectively (*Figure 1F*). Homozygous *Hbs1l*^GTC/GTC^ embryos were not recovered at E6.5 (*Supplementary file 1*) demonstrating that they died even before *Pelo*^-/-^ or *Hbs1l*^-/-^ embryos.

The early embryonic lethality of *Hbs1l*^GTC/GTC^ embryos suggests that the *Hbs1l* isoforms are likely functionally distinct, and that their loss causes additive or synergistic defects during embryogenesis.

### *Hbs1l* is required for cerebellar development

Consistent with transcriptome data from a brain RNA sequencing database (Zhang et al., 2014), we observed expression of *Hbs1l* in multiple cell types of the brain (*Figure 2A*). Expression of HBS1L and its binding partner PELO was observed throughout and after cerebellar development (*Figure 2A, B and C*). However, levels of HBS1L and PELO decreased in the postnatal (P)14 cerebellum after the completion of development, which was similar to the levels observed in the whole brain (*Figure 2B and C, Figure 2 - figure supplement 1A*). To begin to investigate the role of *Hbs1l* in the brain, we deleted this gene in the developing cerebellum and midbrain by crossing the floxed allele to En1-Cre (*En1*^cre/+^) mice (Kimmel et al., 2000). Differences in cerebellar size between mutant and control embryos were already apparent by E13.5 (*Figure 2D*). Although the trilaminar structure of the cerebellum appeared normal in En1-Cre; *Hbs1l*^cKO^ mice at postnatal day P21, cerebellar foliation was delayed in P0 mutant mice, and secondary fissures failed to form compared to control mice (*Figure 2D*) and consistent with previous studies no cerebellar abnormalities were observed in En1-Cre mice (Dong and Kwan, 2020; Guo et al., 2010; Li et al., 2002; Sgaier et al., 2007; Tripathi et al., 2008).

**Figure 2.**
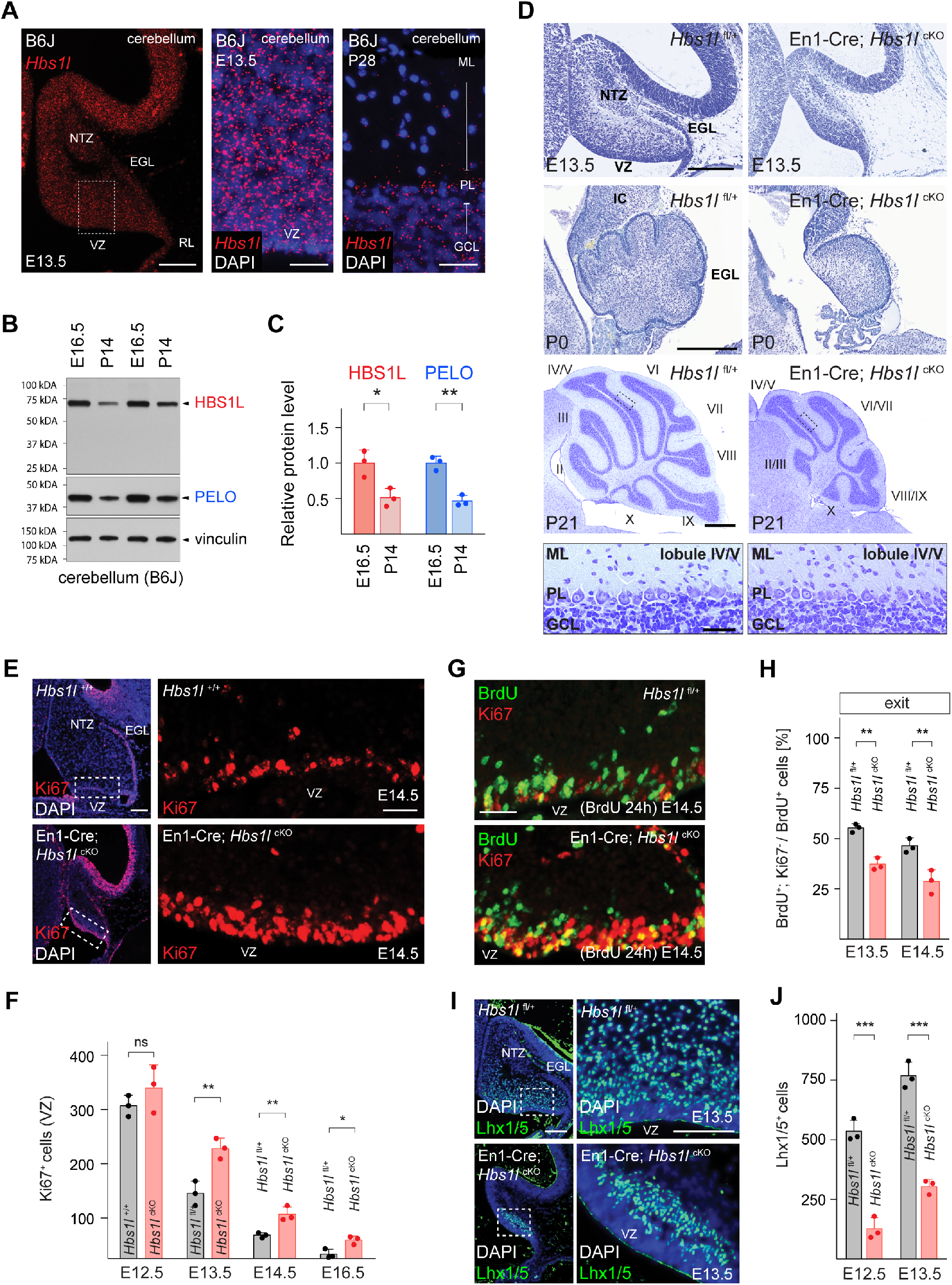
*Hbs1l* is required for cerebellar development. (A) *In situ* hybridization of *Hbs1l* mRNA (red) on cerebellar sections from E13.5 and P28 control (B6J) mice. Sections were counterstained with DAPI. (B) Western blot analysis using cerebellar lysates from B6J mice. Vinculin was used as an input control. (C) The relative protein levels of HBS1L and PELO were normalized to levels of vinculin and protein levels are relative to those of E16.5 B6J cerebella. (D) Parasagittal (E13.5) and sagittal (P0 and P21) cerebellar sections from control (*Hbs1l*^fl/+^) and En1- Cre; *Hbs1l*^cKO^ mice stained with cresyl violet. Higher magnification images of lobules IV/V at P21 are shown below each genotype. Cerebellar lobules are indicated by Roman numerals. (E) Immunofluorescence using antibodies to Ki67 (red) on cerebellar section from E14.5 control (*Hbs1l*^+/+^) and En1-Cre; *Hbs1l*^cKO^ embryos. Sections were counterstained with DAPI and higher magnification images of boxed area are shown. (F) Number of cerebellar VZ-progenitors (Ki67^+^ cells) from control (*Hbs1l*^+/+^ or *Hbs1l*^fl/+^) and En1-Cre; *Hbs1l*^cKO^ embryos. Data represent mean + SD. (G) Immunofluorescence using antibodies to BrdU (green) and Ki67 (red) on cerebellar sections from E14.5 control (*Hbs1l*^fl/+^) and En1-Cre; *Hbs1l*^cKO^ embryos to determine the fraction of cells that exited the cell cycle. Embryos were injected with BrdU 24 hours prior to harvest. (H) Percentage of cerebellar VZ-progenitors that exited the cell cycle (BrdU^+^, Ki67^-^ cells). Data represent mean + SD. (I) Immunofluorescence using antibodies to Lhx1/5 on cerebellar sections from E13.5 control (*Hbs1l*^+/+^) and En1-Cre; *Hbs1l*^cKO^ embryos. Sections were counterstained with DAPI and higher magnification images of boxed areas are shown. (J) Number of cerebellar Purkinje cell precursors (Lhx1/5^+^ cells). Data represent mean + SD. Scale bars: 100μm and 20μm (higher magnifications) (A); 200μm (E13.5 and P0), 500μm and 50μm (higher magnification) (P21) (D); 100μm and 20μm (higher magnification) (E); 20μm (G); 100μm and 50μm (higher magnification) (I). VZ, ventricular zone; NTZ, nuclear transitory zone; RL, rhombic lip; EGL, external granule cell layer; ML, molecular cell layer; PL, Purkinje cell layer; GCL, granule cell layer; IC, inferior colliculus. t-tests were corrected for multiple comparisons using Holm-Sidak method (C, F, H, J). ns, not significant; * *P* ≤ 0.05; ** *P* ≤ 0.01; *** *P* ≤ 0.001.

Lineage-restricted GABAergic precursors are generated beginning at ∼E10.5 from progenitors in the ventricular zone (VZ) of the developing cerebellum (Ju et al., 2016; Leto et al., 2012). The pool of progenitors declines as progenitors either exit the cell cycle to generate precursors or retract from the VZ to form a secondary germinal zone in the prospective white matter to transition from neurogenesis to gliogenesis (Leto et al., 2012; Vong et al., 2015; Wizeman et al., 2019). Immunofluorescence with antibodies to the cell cycle-associated protein Ki67 demonstrated a loss of progenitors between E12.5 to E16.5 in both the control and En1-Cre; *Hbs1l*^cKO^ cerebella.

However, the number of progenitors remained higher in En1-Cre; *Hbs1l*^cKO^ compared to control cerebella (*Figure 2E and F*), suggesting that *Hbs1l*-deficient progenitors may aberrantly proliferate. In agreement, we observed a higher fraction of VZ-progenitors in S-phase in the E12.5 and E13.5 mutant cerebella by pulse labeling with BrdU for 30 minutes and performing co-immunofluorescence with BrdU and Ki67 antibodies (*Figure 2 - figure supplement 1B and C*). Immunofluorescence with antibodies to the M-Phase marker phospho-histone 3 (pH3) and the general cell cycle marker PCNA demonstrated an increase of VZ-progenitors also in M-phase in the mutant cerebellum (*Figure 2 - figure supplement 1D*).

To test if *Hbs1l*-deficient progenitors are able to exit the cell cycle, which is necessary to generate lineage-restricted precursors, we labeled control and mutant embryos at E12.5 or E13.5 with BrdU and then determined the fraction of BrdU^+^ cells that had left the cell cycle (do not express Ki67) twenty-four hours after labeling. Fewer VZ-progenitors exited the cell cycle in En1-Cre; *Hbs1l*^cKO^ relative to control cerebella (*Figure 2G and H*). Concordantly, the number of VZ-derived Lhx1/5^+^ Purkinje cells and Pax2^+^ interneuron precursors was reduced to ∼32% and ∼34% in En1-Cre; *Hbs1l*^cKO^ between E12.5 to E16.5 compared to control cerebella (*Figure 2I and J*, *Figure 2 - figure supplement 1E and F*), suggesting that *Hbs1l* deficient VZ-progenitors remain proliferative at the expense of differentiation.

Cerebellar glutamatergic precursors are generated beginning at ∼E10.5 from the rhombic lip (RL), a germinal zone located in the caudal region of the cerebellar primordia. Between E10.5-12.5 the RL gives rise to Tbr1^+^ cells that migrate subpially to take up residence in the nuclear transitory zone and will develop into deep cerebellar neurons (Fink et al., 2006). Following Tbr1^+^ cell production, proliferating granule cell precursors emerge from the RL and form the external granule cell layer (EGL) (Chung et al., 2010). As observed for VZ-derived neuronal precursors, deletion of *Hbs1l* also decreased progeny generated from the RL. Immunofluorescence with antibodies to Tbr1, revealed fewer Tbr1^+^ cells in the E14.5 mutant cerebella compared to controls (*Figure 2 - figure supplement 1G*). In addition, the EGL of En1-Cre; *Hbs1l*^cKO^ at E13.5 to E16.5 contained only ∼45% of the granule cell precursors compared to controls (*Figure 2 - figure supplement 1H and I*).

Granule cell precursors that derive from the RL continue to proliferate in the EGL to generate additional precursors that exit the cell cycle postnatally prior to migrating to the IGL. To determine if the decrease in granule cell precursors was due to a failure of progenitors in the RL to generate sufficient numbers of granule cell precursors, or a failure of granule cell precursors to proliferate, we labeled E12.5 and E13.5 wild type and mutant embryos with BrdU and then determined the number of BrdU^+^ cells in the EGL twenty-four hours after labeling. Fewer BrdU^+^ cells were present in the En1-Cre; *Hbs1l*^cKO^ EGL compared to that of controls indicating that the mutant rhombic lip generates fewer granule cell precursors (*Figure 2 - figure supplement 1J and K).* However, cell cycle exit of these precursors did not vary between mutant and control cells (*Figure 2 - figure supplement 1L*). Furthermore, no difference in proliferation (S- and M-phase) was observed when granule cell precursors were analyzed at E16.5 or P5 (*Figure 2 - figure supplement 1M and N*). Together, these data suggest that while *Hbs1l* is required for the initial production of granule cell precursors, it is dispensable for the subsequent cell cycle progression of these precursors in the EGL. In agreement, genetic ablation of *Hbs1l* by Atoh1*-*Cre which specifically deletes in cerebellar granule cell precursors in the EGL beginning at ∼E16.5 (Lorenz et al., 2011; Pan et al., 2009; Qiu et al., 2010; Wojcinski et al., 2019), did not impair development of these cells (*Figure 2 - figure supplement 2A*).

*Hbs1l* was also required for gliogenesis in the developing cerebellum. Cerebellar gliogenesis starts at ∼E18 and continues during postnatal development during which time progenitors that have retracted from the VZ switch from a neurogenic to gliogenic fate and produce oligodendrocytes and astrocytes (Götz and Huttner, 2005; Vong et al., 2015). Immunofluorescence with antibodies to Olig2, a marker of oligodendroglial progenitors which gives rise to oligodendrocytes and astrocytes (Chung et al., 2013; Tatsumi et al., 2018), revealed the number of these cells was reduced in P5 En1-Cre; *Hbs1l*^cKO^ cerebella (*Figure 2 - figure supplement 1O*). Together these data indicate that *Hbs1l* is required for the generation of multiple cell types in the developing cerebellum.

We have previously identified a mutation in the common C57BL/6J (B6J) strain that partially disrupts processing of the brain-specific arginine tRNA, *n-Tr20* (*n-Tr20*^B6J/B6J^). This processing defect in turn reduces the pool of available tRNA^Arg^_UCU_, leading to ribosome pausing at the A-site at AGA codons in cerebellar mRNAs (Ishimura et al., 2014). *Hbs1l*^cKO^ mice were generated with the B6J-associated mutation in *n-Tr20*. To test if the *n-Tr20* deficiency influenced the developmental defects observed in the absence of *Hbs1l*, we either restored *n-Tr20* to wild type levels or completely deleted *n-Tr20* in *Hbs1l*^cKO^ mice. Wild type expression of the tRNA (*n-Tr20*^B6N/B6N^) did not rescue defects in En1-Cre; *Hbs1l*^cKO^ cerebella, nor did complete loss of *n-Tr20* (*n-Tr20*^-/-^) cause developmental defects in Atoh1-Cre; *Hbs1l*^cKO^ cerebella (*Figure 2 - figure supplement 2A*). In addition, neither loss of *Hbs1l* or *Pelo* affected cell survival of terminally differentiated granule cells in 9-month-old mice even in the presence of the *n-Tr20* deficiency (*Figure 2 - figure supplement 2B, C, D and E*), indicating that *Hbs1l* and *Pelo* do not respond to AGA pausing.

### Ribosome pausing correlates with pathology in *Hbs1l*-deficient mice

To determine if the developmental defects that occur upon loss of *Hbs1l* are accompanied by alterations in translation elongation, we performed ribosome profiling on wild type and *Hbs1l*^-/-^ embryos at E8.5. Ribosome protected fragments (RPF) mapped primarily to the protein coding sequence of genes in both wild type and mutant embryos (*Figure 3A*). Using the previously described methodology (Ishimura et al., 2014), we found a total of ∼1300 sites with significant (z-score ≥ 10) increases in local ribosome occupancy (‘ribosome pauses’) in wild type and mutant embryos (*Figure 3B*, *Supplementary file 2*). About 40% of the ribosome pauses, which mapped to 319 genes, were shared between genotypes suggesting they occurred independently of the loss of *Hbs1l*. Ten percent of ribosome pauses (mapped to 107 genes) were found only in wild type embryos. Differential gene expression analysis of the translatome using the ribosome protected fragments (DE RPF, *Supplementary file 3*) indicated translation of these genes was decreased in mutant embryos (*Figure 3C*), which may contribute to the apparent specificity of these pauses to wild type embryos. Strikingly, 50% of ribosome pauses (mapped to 459 genes) were uniquely observed in *Hbs1l*^-/-^ embryos and like other pauses, occurred primarily in protein coding sequences (*Supplementary file 2*). Similar to the metagene (*Figure 3A*), we didn’t observe genotype-dependent differences in the ribosome occupancy in different gene regions (i.e. untranslated 5’ or 3’ region). Only 9% of genes associated with *Hbs1l*^-/-^-specific ribosome pauses were differentially translated in *Hbs1l*^-/-^ embryos (DE RPF *Hbs1l*^-/-^) and translation of 26 genes and 15 genes was decreased and increased, respectively (*Figure 3 - figure supplement 1A*).

**Figure 3.**
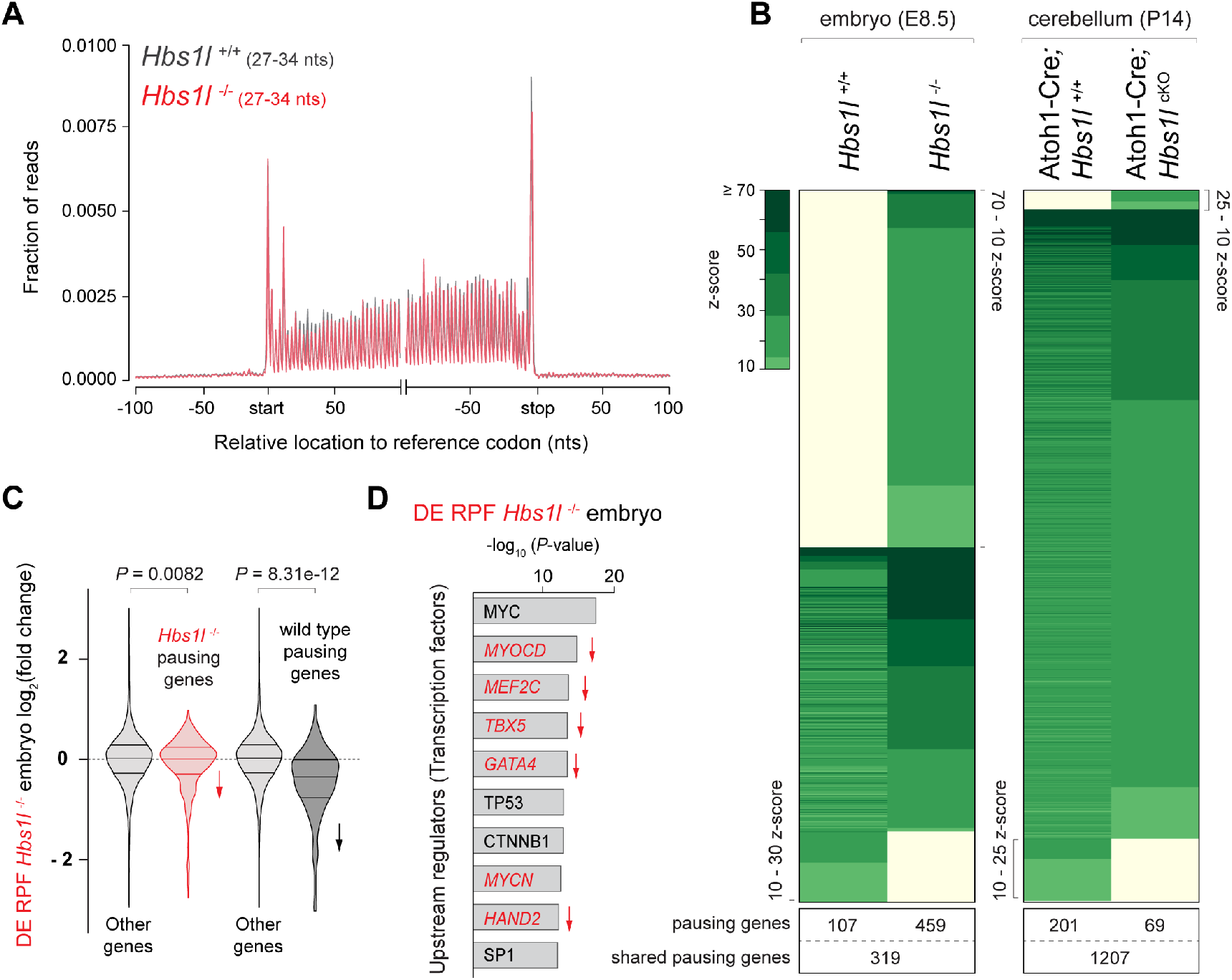
Ribosome pausing correlates with pathology in *Hbs1l* deficient mice. (A) Metagene profiles of RPFs from E8.5 control (*Hbs1l*^+/+^, grey traces) and *Hbs1l*^-/-^ (red traces) embryos. (B) Analysis of significantly increased local ribosome occupancy (z-score ≥ 10, pause site detected in all three replicates) from E8.5 wild type (*Hbs1l*^+/+^) and *Hbs1l*^-/-^ embryos (left) or P14 control (Atoh1-Cre; *Hbs1l*^+/+^) and Atoh1-Cre; *Hbs1l*^cKO^ cerebella (right). The number of genes that pause sites map to is shown below for each genotype. (C) All translated genes (DE RPF *Hbs1l*^-/-^) from E8.5 *Hbs1l*^-/-^ embryos were compared to the translation of genes which contained pauses specific to either *Hbs1l*^-/-^ or wild type embryos. Downward direction of arrow indicates significant reduction in translation of pausing genes in E8.5 *Hbs1l*^-/-^ embryos relative to wild type embryos. (D) Identification of upstream regulators using Ingenuity Pathway Analysis (IPA) of differentially translated genes of E8.5 *Hbs1l*^-/-^ embryos (DE RPF *Hbs1l*^-/-^). Transcription factors that are involved in heart development are shown in red. Downward direction of arrow indicates predicated activity (downregulation) of transcription factors. RPFs, ribosome-protected fragments; nts, nucleotides. Wilcoxon rank-sum test was used to determine statistical significance (C).

Loss of *Hbs1l* did not affect survival of granule cells (*Figures 2 - figure supplement 2B and C*), which constitute the vast majority of the cellular content of the cerebellum. Thus, to determine if elongation defects correlate with pathogenesis in *Hbs1l* mutant tissues, we also performed ribosome profiling on the cerebellum of P14 control and Atoh1-Cre; *Hbs1l*^cKO^ mice (*Supplementary file 2*). We observed more ribosome pausing sites (∼2700) in the cerebellum than in embryos, likely due to the higher amount of input RNA and sequencing depth. Consistent with the lack of a genetic interaction between the *Hbs1l* and the *n-Tr20* mutation, no significant increase in ribosome occupancy on A-site AGA codons was observed in *Hbs1l*^-/-^ cerebella (*Supplementary file 4*). Unlike the high percentage of pauses that were unique to *Hbs1l*^-/-^ embryos, only 2% of ribosome pauses (mapped to 69 genes) were unique to the mutant cerebellum (*Figure 3B*). In addition, the z-scores (*‘*pause scores’) for *Hbs1l*-specific pauses were significantly higher in mutant embryos than the mutant cerebellum (*Figure 3 - figure supplement 1B*).

Mutant embryos also exhibited larger changes in the translatome (DE RPF) than the mutant cerebellum (*Figure 3 - figure supplement 1C*), indicating that defects in translation also correlate with pathology. Kyoto Encyclopedia of Genes and Genomes (KEGG) and Ingenuity Pathway Analysis (IPA) of differentially translated genes (DE RPF *Hbs1l*^-/-^, adj. *P* ≤ 0.05) in *Hbs1l^-/-^* embryos revealed that downregulated genes were significantly enriched for heart/cardiac muscle contraction and calcium signaling (*Figure 3 - figure supplement 1D and E*). In agreement, upstream regulator analysis predicted downregulation of multiple transcription factors required for heart development (Cui et al., 2018; Molkentin et al., 1997; Muñoz-Martín et al., 2019; Steimle and Moskowitz, 2017) (*Figure 3D*). Together these data suggest that HBS1L deficiency may cause defects in the embryonic heart, one of the first organs to begin developing in the mouse embryo.

### Loss of *Pelo/Hbs1l* alters translation regulation and reprograms the translatome

Our embryonic data suggested that defects in translation modulate the translatome. Because *Pelo*^-/-^ embryos could not be profiled due to their early embryonic lethality, we conditionally deleted *Pelo* or *Hbs1l* in primary mouse embryonic fibroblasts (MEFs) using a tamoxifen-inducible Cre transgene (CAG-Cre^ER^) to compare changes in translation upon loss of *Pelo* and *Hbs1l* (*Figure 4A*). Consistent with previous studies (Juszkiewicz et al., 2020a; O’Connell et al., 2019), deletion of *Hbs1l* was accompanied by decreased levels of PELO protein, but not its mRNA levels (*Figure 4B, Figure 4 - figure supplement 1A, B and C*). In addition, we found deletion of *Pelo* also led to the loss of HBS1L without altering *Hbs1l* mRNA levels, suggesting protein degradation of PELO or HBS1L occurs in the absence of either interacting partner (*Figure 4B, Figure 4 - figure supplement 1A, B and C*).

**Figure 4.**
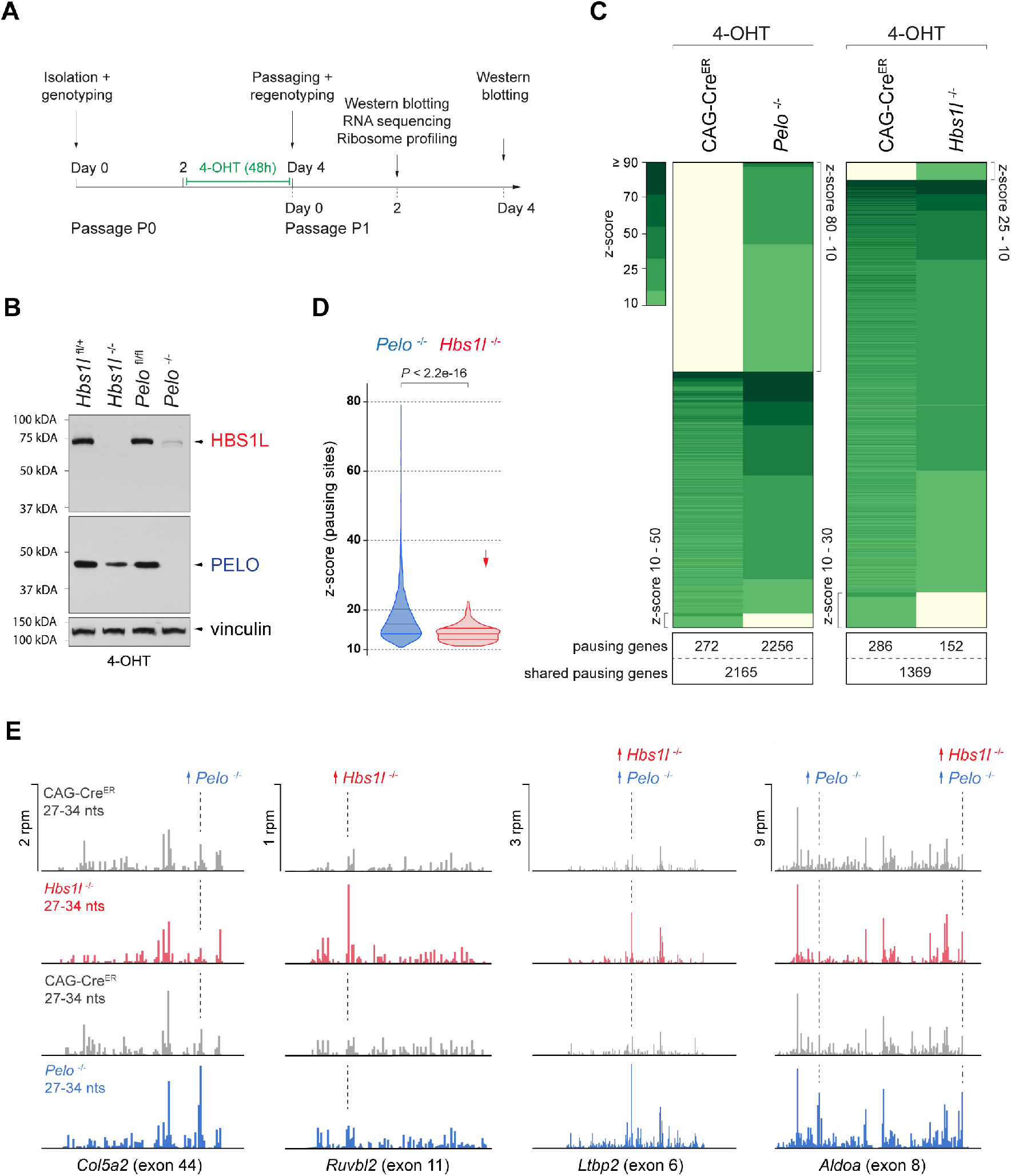
Loss of *Pelo* induces greater defects in translation elongation then loss of *Hbs1l*. (A) Experimental strategy for *in vitro* studies using tamoxifen (4-OHT) treatment of primary mouse embryonic fibroblasts (MEFs). (B) Western blot analysis of HBS1L and PELO using MEF lysates from tamoxifen-treated control (*Hbs1l*^fl/+^ or *Pelo*^fl/fl^), *Hbs1l*^-/-^ and *Pelo*^-/-^ cells. Vinculin was used as an input control. (C) Analysis of significantly increased local ribosome occupancy (z-score ≥ 10, pause site detected in all three replicates) from tamoxifen treated control (CAG-Cre^ER^) and *Pelo*^-/-^ (left), or control and *Hbs1l*^-/-^ cells (right). The number of genes that pause sites map to is shown below for each genotype. (D) Comparison of the z-scores (‘pause score’) for pauses observed in *Pelo*^-/-^ (blue) and *Hbs1l*^-/-^ (red) cells. Downward direction of arrow indicates significant lower pause scores of *Hbs1l*^-/-^- compared to *Pelo*^-/-^-specific pauses. (E) Examples of mapped footprints (27-34 nucleotides) on genes from tamoxifen treated control (CAG-Cre^ER^, grey) and *Pelo*^-/-^ (blue), or control and *Hbs1l*^-/-^ (red) cells. Upward direction of arrow indicates significant increase in local ribosome occupancy and the dashed line indicates the pause site. 4-OHT, 4-hydroxytamoxifin; nts, nucleotides; rpm, reads per million. Wilcoxon rank-sum test was used to determine statistical significance (D).

To analyze defects in translation elongation, we performed ribosome profiling of tamoxifen-treated control (CAG-Cre^ER^), *Pelo^-/-^* (CAG-Cre^ER^; *Pelo*^cKO^), and *Hbs1l^-/-^* (CAG-Cre^ER^; *Hbs1l*^cKO^) cells. Analyzing the ribosome occupancy in *Pelo*^-/-^ and control cells revealed ∼10,000 sites with significant (z-score ≥ 10) increases in local ribosome occupancy which mapped to 4,693 genes (*Figure 4C*, *Supplementary file 2*). One percent of these ribosome pauses were only observed in control cells (‘control-pauses’), 57% were shared between control and *Pelo^-/-^* cells, and 42% were specific to *Pelo*^-/-^ cells (‘*Pelo*^-/-^-pauses’). In contrast, analysis of *Hbs1l*^-/-^ and control cells revealed fewer ribosome pauses (∼4,200 - mapping to 1,807 genes) and only 5% were specific to *Hbs1l*^-/-^ cells (‘*Hbs1l*^-/-^-pauses’) (*Figure 4C*). In addition to the fewer ribosome pause sites, the z-scores (‘pause scores’) for *Hbs1l^-/-^*-pauses were also significantly lower than those of *Pelo*^-/-^-pauses (*Figure 4D*). Approximately 80% of the *Hbs1l^-/-^*-pausing genes also had pauses in *Pelo*^-/-^ cells, however only 30% of the ribosome pauses occurred at the same pause site (*Figure 4E*, *Supplementary file 2*). Together, these data suggest that the loss of *Pelo* or *Hbs1l* leads to translation elongation defects, but particularly severe defects are observed in the absence of *Pelo*.

The Dom34:Hbs1 complex has been implicated in multiple translation-dependent quality control pathways including non-stop decay (NSD) (Collart and Weiss, 2019; Simms et al., 2017). This pathway rescues ribosomes stalled at the ends of truncated mRNAs, ribosomes in polyA-sequences on prematurely polyadenylated mRNAs that lack a termination codon, or ribosomes in the 3’UTR of mRNAs that were not recycled at canonical stop codons (Arribere and Fire, 2018; D’Orazio et al., 2019; Guydosh and Green, 2014; Mills et al., 2017; Young et al., 2015). Examination of *Pelo*^-/-^- and *Hbs1l*^-/-^-pauses revealed that ∼92% of pauses mapped to the protein-coding region of transcripts (*Supplementary file 5*). The remaining local increases in ribosome occupancy were similarly split between the 5’ and 3’UTRs. Although the frequency of pauses in the 3’UTR was similar for mutant specific, control specific and shared pauses, the frequency of control-specific pauses that mapped to the 5’UTR was increased and those in the coding region were decreased (*Figure 4 - figure supplement 1D*). Thus, consistent with our observation in *Hbs1l*^-/-^ embryos, these data suggest that neither loss of *Hbs1l* nor *Pelo* in MEFs led to enrichment of ribosomes in the 5’ or 3’UTR.

We also searched for RPFs containing untemplated stretches of adenosines (A) at the 3’end indicative of ribosomes extending into the poly(A) tail of premature polyadenylated mRNAs. Ribosomes did not protect more then 15 consecutive A’s (*Figure 4 - figure supplement 1E*) suggesting that like in yeast, the poly(A) tract does not extend beyond the P-site of the ribosome (Guydosh and Green, 2014). Interestingly, we observed a significantly higher fraction of these 3’end A reads in *Pelo*^-/-^ compared to control cells (*Figure 4 - figure supplement 1F*), supporting a role for *Pelo* in rescuing ribosomes in polyA tails (D’Orazio et al., 2019; Guydosh and Green, 2017, 2014). However, the fraction of those reads did not significantly increase in *Hbs1l*^-/-^ MEFs (*Figure 4 - figure supplement 1F*), *Hbs1l*^-/-^ embryos (Student’s t-test, *P* = 0.4621) or *Hbs1l*^-/-^ cerebella (Student’s t-test*, P* = 0.3269).

In addition to NSD, Dom34:Hbs1 is implicated in ribosome rescue during No-Go decay (NGD), a quality control pathway in which ribosome stalling is evoked due to stretches of rare codons, secondary mRNA structures, amino acid starvation, or tRNA deficiency. NGD may trigger endonucleolytic cleavage of mRNAs due to ribosome collision, leading to 5’RNA intermediates that lack a stop codon and are then targeted by Dom34:Hbs1 via NSD (D’Orazio et al., 2019; Doma and Parker, 2006; Glover et al., 2020). Studies in Drosophila and C. elegans revealed that RNA intermediates that converge onto NSG may also be generated through additional mechanisms including RNA interference (RNAi) and nonsense-mediated decay (NMD) (Arribere and Fire, 2018; Hashimoto et al., 2017).

To determine if loss of *Pelo* or *Hbs1l* alters the frequency of ribosome pauses that are associated with potential targets such as NMD targets, we mapped pauses to unique coding transcripts, which is generally difficult due to the short nature of RPFs. Indeed, only a fraction (∼30%) of *Pelo*^-/-^ and *Hbs1l^-/-^*-pauses could be assigned to unique coding transcripts (1,518 and 95, respectively) (*Supplementary file 5*). Of these transcripts, ∼8% were classified as NMD transcripts and ∼92% were protein-coding transcripts. A similar percentage of pauses mapping to NMD transcripts (∼6.5%) was observed when we analyzed transcripts, which contained ribosome pauses shared between either control and *Pelo*^-/-^ cells, control and *Hbs1l*^-/-^ cells, or control-specific pausing transcripts (*Figure 4 - figure supplement 1G*). In addition, transcriptome analysis of MEFs revealed that NMD transcripts represented ∼6% of expressed transcripts (*Supplementary file 7*). These data suggest that loss of *Pelo* or *Hbs1l* did not lead to enrichment of ribosome pauses on transcripts predicted to undergo NMD.

Our observation that ribosome pausing is more dramatic in *Pelo*^-/-^ than *Hbs1l*^-/-^ cells correlates with the earlier lethality of *Pelo*^-/-^ embryos. To get a broader perspective of whether the loss of *Pelo* impacts other aspects of translation more than the *Hbs1l* deficiency, we analyzed at first the translational efficiency (TE, *Supplementary file 6*) of genes by normalizing the abundance of ribosomal footprint reads to that of the RNA sequencing reads. In *Pelo*^-/-^ cells about 35% (4,884) of genes displayed significant (adj. *P* ≤ 0.05) alterations in TE compared to control cells. The TE of 57% of these genes was increased and for 43% it was decreased. In contrast, the TE of only 4% of genes (314, 60% up- and 40% downregulated) was significantly changed in *Hbs1l*^-/-^ cells and genes with differential translational efficiency only showed a moderate correlation between *Hbs1l*^-/-^ and *Pelo*^-/-^ cells (Pearson’s correlation*, r* = 0.3739). The TE of *Pelo*^-/-^-specific pausing genes increased, but this effect was less for *Hbs1l^-/-^*-specific pausing genes (*Figure 5 - figure supplement 1A*), suggesting that elongation defects maybe more dramatic in *Pelo*^-/-^ cells as a result of the increase in translation of these genes.

In addition to its greater impact on the TE of genes, the loss of *Pelo* also had a stronger impact on gene translation in general as evidenced by the differential expression of ribosome-protected fragments (DE RPF, *Supplementary file 3*) (*Figure 5 - figure supplement 1B*). While *Hbs1l* deficiency altered translation of 26% (3,445) of genes, the loss of *Pelo* affected translation of 34% (4,965) of genes and led to significantly greater fold changes in gene expression in *Pelo*^-/-^ cells (Wilcoxon test, *P* < 2.2e-16). However, differentially translated genes between *Pelo*^-/-^ (DE RPF *Pelo*^-/-^) and *Hbs1l^-/-^* (DE RPF *Hbs1l*^-/-^) cells were strongly correlated (*Figure 5 - figure supplement 1C*), indicating that, although loss of *Hbs1l* may lead to smaller alterations in translation, most of the changes occur in the same direction.

In addition, 20% (1,060) and 2% (70) of the differentially translated genes in *Pelo*^-/-^ and *Hbs1l^-/-^* cells also contained ribosome pauses specifically found in *Pelo*^-/-^ and *Hbs1l^-/-^* cells, respectively, and translation of these pausing genes was both increased and decreased (*Figure 5 - figure supplement 1B*). However, changes in expression were under 2-fold for the majority of *Pelo*^-/-^- and *Hbs1l^-/-^-*pausing genes (88% of *Pelo*^-/-^- and 93% of *Hbs1l*^-/-^-pausing genes) and these changes were significantly lower compared to those in genes without mutant-specific pauses (*Figure 5 - figure supplement 1B and D*). Thus, similar to *Hbs1l*^-/-^ embryos, these data suggest that most of the translational changes in gene expression are not due to an increase in the ribosome occupancy but are likely a response to changes in mRNA expression and/or translation regulation.

Interestingly, we observed an opposing relationship between transcriptional expression changes (DE mRNA) and changes in translational efficiency (TE) of many genes in *Pelo*^-/-^ and *Hbs1l^-/-^* cells (*Figure 5A*). KEGG pathway analysis of genes with this opposing behavior revealed enrichment of several pathways (ribosome, ribosome biogenesis, RNA transport, spliceosome, cell cycle and lysosome) that overlapped with those of differently translated genes (DE RPF) (*Figure 5B, Figure 5 - figure supplement 1E*), suggesting that expression of genes in these pathways is translationally regulated, perhaps to restore homeostasis between the transcriptome and translatome.

**Figure 5.**
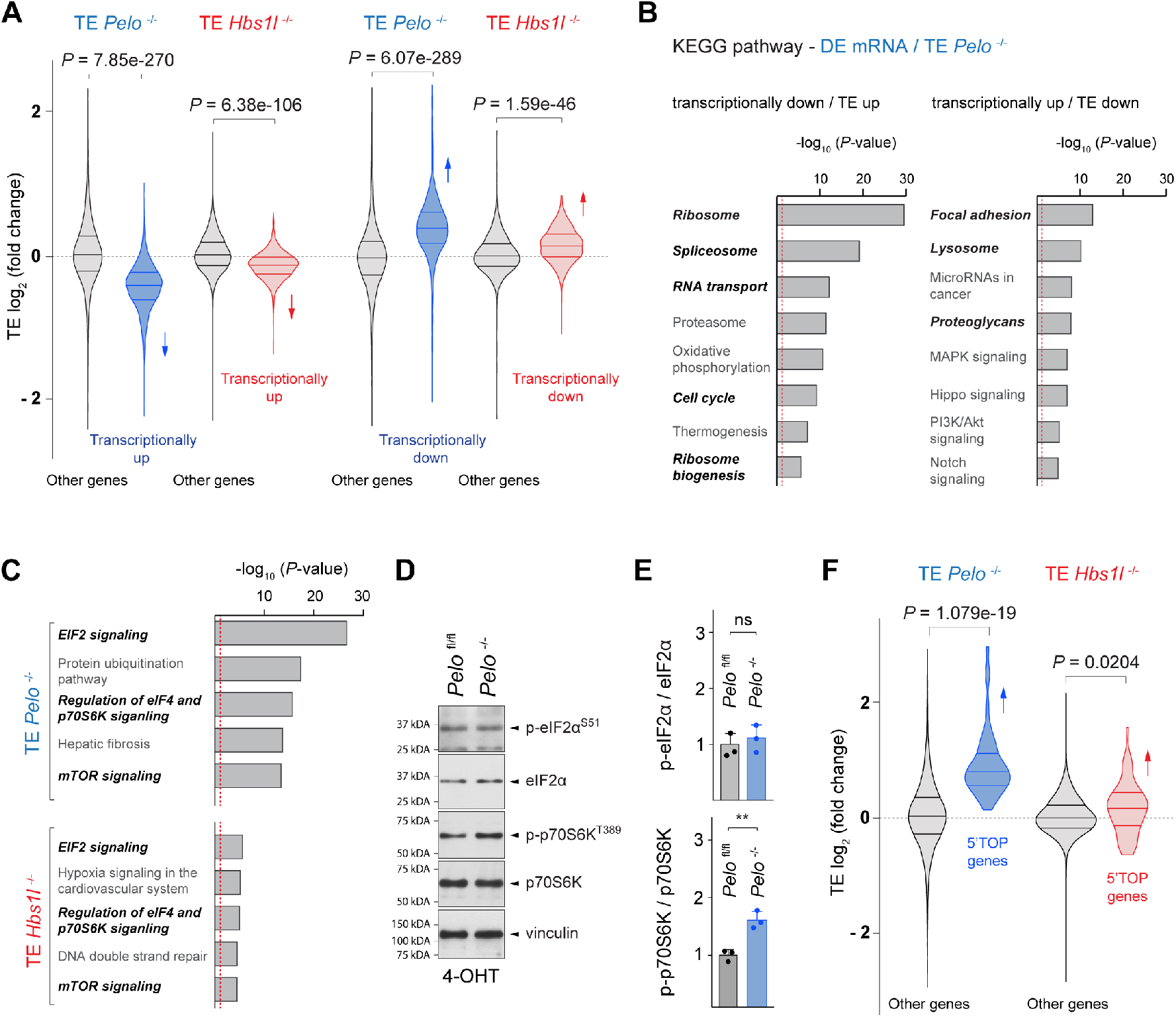
*Pelo*/*Hbs1l* deficiency alters translation regulation and reprograms the translatome. (A) Translational efficiency (TE) of genes that are transcriptionally upregulated or downregulated was compared to the remaining (‘other’) genes from *Pelo*^-/-^ (blue) and *Hbs1l*^-/-^ (red) MEFs, respectively. Downward direction of arrows indicates significant decrease in translational efficiency of transcriptionally upregulated genes in *Pelo*^-/-^ and *Hbs1l*^-/-^ MEFs. Upward direction of arrows indicates significant increase in translational efficiency of transcriptionally downregulated genes in *Pelo*^-/-^ and *Hbs1l*^-/-^ MEFs. (B) KEGG pathway analysis of genes from *Pelo*^-/-^ MEFs in which the translational efficiency (TE *Pelo*^-/-^, adj. *P* ≤ 0.05) is opposite to their transcriptional expression (DE mRNA *Pelo*^-/-^, *q*-value ≤ 0.05). Italicized pathways indicate pathways that overlapped with enriched pathways of differentially translated genes in *Pelo*^-/-^ MEFs (DE RPF *Pelo*^-/-^, adj. *P* ≤ 0.05) (Figure S6E). The red dashed line indicates the significance threshold (*P* = 0.05). (C) Ingenuity pathway analysis (IPA) of genes with differential translational efficiency from *Pelo*^-/-^ (TE *Pelo*^-/-^) and *Hbs1l*^-/-^ (TE *Hbs1l*^-/-^) MEFs. EIF2 and mTOR/p70S6K signaling are in italics. The red dashed line indicates the significance threshold (*P* = 0.05). (D) Western blot analysis of p-eIF2*α*^S51^ and p-p70S6K^T389^ using lysates of tamoxifen-treated control (*Pelo*^fl/fl^) and *Pelo*^-/-^ MEFs at Day 2 (Passage P1). Vinculin was used as an input control. (E) Levels of p-eIF2*α*^S51^ or p-p70S6K^T389^ were normalized to total level of eIF2*α* or p70S6K, and phosphorylation levels are relative to those of control (*Pelo*^fl/fl^). Data represent mean + SD. (F) Translational efficiency (TE) of genes with translational regulation by mTOR via their 5’TOP motif was compared to the remaining (‘other’) genes from *Pelo*^-/-^ (blue) and *Hbs1l*^-/-^ (red) MEFs. Upward direction of arrows indicates significant increase in translational efficiency of 5’TOP genes. 4-OHT, 4-hydroxytamoxifin; 5’TOP, 5’terminal oligopyrimidine motif. Student’s t-test (E); Wilcoxon rank-sum test was used to determine statistical significance (A, F). ns, not significant; ** *P* ≤ 0.01.

To identify signaling pathways that might control these changes in translation regulation upon loss of *Pelo*/*Hbs1l*, we performed IPA analysis on differentially transcribed (DE mRNA) genes and genes with altered translation efficiency (TE). EIF2 and mTOR/p70S6K signaling, both of which are known to regulate translation, were highly enriched in *Pelo*^-/-^ cells but less enriched in *Hbs1l*^-/-^ cells (*Figure 5C, Figure 5 - figure supplement 1F*). Phosphorylation of eIF2*α* decreases translation initiation, while the activity of mTOR, in particular mTORC1 (mechanistic target of rapamycin complex 1), increases translation initiation and elongation. About 50% of the differentially regulated genes identified by IPA overlapped between the two pathways and therefore, we assessed the phosphorylation status of p-eIF2*α*^S51^ and p-p70S6^T389^, a known target of mTORC1, to determine if one or both signaling pathways were affected. Levels of p-eIF2*α*^S51^ in *Pelo*^-/-^ cells were unchanged from those of control cells (*Figure 5D and E*). However, levels of p-p70S6^T389^ were significantly increased in *Pelo*^-/-^ cells, indicating activation of mTOR signaling (*Figure 5D and E*). In agreement, the TE of genes that are known to be translationally regulated by mTORC1 via their 5’terminal oligopyrimidine motifs (5’TOP) (Yamashita et al., 2008) was significantly increased in *Pelo*^-/-^ and *Hbs1l*^-/-^ cells, although this increase was less pronounced in the latter (*Figure 5F*). Activation of mTORC1 may underlie some of the observed gene expression changes (DE RPF) given its role as a positive regulator for ribosome biogenesis and translation of ribosomal genes and a negative regulator of lysosomal biogenesis and autophagy (Kim et al., 2011; Mayer and Grummt, 2006; Puertollano, 2014; Rabanal-Ruiz and Korolchuk, 2018; Roczniak- Ferguson et al., 2012), which parallels the directionality of the changes in these pathways (*Figure 5 - figure supplement 1E*).

### Convergent modulation of the translatome in cells with defects in translation-dependent quality control pathways

Our findings suggest that translational reprogramming occurs upon loss of *Pelo/Hbs1l*. However, whether these changes are unique to *Pelo/Hbs1l* or reflect a more general cellular response upon impairment of translation-dependent quality control pathways is unclear. To investigate this possibility, we conditionally deleted the core NMD component, *Upf2*, in MEFs (Lelivelt and Culbertson, 1999; Serin et al., 2001). In contrast to NGD and NSD that resolve ribosomes on mRNAs impeding translation elongation, nonsense-mediated decay (NMD) targets aberrant mRNAs (e.g. mRNAs containing a premature termination codon) for degradation during translation (Karousis and Mühlemann, 2019; Schuller and Green, 2019). Consistent with the function of *Upf2* in NMD, about 20% and 13% of upregulated transcripts (DE mRNA *Upf2*, *q-value* ≤ 0.05) were NMD transcripts or transcripts with retained introns, respectively (*Figure 6 - figure supplement 1A*, *Supplementary file 7*). Although the accumulation of these transcripts was specific to *Upf2* ^-/-^ cells (*Figure 6 - figure supplement 1A and B*), we observed an inverse relationship between transcriptional gene expression changes (DE mRNA *Upf2*) and changes in translational efficiency (TE *Upf2*) in *Upf2^-/-^* cells as we did in *Pelo*^-/-^ and *Hbs1l^-/-^* cells (*Figure 6A*). Surprisingly, IPA analysis on differentially transcribed (DE mRNA *Upf2*^-/-^) genes and genes with altered translation efficiency (TE *Upf2*^-/-^) revealed enrichment for EIF2 and mTOR signaling in *Upf2^-/-^* as observed in *Pelo*^-/-^ and *Hbs1l^-/-^* cells (*Figure 6B, Figure 6 - figure supplement 1C*). Western blot analysis revealed the level of p-eIF2*α*^S51^ in *Upf2^-/-^* cells was unchanged from that of controls, but p-p70S6^T389^ levels and translation of 5’TOP genes were significantly increased (*Figure 6C, D and E*), indicating that mTORC1 is activated in *Upf2^-/-^* cells. Although translation of ribosomal genes was increased, transcriptional levels of ribosomal genes were decreased in *Upf2*^-/-^ similar to *Pelo*^-/-^ cells (*Figure 6F*), further supporting that mTORC1 activation may be a general response in an attempt to restore cellular homeostasis in *Upf2*^-/-^ and *Pelo*^-/-^ cells.

**Figure 6.**
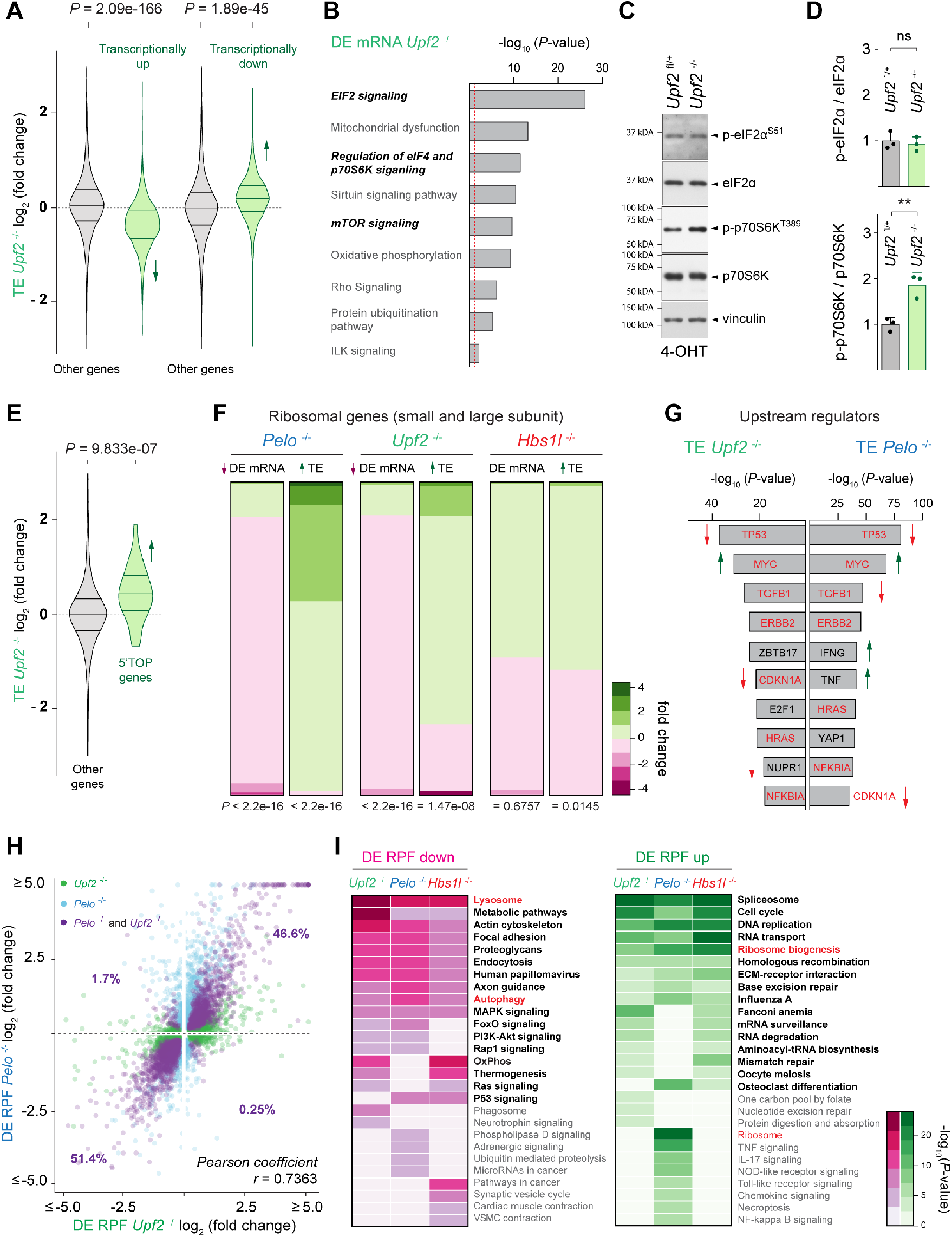
Defects in different translation-dependent quality control pathways similarly alter translation regulation and the translatome. (A) Translational efficiency (TE) of genes that are transcriptionally upregulated or downregulated was compared to the remaining (‘other’) genes from *Upf2*^-/-^ (green) MEFs. Downward direction of arrows indicates significant decrease in translational efficiency of transcriptionally upregulated genes in *Upf2*^-/-^ MEFs. Upward direction of arrows indicates significant increase in translational efficiency of transcriptionally downregulated genes in *Upf2*^-/-^ MEFs. (B) Ingenuity pathway analysis (IPA) of differentially transcribed genes from *Upf2*^-/-^ MEFs (DE mRNA *Upf2*^-/-^). EIF2 and mTOR/p70S6K signaling are in italics. The red dashed line indicates the significance threshold (*P* = 0.05). (C) Western blot analysis of p-eIF2*α*^S51^ and p-p70S6K^T389^ of lysates from tamoxifen-treated control (*Upf2*^fl/+^) and *Upf2*^-/-^ MEFs at Day 2 (Passage P1). Vinculin was used as an input control. (F) Levels of p- eIF2*α*^S51^ or p-p70S6K^T389^ were normalized to total level of eIF2*α* or p70S6K, and phosphorylation levels are relative to those of control (*Upf2*^fl/+^) MEFs. Data represent mean + SD. (E) Translational efficiency (TE) of genes those translation is regulated by mTOR via their 5’TOP motif was compared to the remaining (‘other’) genes from *Upf2*^-/-^ (green) MEFs. Upward direction of arrows indicates significant increase in translational efficiency of 5’TOP genes. (F) Differentially transcription (DE mRNA) or translational efficiency (TE) of ribosomal protein genes (small and large ribosomal subunit) was compared to the remaining (‘other’) genes from *Pelo*^-/-^ (blue), *Upf2*^-/-^ (green) and *Hbs1l*^-/-^ (red) MEFs. Up- and downward direction of arrow indicates significant up- and downregulation of ribosomal protein genes, respectively. The heatmap indicates the gene expression changes of ribosomal protein genes. (G) Identification of upstream regulators of genes with differential translational efficiency in *Upf2*^-/-^ (TE *Upf2*^-/-^) and *Pelo*^-/-^ (TE *Pelo*^-/-^) MEFs. Top ten transcription factors are shown. Those enriched in both *Upf2*^-/-^ and *Pelo*^-/-^ MEFs and shown in red. Up- or downward direction of arrow indicates predicated up- or downregulation of transcription factors, respectively. (H) Differentially translated genes in *Upf2*^-/-^ MEFs (DE RPF *Upf2*^-/-^, adj. *P* ≤ 0.05, x-axis, green) were plotted against genes that are differentially translated in *Pelo*^-/-^ MEFs (DE RPF *Pelo*^-/-^, adj. *P* ≤ 0.05, y-axis, blue). Genes those translation was significantly different in both *Upf2*^-/-^ and *Pelo*^-/-^ MEFs are shown in purple. (I) KEGG pathway analysis of differentially translated (up- and downregulated) genes (DE RPF, adj. *P* ≤ 0.05) in *Upf2*^-/-^ (green), *Pelo*^-/-^ (blue) and *Hbs1l*^-/-^ (red) MEFs. Significantly (*P* ≤ 0.05) enriched pathways are shown and pathways in bold indicate pathways that are shared between any of the mutant MEFs. Pathways known to be positively and negatively regulated by mTORC1 are in red. 4-OHT, 4- hydroxytamoxifin; 5’TOP, 5’terminal oligopyrimidine motif. Student’s t-test (D); Wilcoxon rank-sum test was used to determine statistical significance (A, E, F); Pearson coefficient (*r*) was determined to analyze linearity of gene expression changes (H). ns, not significant; ** *P* ≤ 0.01.

Surprisingly, most of the top upstream regulators including *TP53*, *MYC*, *TGFB1*, *ERBB2*, *CDKN1A*, *HRAS* and *NFKBIA* were shared between *Upf2*^-/-^ and *Pelo*^-/-^ cells (*Figure 6G*) and genes with differential translational efficiency were strongly correlated between *Upf2*^-/-^ and *Pelo*^-/-^ cells (Pearson’s correlation*, r* = 0.6576). Furthermore, differentially translated genes in *Upf2*^-/-^ (32% of genes, DE RPF *Upf2*^-/-^) and *Pelo*^-/-^ (34% of genes, DE RPF *Pelo*^-/-^) cells also showed a strong linear correlation (*Figure 6H*), indicating that defects in these quality control pathways may not only lead to similar changes in translation regulation (TE) but also in global gene translation (DE RPF). Consistent with the similar changes in translation, KEGG pathway analysis of differentially translated genes (DE RPF, adj. *P* ≤ 0.05) revealed similar enrichment of multiple pathways in *Upf2*^-/-^, *Pelo*^-/-^ and *Hbs1*^-/-^ cells (*Figure 6I*).

In contrast with the many ribosome pauses we observed specifically in *Pelo*^-/-^ cells, only 4% of all ribosome pausing events observed in *Upf2*^-/-^ and control cells were uniquely found in *Upf2*^-/-^ cells (*Figure 6 - figure supplement 1D*). Although, only a small fraction (8%) of *Pelo-* and *Hbs1l*- specific pausing transcripts corresponded to NMD transcripts, we considered the possibility that some of the protein-coding transcripts with ribosome pauses in *Pelo*^-/-^ and *Hbs1l*^-/-^ cells may also be NMD sensitive given that 5-10% of mRNAs even without premature termination codons are thought to be degraded by the NMD pathway (He et al., 2003; Jaffrey and Wilkinson, 2018; Lelivelt and Culbertson, 1999; Mendell et al., 2004). However, upregulation of these protein-coding pausing transcripts was not observed in *Upf2*^-/-^ cells (*Figure 6 - figure supplement 1E*), suggesting that these pausing transcripts are not NMD sensitive. Together, these findings suggest that while *Pelo/Hbs1l* and *Upf2* largely function in distinct quality control pathways, disruption of either pathway results in similar translational gene expression changes.

### Deletion of *Upf2* or *Pelo* cause similar cerebellar developmental defects

Intrigued by the similar changes in translation upon impairment of *Upf2* and *Pelo/Hbs1l* in MEFs, we conditionally deleted *Upf2* during cerebellar development to determine if phenotypic similarities exist upon loss of these different translation-dependent quality control pathways. Surprisingly, deletion of *Upf2* (En1-Cre; *Upf2*^cKO^) or *Pelo* (En1-Cre; *Pelo*^cKO^) using En1-Cre resulted in largely indistinguishable defects with a grossly hypoplastic cerebellum and regions of the midbrain (superior and inferior colliculus) being nearly absent unlike in En1-Cre; *Hbs1l*^cKO^ mice (*Figure 7A and 2D*). Both *Upf2* and *Pelo* mutant pups died shortly after birth, perhaps due to En1-Cre deletion of these genes in other cell types (Britz et al., 2015; Kimmel et al., 2000; Sapir et al., 2004; Sgaier et al., 2007; Wurst et al., 1994).

**Figure 7.**
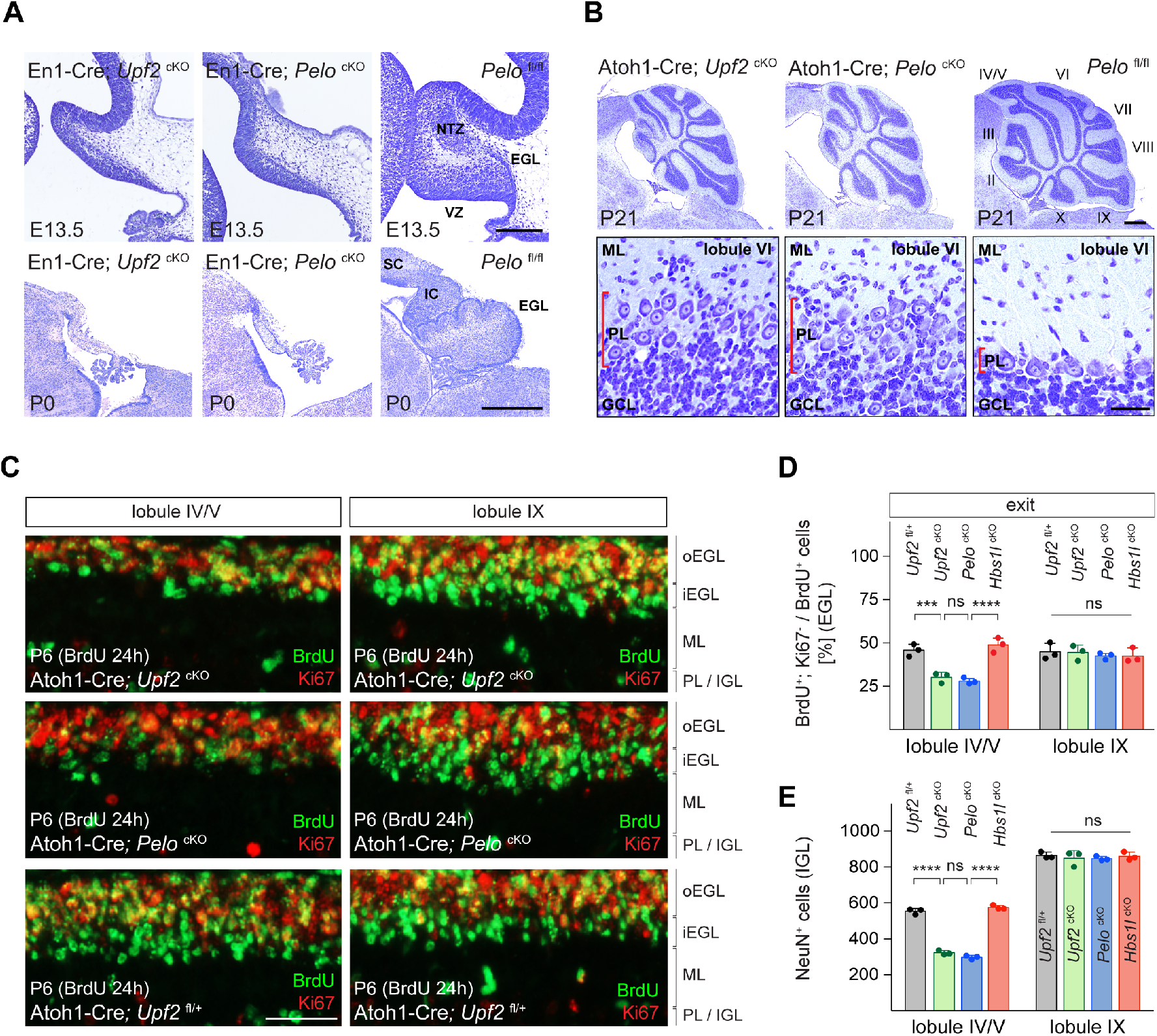
Loss of *Upf2* and *Pelo* cause similar cerebellar developmental defects. (A) Parasagittal (E13.5) and sagittal (P0) cerebellar sections of control (*Pelo*^fl/fl^), En1-Cre; *Upf2*^cKO^ and En1-Cre; *Pelo*^cKO^ mice stained with cresyl violet. (B) Sagittal cerebellar sections of P21 control (*Pelo*^fl/fl^), Atoh1-Cre; *Upf2*^cKO^ and Atoh1-Cre; *Pelo*^cKO^ mice stained with cresyl violet. Higher magnification images of lobule VI are shown below for each genotype. Cerebellar lobules are indicated by Roman numerals. (C) Immunofluorescence with antibodies to BrdU (green) and Ki67 (red) on sections of P6 control (Atoh1-Cre; *Upf2*^fl/+^), Atoh1-Cre; *Upf2*^cKO^ and Atoh1-Cre; *Pelo*^cKO^ cerebellum. Mice were injected with BrdU 24 hours prior to harvest to determine the fraction of granule cell precursors in the EGL that exited the cell cycle. Images are shown for anterior (IV/V) and posterior (IX) lobules. (D) Quantification of the fraction of granule cell precursors in the EGL (lobules IV/V and IX) that exited the cell cycle (BrdU^+^, Ki67^-^ cells) of control (Atoh1-Cre; *Upf2*^fl/+^), Atoh1-Cre; *Upf2*^cKO^, Atoh1-Cre; *Pelo*^cKO^ and Atoh1-Cre; *Hbs1l*^cKO^ mice. Data represent mean + SD. (E) Quantification of terminally differentiated granule cells (NeuN^+^ cells) in the IGL in lobules IV/V and IX of control (Atoh1-Cre; *Upf2*^fl/+^), Atoh1-Cre; *Upf2*^cKO^, Atoh1-Cre; *Pelo*^cKO^ and Atoh1-Cre; *Hbs1l*^cKO^ mice. Data represent mean + SD. Scale bars: 200μm (E13.5) and 500μm (P0) (A); 500μm and 20μm (higher magnification) (B); 50μm (C). VZ, ventricular zone, EGL, external granule cell layer; NTZ, nuclear transitory zone; SC, superior colliculus; IC, inferior colliculus; ML, molecular cell layer; PL, Purkinje cell layer; GCL, granule cell layer; oEGL, outer external granule cell layer; iEGL, inner external granule cell layer; IGL, internal granule cell layer. Two-way ANOVA was corrected for multiple comparisons using Tukey method (D, E). ns, not significant; *** *P* ≤ 0.001; **** *P* ≤ 0.0001.

Similar neurogenesis defects were observed in the *Upf2* and *Pelo* mutant cerebellum. The fraction of ventricular zone (VZ) progenitors that remained in the cell cycle was higher in E13.5 *Upf2*- or *Pelo*-deficient cerebella compared to controls (*Figure 7 - figure supplement 1A and B*), indicating that like *Hbs1l*, loss of *Upf2* or *Pelo* impairs cell cycle exit of VZ-progenitors. Inversely, the number of VZ-derived precursors, e.g. Purkinje cells (Lhx1/5^+^ cells) and Pax2^+^ interneurons were reduced by ∼83% and ∼90% in the En1-Cre; *Upf2*^cKO^ and En1-Cre; *Pelo*^cKO^ cerebellum (*Figure 7 - figure supplement 1C, D, E and F*). Glutamatergic cerebellar neurons (Tbr1^+^) were nearly absent in the E13.5 En1-Cre; *Upf2*^cKO^ and En1-Cre; *Pelo*^cKO^ cerebellum (*Figure 7 - figure supplement 1G and H*). In addition, the EGL was missing in En1-Cre; *Upf2*^cKO^ and En1-Cre; *Pelo*^cKO^ cerebella at both E13.5 and P0 (*Figure 7A*, *Figure 7 - figure supplement 1I*), indicating that neurogenesis defects are particularly more severe in the absence of *Upf2* or *Pelo* relative to those observed upon *Hbs1l* loss.

Thus, we considered that in addition to cell cycle exit abnormalities, cell death might also contribute to impaired cerebellar development. Indeed, the number of apoptotic cells (Casp3^+^ cells) was significantly increased in the En1-Cre; *Upf2*^cKO^ and En1-Cre; *Pelo*^cKO^ cerebellum compared to that of controls or En1-Cre; *Hbs1l*^cKO^ embryos, and apoptotic cells were observed in both the ventricular zone and the prospective white matter where progenitor-derived progeny reside (*Figure 7 - figure supplement 1J and K*). Together, these data suggest that defects in cell cycle exit and increased cell death likely impair cerebellar development in *Upf2* and *Pelo* mutant mice. To determine if loss of *Upf2* or *Pelo* also increased mTORC1 signaling in the developing cerebellum as it did in MEFs, we analyzed levels p-S6^S240/244^, a known downstream target of mTORC1. In agreement, levels of p-S6^S240/244^ were significantly increased in the En1-Cre; *Upf2*^cKO^ and En1-Cre; *Pelo*^cKO^ cerebellum, including the VZ (*Figure 7 - figure supplement 1L and M*).

Because loss of *Upf2* or *Pelo* severely impaired early granule cell neurogenesis, we investigated if these genes are also required later in the development of these cells. Conditional deletion of either *Upf2* or *Pelo* using Atoh1*-*Cre resulted in abnormalities of the anterior lobes of the cerebellum in P21 mice (*Figure 7B*). These lobes were reduced in length and Purkinje cells failed to form a monolayer. In contrast, the posterior lobes appeared unaffected, possibly due to the anterior-to-posterior gradient of *Cre* expression in granule cell precursors in the developing cerebellum (Pan et al., 2009; Qiu et al., 2010; Wojcinski et al., 2019).

During postnatal cerebellar development, granule cell precursors in the outer region of the EGL (oEGL) exit the cell cycle and transiently reside in the inner EGL (iEGL) before migrating to the internal granule cell layer (IGL). To determine if granule cell precursors in postnatal Atoh1-Cre; *Upf2*^cKO^ or Atoh1-Cre; *Pelo*^cKO^ cerebellum properly exited the cell cycle, we labeled granule cell precursors with BrdU to determine the fraction of precursors that exited the cell cycle twenty-four hours later (BrdU^+^; Ki67^-^ cells). In P6 control cerebella, BrdU-labeled granule cell precursors that were negative for Ki67 formed a distinct layer on the ventral surface of the EGL consistent with the appearance and location of the iEGL (Legué et al., 2016) (*Figure 7C*). However, no clear separation between the oEGL and iEGL was observed in the anterior lobes in Atoh1-Cre; *Upf2*^cKO^ or Atoh1-Cre; *Pelo*^cKO^ cerebella (*Figure 7C*), and the fraction of granule cell precursors that exited the cell cycle (BrdU^+^; Ki67^-^ cells) in these lobes, but not posterior lobes, was significantly lower compared to control or Atoh1-Cre; *Hbs1l*^cKO^ cerebella (*Figure 7D*). Correspondingly, the number of terminally differentiated granule cells in the IGL (NeuN^+^ cells) of the anterior, but not posterior, lobes was reduced by ∼45% in P6 Atoh1-Cre; *Upf2*^cKO^ and Atoh1-Cre; *Pelo*^cKO^ cerebella (*Figure 7E*), indicating that both of these genes are necessary for differentiation of granule cell precursors.

## Discussion

Translation-dependent quality control pathways govern protein synthesis and proteostasis by degrading aberrant mRNAs that result in potentially toxic peptide products. However, little is known about the *in vivo* defects that arise in the absence of these pathways and how translation is altered in the absence of these pathways. To interrogate these questions, we investigated the translational and phenotypic alterations from different translation-dependent quality control pathways including the NSG/NGD (*Pelo*/*Hbs1l*) and the NMD (*Upf2*) pathways.

Here we show that *Pelo*/*Hbs1l* are both critical for embryonic and cerebellar development but dispensable after cerebellar development of granule cells in mice. In addition, loss of *Pelo* or *Hbs1l* locally increases the ribosome occupancy (‘ribosome pauses’) of genes. Interestingly, the number and strength of these pauses were higher in E8.5 *Hbs1l*^-/-^ embryos, just prior to the time when these embryos cease developing, compared to the postnatal cerebellum, where loss of *Hbs1l* had no effect on cerebellar granule cells. Together these data suggest that this translation-dependent quality control pathway may be needed in cell- and/or tissue-specific manner.

In agreement with the differential severity between *Pelo*^-/-^*-* and *Hbs1l*^-/-^-mediated defects during embryonic and cerebellar development, pauses were higher and more frequent in *Pelo*^-/-^ MEFs and coincided with greater changes in gene expression than in *Hbs1l*^-/-^ MEFs. Loss of *Pelo* induced greater translation elongation defects compared to its binding partner *Hbs1l*, which may be in part be influenced by differences in gene expression and translational efficiency between *Pelo* and *Hbs1l* deficient cells. However, additional factors may also play a role. For example, deletion of either *Pelo* or *Hbs1l* led to a reduction of their respective binding partners, the kinetics of this decrease upon deletion of *Pelo* or *Hbs1l* is unknown and may not be equal. In addition, Dom34 (*Pelo*) promotes dissociation of ribosomes by Rli1 (*Abce1*) and this activity increased in the presence of Hbs1 (*Hbs1l*) (Pisareva et al., 2011; Shoemaker and Green, 2011). Hence, we cannot rule out a scenario in which the remaining levels of PELO in *Hbs1l*^-/-^ cells might be able to mitigate and/or delay elongation defects even in the absence of *Hbs1l*.

Most of the ribosome pauses in *Pelo*- and *Hbs1l-*deficient tissues or cells were located in the coding sequence. Although these pauses with a footprint length 27-34 nucleotides could indicate ribosomes that paused during translation, phenotypes in *Hbs1l* and *Pelo* mutant mice were not dependent on the B6J-associated mutation in *n-Tr20* that introduces genome-wide AGA pausing within mRNAs. Instead, these pauses may reflect pausing of the ‘trailing’ ribosomes upstream of the ‘lead’ ribosome that reached the 3’end of truncated mRNAs (Guydosh et al., 2017; Guydosh and Green, 2014). Footprints of the ‘lead’ ribosome have been analyzed on an exosome-deficient background (Arribere and Fire, 2018; D’Orazio et al., 2019; Glover et al., 2020; Guydosh and Green, 2014), and its short length of 15-18 nucleotides makes it difficult to uniquely map these footprints in mammals with their larger genomes. Regardless, recent biochemical studies have demonstrated that the *Pelo*/*Hbs1l* complex rescues trapped ribosomes near the 3’end of truncated mRNAs but is not necessary for the resolution of internally stalled ribosomes within the mRNA (Juszkiewicz et al., 2020b), which is consistent with early biochemical and structural studies demonstrating that Dom34:Hbs1 preferentially rescues ribosomes arrested at sites of truncation (Hilal et al., 2016; Pisareva et al., 2011; Shoemaker and Green, 2011).

RNA intermediates that converge on *Pelo*/*Hbs1l* may derive from endonucleolytic cleavage for example during NMD, RNAi or NGD (Arribere and Fire, 2018; D’Orazio et al., 2019; Doma and Parker, 2006; Eberle et al., 2009; Hashimoto et al., 2017). However, most of the *Pelo*^-/-^- and *Hbs1l*^-/-^-pausing transcripts were not NMD sensitive, suggesting that if pausing occurred on truncated mRNAs, many of the RNA intermediates likely derived from additional mechanisms.

Endonucleolytic cleavage of mRNAs may occur upon persistent ribosome collision during NGD (D’Orazio et al., 2019; Doma and Parker, 2006; Shoemaker and Green, 2012). Intriguingly, widespread ribosome collision has been observed under normal conditions and may present 10% of the pool of translating ribosomes (Arpat et al., 2020; Diament et al., 2018; Han et al., 2020; Meydan and Guydosh, 2020). Numerous human mRNAs are subject to repeated, co-translational endonucleolytic cleavages and this process is similar to NGD, but independent of NMD-associated nucleases (Ibrahim et al., 2018). Because translation changes during development and varies between cell types (Blair et al., 2017; Buszczak et al., 2014; Castelo-Szekely et al., 2017; Gonzalez et al., 2014; Sudmant et al., 2018), RNA intermediates that are generated during translation may also vary and thereby, could introduce a need for quality control pathways in a tissue- and/or cell type-specific manner.

We also observed that loss of *Pelo/Hbs1l* was associated with the activation of mTORC1 signaling. mTORC1 activation was previously observed in *Hbs1l* patient derived fibroblasts (O’Connell et al., 2019) and upon deletion of *Pelo* during epidermal development (Liakath-Ali et al., 2018). Interestingly, inhibition of translation/mTORC1 partially prevented *Pelo*^-/-^-mediated epidermal defects (Liakath-Ali et al., 2018). How mTORC1 responds to loss of *Pelo*/*Hbs1l* is unknown. Multiple mTOR-dependent phosphorylation sites on the surface of the ribosome have been observed, suggesting that mTORC1 and/or mTORC1-associated kinases interact with the ribosomes and might provide a mechanism to detect changes in translation elongation (Jiang et al., 2016). However, changes in levels of ribosomal genes and/or impaired ribosomal biogenesis also activate mTORC1 signaling (Liu et al., 2014). Indeed, deletion of *Pelo/Hbs1l* led to decreased transcript levels but increased translation of ribosomal genes. Thus, mTORC1 may be activated to compensate for decreases in expression of these genes rather than directly sensing defects in translation elongation. To test whether mTORC1 activation was specific to *Pelo*/*Hbs1l* deficiency or was generally associated with defects in translation-dependent quality control pathways, we examined MEFs deficient for *Upf2*, an essential component of the NMD pathway. Loss of *Upf2* also led to decreased expression and increased translation of ribosomal genes and mTORC1 activation, supporting that changes in mTORC1 signaling are not a direct consequence of the elongation defects, but likely occurs as a compensatory response.

In general, impairment of either translation-dependent quality control pathway led to strikingly similar alterations in gene expression with *Pelo* and *Upf2* showing the most changes compared to *Hbs1l*. Reminiscent of these changes, loss of *Pelo* or *Upf2* had remarkably similar effects on multiple cerebellar neuronal populations during early embryonic and late postnatal neurogenesis, causing comparable morphological defects in the cerebellum (e.g. inhibition of differentiation and increase in cell death). Growing evidence highlights the role of mTOR in the decision of stem cells to self-renew or differentiate (Meng et al., 2018; Xiang et al., 2011). However, increased mTOR activity generally reduces self-renewal and promotes differentiation of neuronal stem/progenitor cells (Hartman et al., 2013; Licausi and Hartman, 2018; Magri et al., 2011; Way et al., 2009), suggesting that likely other molecular changes contributed to the observed defects in neurogenesis.

The similarities in gene expression and developmental alterations in mice deficient for these translation-dependent quality control pathways suggest a convergence of molecular and cellular pathology. In fact, several changes in gene expression and signaling pathways were altered in the same direction in mutant cells. Thus, phenotypic changes could be due to either a change in a single molecular pathway or interactions of multiple pathways. For example, *Myc* activation was predicted as an upstream regulator of gene expression in both *Pelo*^-/-^ and *Upf2*^-/-^ but less in *Hbs1l*^-/-^ MEFs. *Myc* functions as a switch between proliferation and differentiation during cerebellar development (Knoepfler et al., 2002; Ma et al., 2015; Wey et al., 2010), in which loss of *Myc* allows precursors to exit the cell cycle and to undergo differentiation, while maintained *Myc* expression retains cells in the proliferation cycle. Intriguingly, previous studies revealed that depletion of NMD factors inhibited differentiation of embryonic stem cells, which coincided with sustained *Myc* expression and its downregulation released the differentiation blockage in NMD deficient cells (Li et al., 2015).

How different translation-dependent quality control pathways can lead to similar changes in translation and cellular defects is unclear. Recent studies in yeast demonstrated that impairment of these quality control mechanisms (*Hbs1*, *Dom34*, *Upf1*, *Upf2*, *Ski7* and *Ski8*) caused protein aggregation (Jamar et al., 2018). Protein misfolding occurred co-translationally on highly translated genes and the aggregated proteins overlapped between the different mutant strains. These data suggest that increased translation and protein aggregation may be common properties among different quality control mutants (Jamar et al., 2018). Perhaps, regardless of specific targets of various quality control pathways, defects in protein folding, clearance of defective peptide products, and/or defects in mRNA decay may trigger similar cellular responses leading to similar phenotypes.

## Material and Methods

### Mouse strains

Generation of *Hbs1l*^GTC^ mice was performed by injection of targeted ES cells (International Gene Trap Consortium, IGTC, cell line ID: XE494) into C57BL/6J (B6J) blastocysts. B6N-*Hbs1l*^tm1a^ (C57BL/6N-*A^tm1Brd^Hbs1l^tm1a(KOMP)Wtsi^*, MMRRC:048037) mice were produced at the Wellcome Trust Sanger Institute Mouse Genetics Project as part of the International Mouse Phenotype Consortium (IMPC). In order to generate the conditional *Hbs1l* knock out allele, heterozygous B6N-*Hbs1l*^tm1a/+^ mice were crossed to B6N.129S4-Gt(ROSA)26Sor^tm1(FLP1)Dym^/J (The Jackson Laboratory, #016226, MGI:5425632) to remove the gene trap cassette and generate B6N-*Hbs1l*^fl/+^ mice. Generation of the constitutive B6N-*Hbs1l*^+/-^ knock out allele was accomplished by crossing homozygous B6N-*Hbs1l*^fl/fl^ mice to B6N.Cg-Edil3Tg(Sox2-Cre)1Amc/J mice (The Jackson Laboratory, #014094, MGI:4943744). The B6N-*Hbs1l*^+/-^ or B6N-*Hbs1l*^fl/fl^ mice were backcrossed to congenic B6N.B6J^n-Tr20^ mice (n = 2 backcross generations) to introduce the B6J-associated *n-Tr20* mutation. The conditional knock out *Pelo* allele was generated by placing the 5’loxP site 117bp upstream of exon 2 and the 3’loxP site 302bp downstream of exon2. Targeted B6J ES cells were injected into B6J blastocysts to generate heterozygous B6J-*Pelo*^fl/+^. Generation of the ubiquitous B6J-*Pelo*^+/-^ knock out allele was accomplished by crossing homozygous B6J-*Pelo*^fl/fl^ mice to B6.FVB-Tg(EIIa-cre)C5379Lmgd/J mice (The Jackson Laboratory, #003724, MGI:2174520). The conditional *Upf2* knock out allele (*Upf2*^fl/fl^ mice) was kindly provided from Drs. Bo Torben Porse and Miles Wilkinson. Genotyping primers are listed below and for the conditional knock out alleles of *Hbs1l* and *Pelo,* genotyping primers were multiplexed to simultaneously detect the wild type, flox (fl) and delta allele. Genotyping for the conditional knock out allele of *Upf2*, was performed as previously described (Weischenfeldt et al., 2008).

*Hbs1l common* Forward: 5’AGTCCAGGTGTTTCCTCACG’3
*Hbs1l wild type* Reverse: 5’CCCTGGCCTATTTTTGGTTT’3
*Hbs1l GTC* Reverse: 5’TGTCCTCCAGTCTCCTCCAC’3

*Hbs1l cKO* Forward I: 5’CATGGCCTCCTATGGGTTGA’3
*Hbs1l cKO* Forward II: 5’GCCTACAGTGAGCACAGAGT’3
*Hbs1l cKO* Reverse: 5’TAGGTGCTGGGATTTGAACC’3

*Pelo cKO* Forward: 5’TGTAACTGAACCCTGCAGTATCT’3
*Pelo cKO* Reverse I: 5’GTGGAGCATGAAATGAAATTCGG’3
*Pelo cKO* Reverse II: 5’ATCCAAGGCTTTTACTTCGCC’3

For conditional knock out experiments, the following *Cre*-lines were used and the *Cre* allele was maternally inherited to generate mutant mice (F_2_ generation): En1^tm2(cre)Wrst^/J (En1-Cre, The Jackson Laboratory, #007916, MGI:3815003), B6.Cg-Tg(Atoh1-cre)1Bfri/J (Atoh1-Cre, The Jackson Laboratory, #011104, MGI:4415810) and B6.Tg(Gabra6-cre)B1Lfr (Gabra6-Cre, MGI:4358481, Fünfschilling and Reichardt, 2002). In order to avoid the introduction of the B6J- associated *n-Tr20* mutation in En1-Cre; *Hbs1l*^cKO^ mice, En1^tm2(cre)Wrst^/J mice were backcrossed to congenic B6J.B6N^n-Tr20^ mice (Ishimura et al., 2016) to generate En1^tm2(cre)Wrst^/J mice that no longer carry the B6J-assciated mutation in *n-Tr20*. Subsequently, B6J.B6N^n-Tr20^; En1^cre^ mice were intercrossed with B6N-*Hbs1l*^+/-^ to produce F1 mice, then these mice were crossed to B6N-*Hbs1l*^fl/fl^ mice to generate En1-Cre; *Hbs1l*^cKO^; *n-Tr20*^B6N/B6N^ without the B6J-associated tRNA mutation. To generate Atoh1-Cre; *Hbs1l*^cKO^ mice lacking *n-Tr20*, B6.Cg-Tg(Atoh1-cre)1Bfri/J, B6N-*Hbs1l*^+/-^, and B6N-*Hbs1l*^fl/fl^ mice were crossed to B6J-*nTr20*^-/-^ mice. Subsequently, these strains were intercrossed to generate Atoh1-Cre; *Hbs1l* ^cKO^; *n-Tr20* ^-/-^ mice. For BrdU experiments, pregnant females or pups were injected with BrdU (0.05mg/g, Sigma-Aldrich, B9285) and collected 30 minutes (S-phase analysis) or 24 hours (cell cycle exit analysis) post injection. For the isolation of MEFs or embryos, the day that a vaginal plug was detected was defined as embryonic day 0.5 (E0.5).

For conditional knock out experiments in primary mouse embryonic fibroblasts (MEFs), we crossed mice to the tamoxifen inducible *Cre*-line B6.Cg-Tg(CAG-cre/Esr1*)5Amc/J (CAG-Cre^ER^, The Jackson Laboratory, #004682, MGI:2680708). The *Cre* allele was paternally inherited to generate embryos (F_2_ generation) because *Cre*-mediated recombination (“*leaky Cre expression*”) was occasionally observed even in the absence of tamoxifen (4-OHT) treatment of MEFs when the *Cre* allele was maternally inherited (F_2_ generation).

All experiments and quantifications were performed with at least three mice/embryos of each genotype and time point using mice of either sex (embryos were not sexed). The Jackson Laboratory Animal Care and Use Committee and The University of California San Diego Animal Care and Use Committee approved all mouse protocols.

### Strain abbreviation

For conditional knock out (cKO) experiments, animals were given a simplified abbreviation throughout the article. The complete genotype is shown below.

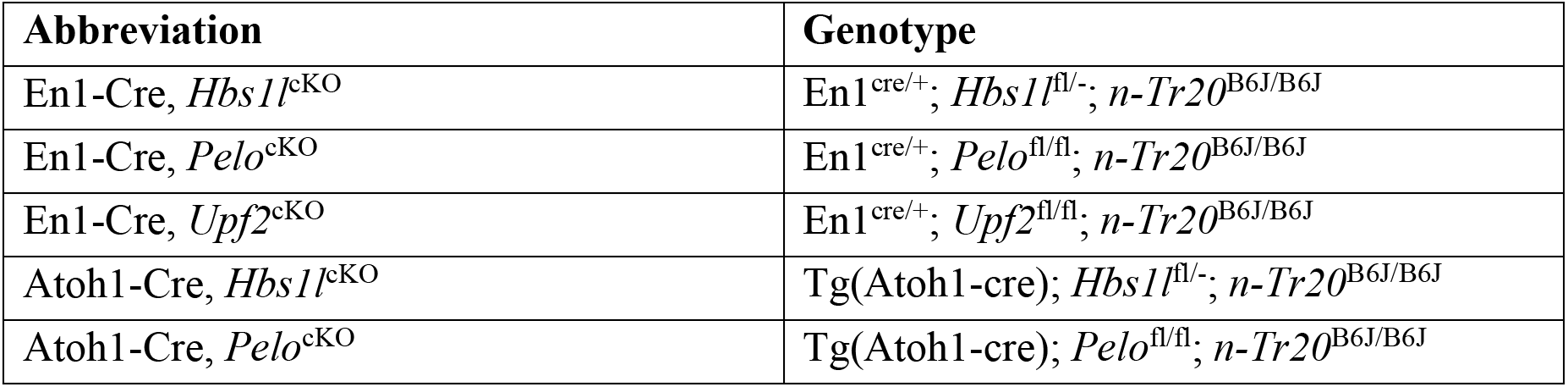
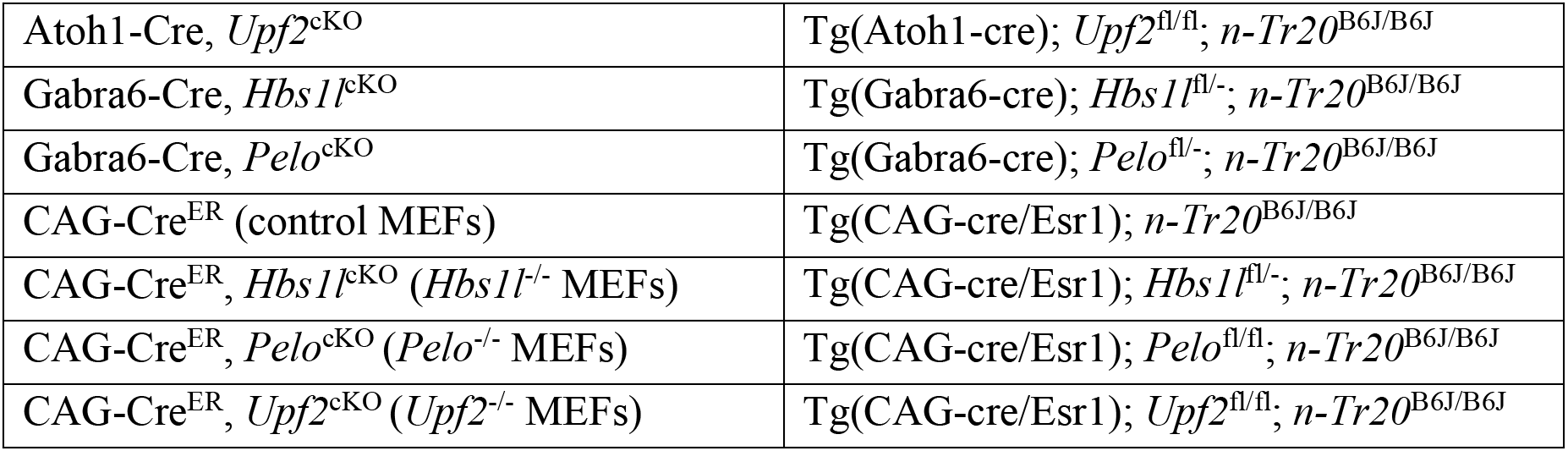

### Cell culture

Primary mouse embryonic fibroblasts (MEFs) were isolated on embryonic day E13.0 and prepared by standard procedures (Nagy et al., 2014). MEFs were maintained in Dulbecco’s modified Eagle’s medium (Gibco, #41965039) with GlutaMAX (Gibco, #35050061), PSN (Gibco, # 15640055), and 10% embryonic stem cell fetal bovine serum (Gibco, #10439024) at 37°C in 5% CO_2_. Two days post-isolation (Passage P0), the cell culture media was replaced with fresh media and supplemented with 1μM 4-OHT (4-hydroxytamoxifen, Sigma, H7904) for both control and mutant cells. After 48 hours, cells were washed, trypsinized, and seeded on a 10-cm dish (Passage P1). For RNA sequencing and ribosome footprint profiling experiments, cells were collected 48 hours (Passage P1, Day 2) later when cells reached ∼80% confluency. For western blotting experiments, cells of Passage P1 were collected 48 hours (Day 2) and 96 hours (Day 4) post-seeding.

### Histology and immunohistochemistry

Anesthetized mice were transcardially perfused with 4% paraformaldehyde (PFA, for immunofluorescence and histology), bouins (for histology) or 10% neutral buffered formalin (NBF, for *in situ* hybridization). Tissues were post-fixed overnight and embedded in paraffin. For histological analysis, sections were deparaffinized, rehydrated, and were stained with cresyl violet according to standard procedures. Histological slides were imaged using a digital slide scanner (Hamamatsu).

All quantifications were performed with three mice or embryos of each genotype and time point using animals of either sex (embryos were not sexed). For cell quantification in embryos at E12.5 to E16.5 and pups at P5 (Olig2^+^ cells), cells were counted from the entire section and values were averaged from three parasagittal (embryos) or sagittal (pups) sections (spaced 35μm apart) per mouse. For the analysis of granule cell precursors or granule cells in P5 or P6 pups, the total number of cells was determined within lobule IV/V or IX, and values were averaged from three sagittal sections (spaced 50μm apart) per mouse. For the analysis of granule cells in adult mice (35 weeks of age), granule cells were counted in a 0.025 mm^2^ area of lobule IX and values were averaged from three midline sections per mouse spaced 100μm apart.

For immunofluorescence, antigen retrieval on deparaffinized PFA-fixed sections was performed by microwaving sections in 0.01M sodium citrate buffer (pH 6.0, 0.05% Tween-20) for three times with three minutes each or three times for three minutes, followed by two times for nine minutes. PFA-fixed sections were incubated with the following primary antibodies overnight at 4°C: rabbit anti-cleaved caspase 3 (Cell signaling, #9661, 1:100), mouse anti-BrdU (Dako, M0744, 1:50), rabbit anti-Ki67 (Abcam, ab15580, 1:100), mouse anti-Lhx1/5 (DSHB, 4F2-c, 1:100), mouse anti-NeuN (Millipore, MAB377, 1:500), mouse anti-PCNA (Invitrogen, MA5-11358, 1:100), rabbit anti-p-S6^240/244^ (Cell Signaling, #5364, 1:1000), rabbit anti-Pax2 (ThermoFisher, 71-6000, 1:50), rabbit anti-Olig2 (Abcam, ab109186, 1:200), rabbit anti-Tbr1 (Chemicon, AB9616, 1:1000), and rabbit anti-pH3 (Upstate, 06-570, 1:1000). Immunofluorescence with antibodies to BrdU was performed on sections treated with DNase I (5mU/µl, Worthington, LS002139) for 45 minutes at 37°C after antigen retrieval. Detection of primary antibodies was performed with goat anti-mouse Alexa Fluor-488 or -555, goat anti-rabbit Alexa Fluor-488 or -555, and donkey anti-rabbit Alexa Fluor-555 secondary antibodies (Invitrogen). Sections were counterstained with DAPI and Sudan Black to quench autofluorescence.

For immunofluorescence quantification, the fluorescence intensity was measured in an area of 60×125μm using ImageJ, averaged from three sections (spaced 35μm apart) per embryo and expressed as the fold change relative to control.

### RNAscope (*in situ* hybridization)

*In situ* hybridization of *Hbs1l-C2* probes (ACD, #527471-C2) was performed with the ACD RNAscope Multiplex Fluorescent Reagent Kit v2 (ACD, #323100) using the manufacturer’s protocol. Briefly, deparaffinized NBF-fixed sections were treated for 15 minutes with Target Retrieval Reagent at 100°C and subsequently, treated with Protease Plus for 20 minutes (E13.5 embryos) or 30 minutes (P28 mice) at 40°C. RNAScope probes were hybridized for 2 hours, TSA^®^ Plus Cyanine 3 (PerkinElmer, 1:1,500) was used as a secondary fluorophore for *Hbs1l-C2* probes.

### Reverse transcription and quantitative PCR (qPCR) analysis

Whole mouse embryos, primary mouse embryonic fibroblasts, or adult mouse tissues were isolated and immediately frozen in liquid nitrogen. Total RNA was extracted with Trizol reagent (Life Technologies). cDNA synthesis was performed on DNase-treated (DNA-free DNA Removal Kit, Life Technologies AM1906) total RNA using oligo(dT) primers and SuperScript III First-Strand Synthesis System (Life Technologies). Quantitative RT-PCR reactions were performed using iQ SYBR Green Supermix (Bio-Rad) and an CFX96 Real-Time PCR Detection System (Bio-Rad). Expression levels of *β-actin* were used as input control for semi-quantitative RT-PCR. For quantitative RT-PCR (qPCR) analysis, expression levels of the genes of interest were normalized to *Gapdh* using the 2^-ΔΔCT^ method (Livak and Schmittgen, 2001) and expressed as the fold change + standard error of the mean (SEM) relative to control.

Semi-quantitative RT-PCR Primers (F, Forward; R, Reverse):

***Hbs1l Exon 3* F:**
5’GAAATTGACCAAGCTCGCCTGTA3’
***Hbs1l Exon 6* R:**
5’CTCAGAAGTTAAGCCAGGCACT3’
***β-actin* F:**
5’GGCTGTATTCCCCTCCATCG3’
***β-actin* R:**
5’CCAGTTGGTAACAATGCCATGT3’

Quantitative RT-PCR Primers (F, Forward; R, Reverse):

***Hbs1l* F**: 5’AGACCATGGGATTTGAAGTGC3’
***Hbs1l* R**: 5’CCGGTCTCAGGAATGTTAGGA3’
***Hbs1l II* F**: 5’TGAAGTTGAACAAAGTGCCAAG3’
***Hbs1l II* R**: 5’CTGCTTCCTCTGTGTTCCTC3’
***Pelo* F**: 5’CCCCAGGAAACGGAAAGGC3’
***Pelo* R**: 5’ACGCACTTTACAACCTCGAAG3’
***Gapdh* F**: 5’CATTGTCATACCAGGAAATG3’,
***Gapdh* R**: 5’GGAGAAACCTGCCAAGTATG3’

### Western blotting

MEFs or tissues were immediately frozen in liquid nitrogen. Proteins were extracted by homogenizing frozen tissue or cell samples in 5 volumes of RIPA buffer with cOmplete Mini, EDTA-free Protease inhibitor Cocktail (Roche), sonicating tissues two times for 10 seconds (Branson, 35% amplitude) or triturating cells 10 times using a 26G needle. Lysates were incubated for 30 minutes at 4°C, centrifuged at 16’000xg for 25 minutes, and 25µg of whole protein lysate were resolved on SDS-PAGE gels prior to transfer to PVDF membranes (GE Healthcare Life Sciences, #10600023) using a tank blotting apparatus (BioRad).

For detection of phosphoproteins, frozen tissue samples were homogenized in 5 volumes of homogenization buffer (50mM Hepes/KOH, pH 7.5, 140mM potassium acetate, 4mM magnesium acetate, 2.5mM dithiothreitol, 0.32M sucrose, 1mM EDTA, 2mM EGTA) (Carnevalli et al., 2004), supplemented with phosphatase and protease inhibitors (PhosStop and cOmplete Mini, EDTA-free Protease inhibitor Cocktail, Roche). Frozen cell samples were homogenized by using a 26G needle (5 times) in homogenization buffer (see above). Lysate samples were immediately centrifuged at 12,000xg for 7 minutes and whole protein lysate were resolved on SDS-PAGE gels prior to transfer to PVDF membranes.

After blocking in 5% nonfat dry milk (Cell Signaling, #9999S), blots were probed with primary antibodies at 4°C overnight: rabbit anti-phospho-eIF2*α*^S51^ (Cell Signaling, #9721, 1:1000), rabbit anti-eIF2*α*(Cell Signaling, #9722, 1:2000), rabbit anti-Hbs1l (Proteintech, 10359-1-AP, 1;1000), rabbit anti-Pelo (Proteintech, 10582-1-AP, 1:2000), rabbit anti-GAPDH (Cell Signaling, #2118, 1:10,000), rabbit anti-phospho-p70S6K^T389^ (Cell Signaling, #9234, 1:1000), rabbit anti-phospho-p70S6K (Cell Signaling, #2708, 1:1000), mouse anti-vinculin (Sigma, V-9131, 1:20,000). Primary antibodies were detected by incubation with HRP-conjugated secondary antibodies for 2 hours at room temperature: goat anti-rabbit IgG (BioRad, #170-6515) or goat anti-mouse IgG (BioRad, #170-6516). Signals were detected with SuperSignal West Pico Chemiluminescent substrate (ThermoScientific, #34080) or ProSignal Femto ECL Reagent (Genesee Scientific, #20-302).

### RNA sequencing library construction

For RNA sequencing of primary mouse embryonic fibroblasts (MEFs), one 10-cm dish of tamoxifen (4-OHT) treated MEFs at Day 2 (Passage P1) from CAG-Cre^ER^ (control) CAG-Cre^ER^; *Hbs1l*^cKO^ (*Hbs1l*^-/-^), CAG-Cre^ER^; *Pelo*^cKO^ (*Pelo*^-/-^) and CAG-Cre^ER^; *Upf2*^cKO^ (*Upf2*^-/-^) were collected. One 10-cm dish from one individual embryo was used as one biological replicate, and either two (*Pelo*) or three (control, *Hbs1*, *Upf2*) biological replicates (individual embryos) were used per genotype. Two micrograms of DNase-treated (DNA-free DNA Removal Kit, Life Technologies AM1906) total RNA were used for the RNA library construction, performed as per the manufacturer’s protocol (KAPA Stranded mRNA-Seq Kit, KR0960) and the adapter ligation was performed for 3 hours at room temperature. Library quality and concentration was assessed using D1000 screen tape on the Agilent TapeStation and Qubit 2.0 Fluorometer. All libraries were pooled, and the pool of libraries was sequenced on 2 lanes using HiSeq4000 (PE100).

### Data analysis of RNA sequencing data

Reads were quantified using kallisto version 0.42.4 (Bray et al., 2016) and pseudo-aligned to a Gencode M24 transcriptome reference with parameters –bias and -b 100. Differential expression was performed using sleuth version 0.30.0 (Pimentel et al., 2017). Pairwise comparisons were performed to identify differentially expressed transcripts and genes: CAG-Cre^ER^ vs. CAG-Cre^ER^; *Pelo*^cKO^, CAG-Cre^ER^ vs. CAG-Cre^ER^; *Hbs1l*^cKO^, and CAG-Cre^ER^ vs. CAG-Cre^ER^; *Upf2*^cKO^. We used functions within sleuth to perform differential transcript and gene expression. Briefly, we fit null models and models corresponding to the genotype of the samples for each transcript and performed Wald tests on the models for each transcript to identify differentially expressed transcripts. Differential gene expression was performed by aggregating transcript expression on Ensembl gene identifiers. Multiple hypotheses testing was corrected using Benjamini-Hochberg correction, referred to as q-value. Transcript biotypes were identified using biomaRt version 2.42.1 (Durinck et al., 2005). For downstream TE analysis, mapping to mm10 using a Gencode M24 transcript annotation was performed using hisat2 version 2.1.0 (Kim et al., 2019) using default parameters.

### Ribosome profiling library construction

Ribosome profiling libraries were generated as previously described (Ingolia et al., 2012; Ishimura et al., 2014) with some modifications. Cerebella were dissected and immediately frozen in liquid nitrogen. One cerebellum from P14 mice was used for each biological replicate, and three biological replicates were prepared for each control (Atoh1-Cre; *Hbs1l*^+/+^) and mutant (Atoh1-Cre; *Hbs1l*^cKO^) genotype. For profiling of mouse embryos, embryos from timed mating were dissected at embryonic day E8.5, frozen in a drop of nuclease free water on dry ice, and then flash frozen in liquid nitrogen. Five embryos were pooled for each biological replicate, and three biological replicates were prepared for each control (*Hbs1l*^+/+^) and mutant (*Hbs1l*^-/-^) genotype. For profiling of primary mouse embryonic fibroblasts (MEFs), tamoxifen (4-OHT) treated MEFs at Day 2 (Passage P1) from CAG-Cre^ER^ (control), CAG-Cre^ER^; *Hbs1l*^cKO^ (*Hbs1l*^-/-^), CAG-Cre^ER^; *Pelo*^cKO^ (*Pelo*^-/-^) and CAG-Cre^ER^; *Upf2*^cKO^ (*Upf2*^-/-^) were collected. Two 10-cm dishes from one individual embryo were pooled as one biological replicate, and three biological replicates (three individual embryos) were used for each genotype. The tissue and embryo homogenization were performed with a mixer mill (Retsch MM400) in lysis buffer (20mM Tris-Cl, pH 8.0, 150 mM NaCl, 5mM MgCl_2_, 1mM DTT, 100µg/ml CHX, 1% (v/v) TritonX-100, 25units/ml Turbo DNase I). The cell homogenization was performed by triturating the cells in lysis buffer (20mM Tris-Cl, pH 8.0, 150 mM NaCl, 5mM MgCl_2_, 1mM DTT, 100µg/ml CHX, 1% (v/v) TritonX-100, 25units/ml Turbo DNase I) ten times through a 26G needle. RNase I-treated lysates were overlaid on top of a sucrose cushion in 5ml Beckman Ultraclear tubes and centrifuged in an SW55Ti rotor for 4 hours at 4°C at 46,700 rpm. Pellets were resuspended and RNA was extracted using the miRNeasy kit (Qiagen) according to manufacturer’s instructions. 26-34 nts (cerebella samples) or 15-34 nts (embryo and MEF samples) RNA fragments were purified by electrophoresis on a denaturing 15% gel. Linker addition, cDNA generation (first-strand synthesis was performed at 50°C for 1 hour), circularization, rRNA depletion, and amplification of cDNAs with indexing primers were performed. Library quality and concentration was assessed using high sensitivity D1000 screen tape on the Agilent TapeStation and Qubit 2.0 Fluorometer. All libraries were pooled as set of six samples consisting of three control and mutant samples. Libraries were run on HiSeq4000 (SR75) and either 3 lanes (cerebella samples) or 2 lanes (embryo and MEF samples) per set of samples were sequenced.

### Data analysis of ribosome profiling data

Reads were clipped to remove adaptor sequences (CTGTAGGCACCATCAAT) using fastx_clipper and trimmed so that reads start on the second base using fastx_trimmer (http://hannonlab.cshl.edu/fastx_toolkit/). Reads containing ribosomal RNAs, snoRNAs, and tRNAs were then filtered out by mapping to a ribosomal RNA reference using bowtie2 version 2.2.3 using parameters -L 13 (Langmead and Salzberg, 2012). Remaining reads were mapped to an mm10 mouse reference using a Gencode M24 annotation, or a Gencode M24 protein-coding transcript reference using hisat2 version 2.1.0 (Kim et al., 2019). Ribosomal A-sites were identified using RiboWaltz version 1.0.1 (Lauria et al., 2018), and reads with lengths of 27-34 nucleotides were retained for further analysis.

To identify potential codon bias in the A-site, observed/expected reads were calculated for each transcript with alignments with the expected reads being the read density expected at a given site with a given codon, assuming that reads are uniformly distributed across the coding part of the transcript. Differences in codon usage were then tested with a student’s t-test followed by Benjamini-Hochberg multiple hypothesis testing correction.

Pauses were identified using previous methodology (Ishimura et al., 2014). Reads with lengths of 27-34 nucleotides were analyzed using a 0.5 reads/codon threshold in all samples to analyze pausing on transcripts. Additionally, pauses with P-sites nearby start and stop codons (P-sites at the -3, 0,1,2,3, and 4 positions for start codons and P-sites at the -1 and -2 positions for stop codons) in any isoform of the gene were excluded from the analysis. To identify pauses overlapping start and stop codons in other isoforms, we used ensembldb v.2.6.8 (Rainer et al., 2019) to map transcript coordinates back to the genome and analyzed whether pauses with identical genome coordinates overlapped start/stop codons.

To extend this approach to identify pauses at the gene level, we removed start/stop overlapping pauses (in any gene-matched transcript) and collapsed pauses based on the genomic location of the A-site, which was also identified with ensembldb v.2.6.8. Mean pause score across transcripts of genomically identical pauses for each sample was the reported pause score (excluding transcripts without a reported pause in a sample). If genomically identical pauses were in multiple transcript regions (i.e. 3’UTR or CDS depending on the isoform), all were reported.

Pauses were further filtered such that the pause locus appeared in all three replicates of the genotype and thereby, pauses that were only detected in one or two of three replicates of the genotype were removed from the analysis. Pauses were compared between the 4-OHT treated knock out (CAG-Cre^ER^; *Hbs1l*^cKO^, CAG-Cre^ER^; *Pelo*^cKO^ and CAG-Cre^ER^; *Upf2*^cKO^) and the control (CAG-Cre^ER^) sample that each mutant sample was pooled with for sequencing.

To identify ribosome footprint reads with 3’end As (untemplated reads), we extracted at first reads from a genome mapped bam file with six or more As at the 3’end of the read (3’end As were soft-clipped). Mapped reads were only considered if the 3’end A reads were not matching the reference sequence, while all unmapped reads with six or more As at the 3’ end were considered. Afterwards, the 3’end As were then removed from these untemplated reads and they were mapped back to the transcriptome using parameters described above for ribosome footprint profiling mapping to the transcriptome. Only the untemplated reads that mapped after the removal of the 3’end As were considered as ribosome footprints that may derive from ribosomes translating premature polyadenylated transcripts.

For differential RPF and differential TE analysis, genome mapped reads were quantified using featureCounts (Liao et al., 2014) with footprints overlapping CDS features. For TE analysis specifically, RNA-seq read pairs overlapping gene exon features were also quantified using featureCounts. Differential RPF analysis was performed using DESeq2 (v1.26.0) (Love et al., 2014) comparing 4-OHT treated knock out (CAG-Cre^ER^; *Hbs1l*^cKO^, CAG-Cre^ER^; *Pelo*^cKO^ and CAG-Cre^ER^; *Upf2*^cKO^) to control (CAG-Cre^ER^) cells using default parameters. Histone mRNAs were removed from the analysis by removing gene names with the prefixes “Hist”, “H1f”, “H2a”, “H2b”, “H3” and “H4”. TE was quantified and tested for using riborex version 2.3.4 using the DESeq2 engine (Li et al., 2017).

### Pathway analysis

Data were analyzed using Ingenuity Pathway Analysis (IPA, QIAGEN Inc., https://www.qiagenbioinformatics.com/products/ingenuity-pathway-analysis).

Kyoto Encyclopedia of Genes and Genomes (KEGG) pathway analysis was performed using the ShinyGO v0.61 bioinformatics web server (http://bioinformatics.sdstate.edu/go) (Ge et al., 2020)

by uploading the gene lists from our RNA sequencing or ribosome profiling analysis. Pathway enrichment terms with a *P*-value cutoff (FDR) ≤ 0.05 were considered enriched.

### Statistics

For quantification of protein expression, RNA expression (quantitative RT-PCR), fluorescence intensity or histological quantifications, *P*-values were computed in GraphPad Prism using either student’s t-test, multiple t-tests, one-way ANOVA, or two-way ANOVA and statistical tests were corrected for multiple comparisons as indicated in the figure legends. All quantifications were performed with at least three mice/embryos of each genotype and time point using mice of either sex (embryos were not sexed).

### Data availability

The RNA sequencing and ribosome footprint data have been made available (GSE162556).

## Acknowledgements

We thank T. Jucius and A. Kano for technical assistance, Drs. Bo Torben Porse and Miles Wilkinson for providing the *Upf2*^fl/fl^ mice, the IGM Genomics Center at the University of California San Diego (UCSD) for support with RNA sequencing and the UCSD School of Medicine Microcopy Core for providing access to microscopy equipment (Grant P30 NS047101). This work was supported in part by NIH R01 NS094637 (SLA). SLA is an investigator of the Howard Hughes Medical Institute.

## Author Contributions

MT and SLA designed experiments and wrote the manuscript. MT performed mouse and molecular biology experiments under SLA’s guidance. SIA performed the computational analysis of RNA-sequencing and ribosome profiling data under JHC’s guidance.

## Declaration of Interests

The authors declare no competing interests.

## Key Resource Table

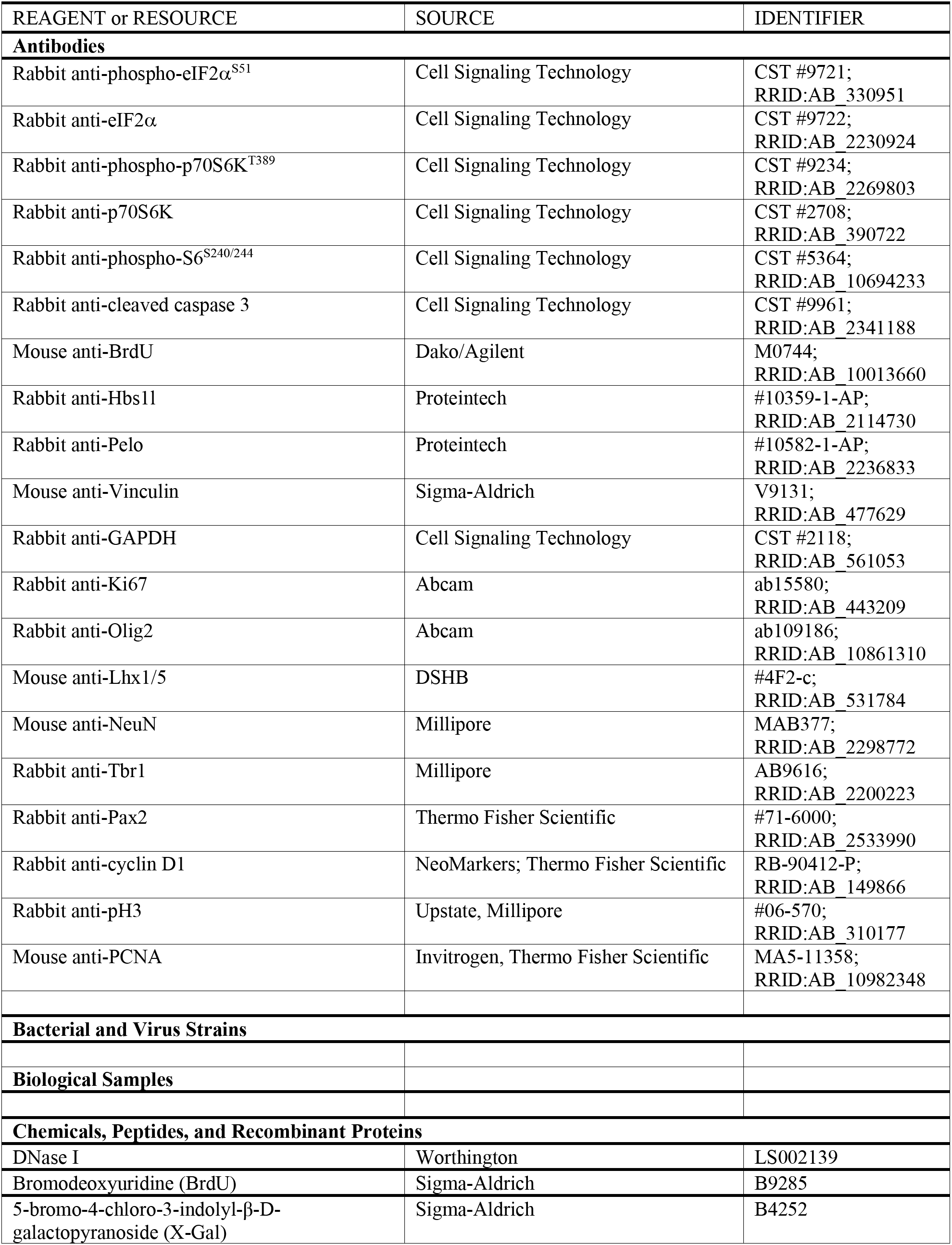
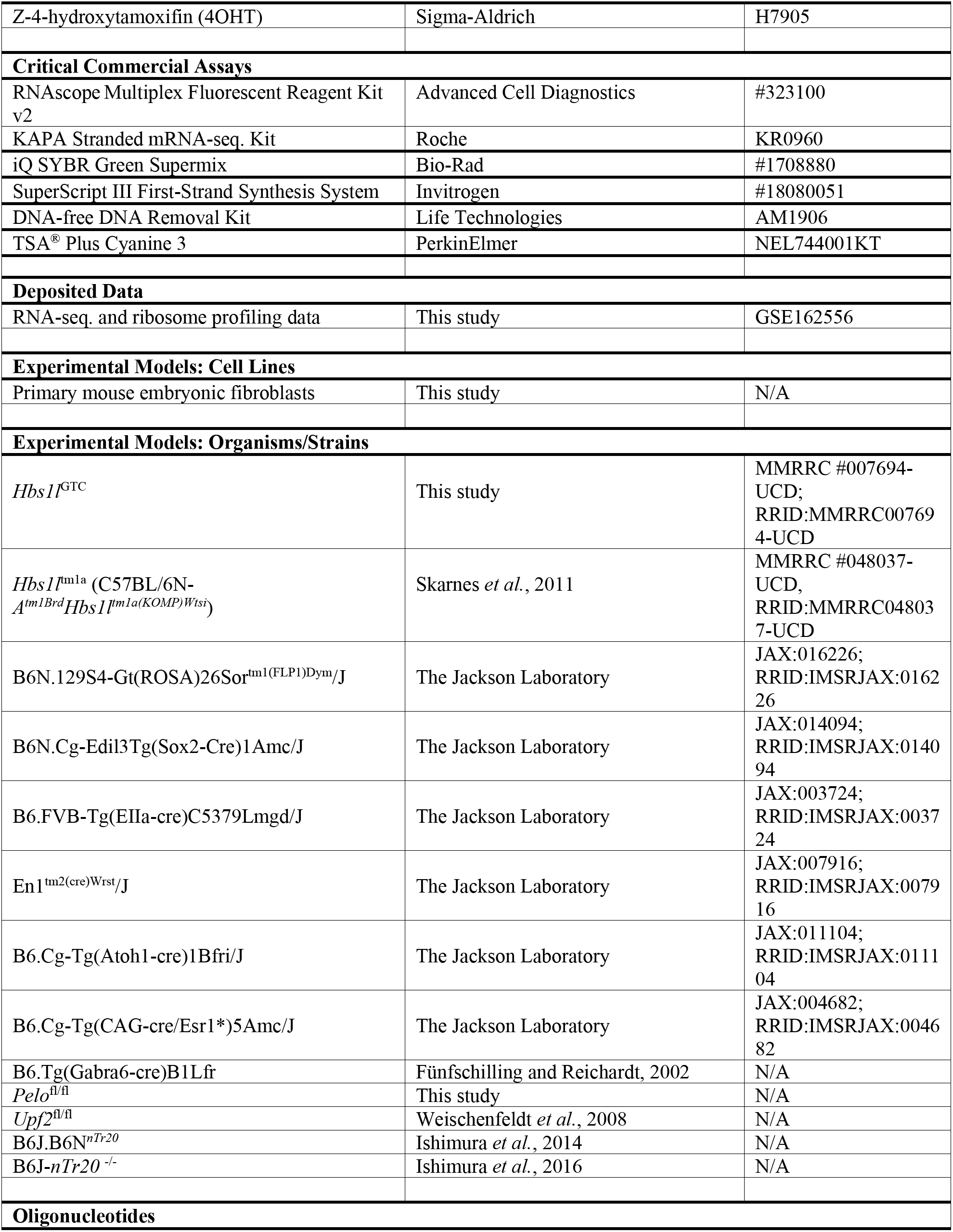
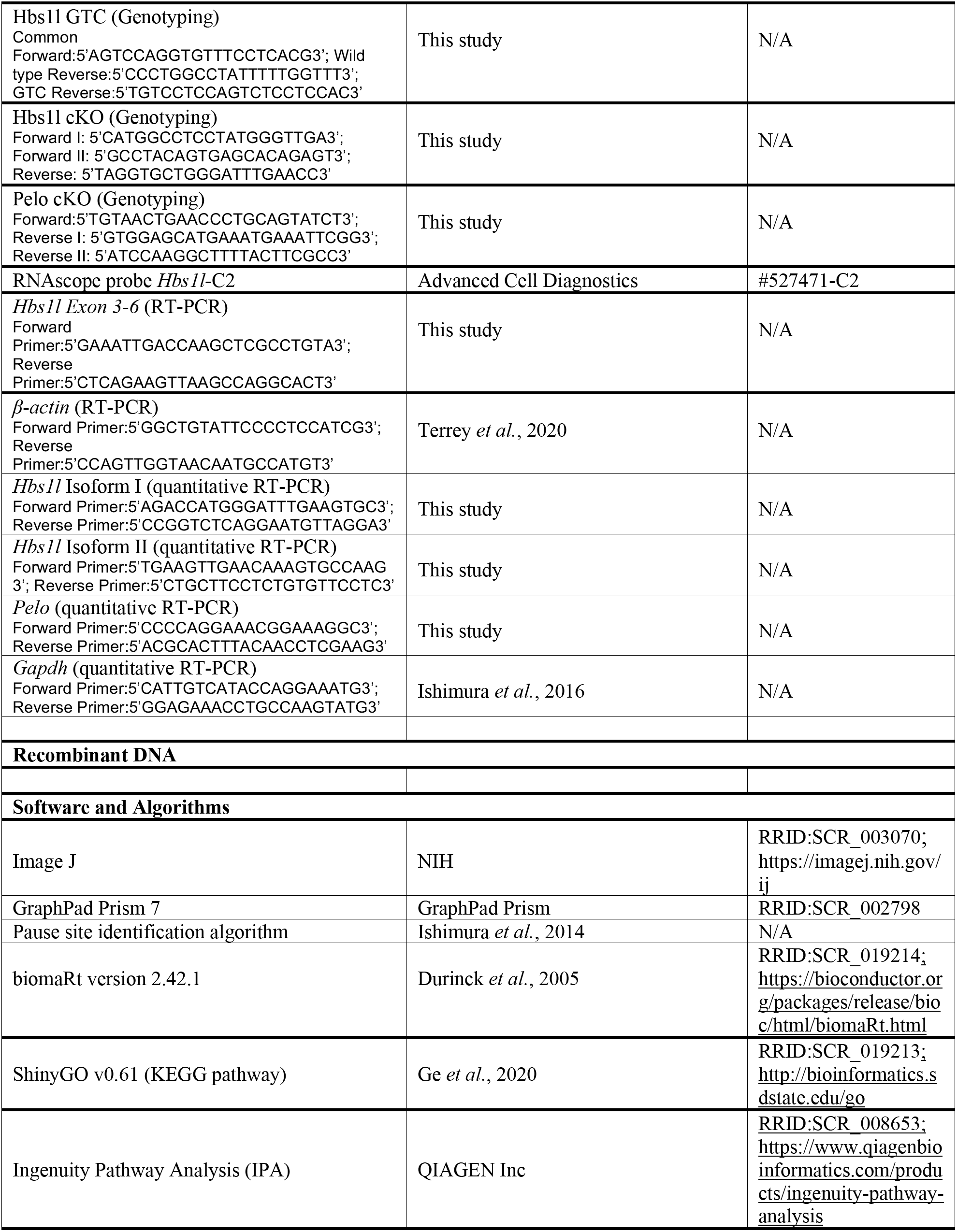
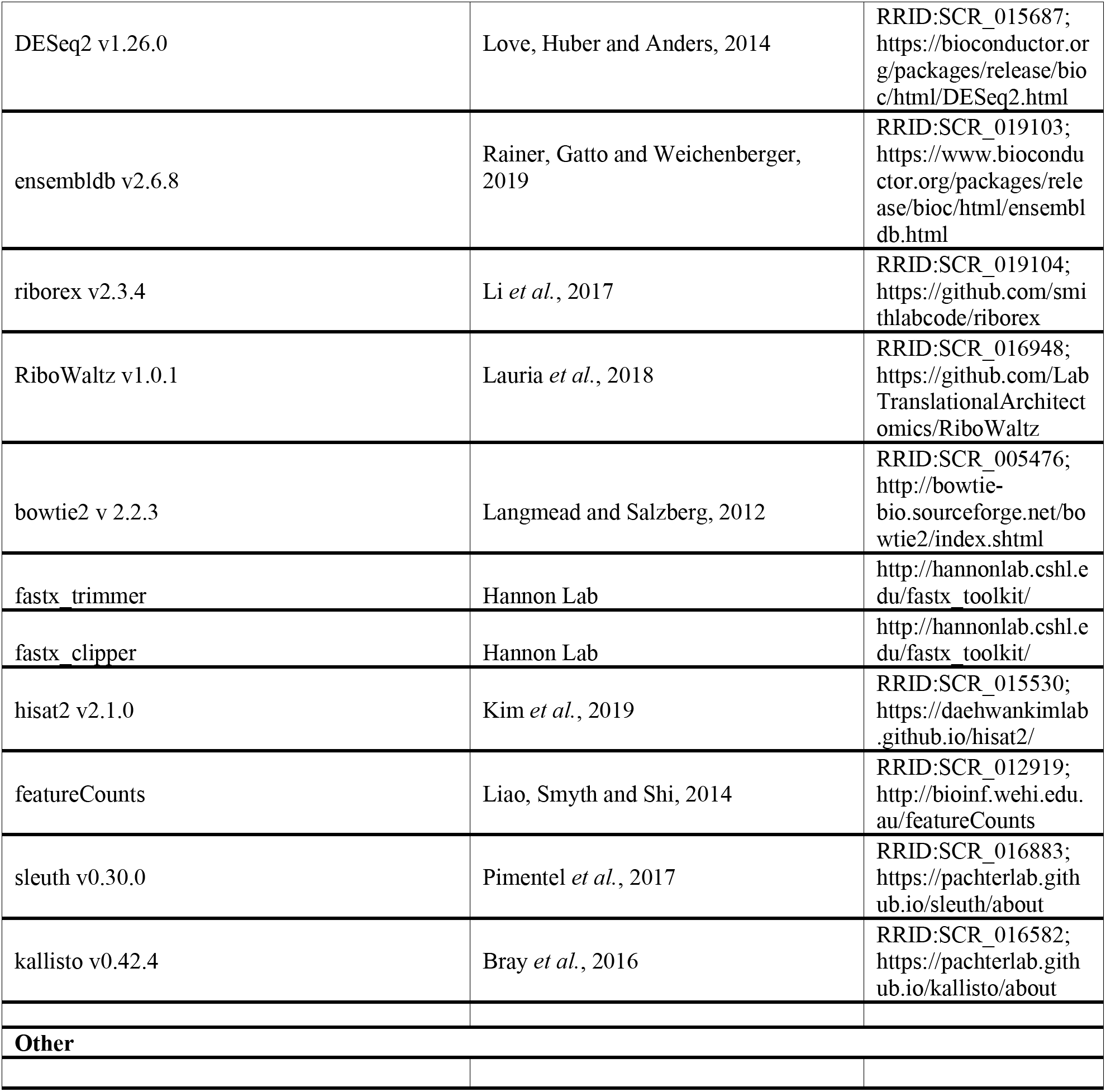

**Supplementary file 1.** Embryonic lethality.

**Supplementary file 2.** Locus specific pausing genome level.

**Supplementary file 3.** DE RPF footprints.

**Supplementary file 4.** A-site pausing.

**Supplementary file 5.** Locus specific pausing transcript level.

**Supplementary file 6.** TE MEFs.

**Supplementary file 7.** DE mRNA MEFs.

---

**Figure 2 - figure supplement 1.**
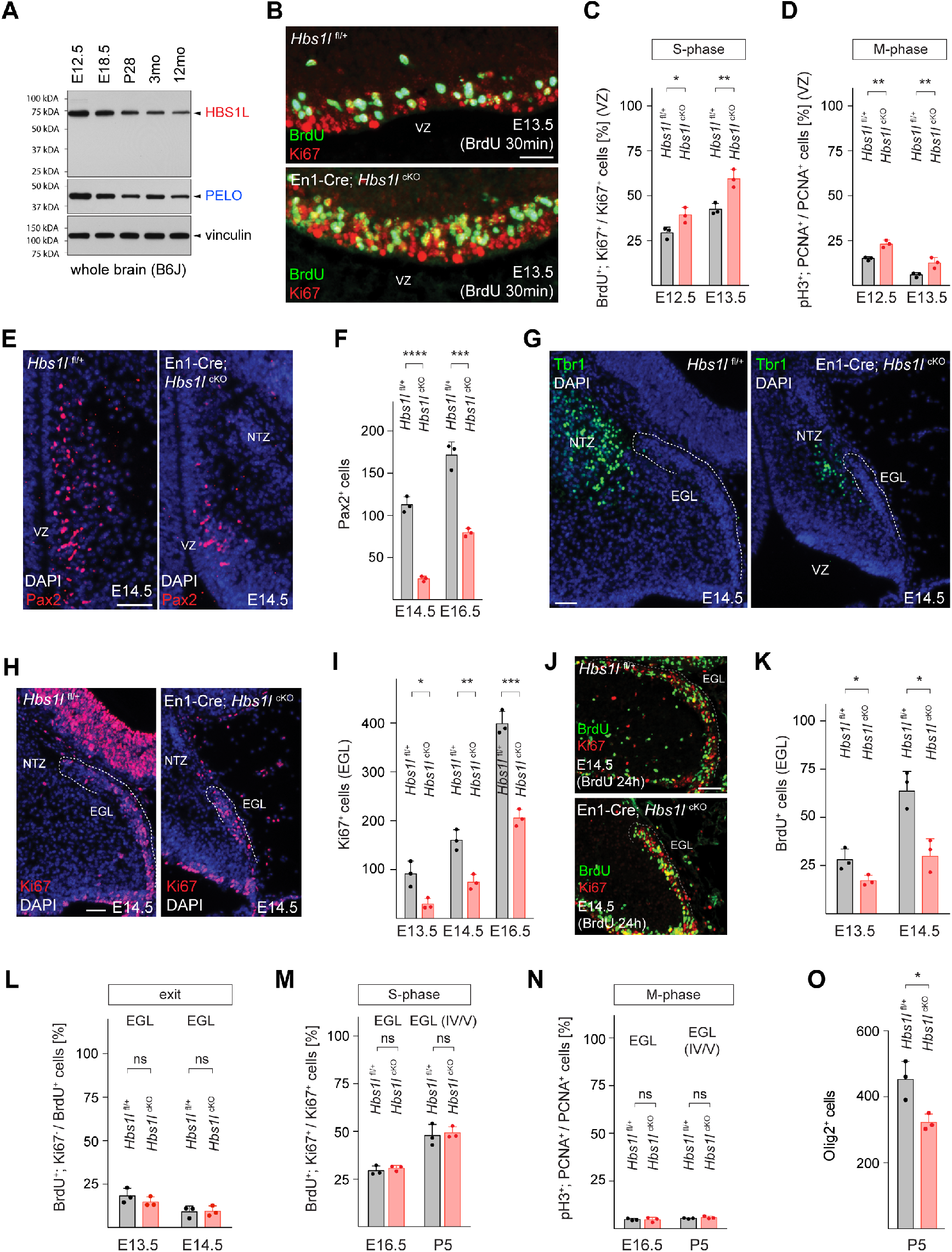
*Hbs1l* is required for the development of multiple cerebellar linages. (A) Western blot analysis using brain lysates from B6J mice. Vinculin was used an input control. (B) Immunofluorescence using antibodies to BrdU (green) and Ki67 (red) from cerebellar sections of E13.5 control (*Hbs1l*^fl/+^) and En1-Cre; *Hbs1l*^cKO^ embryos to determine the fraction of cells in S-phase. Embryos were injected with BrdU 30 minutes prior to harvest. Sections were counterstained with DAPI. (C) Percentage of cerebellar VZ-progenitors in S-phase (BrdU^+^, Ki67^+^ cells). Data represent mean + SD. (D) Percentage of cerebellar VZ-progenitors in M-phase (pH3^+^, PCNA^+^ cells). Data represent mean + SD. (E) Immunofluorescence using antibodies to Pax2 (red) on cerebellar sections from E13.5 control (*Hbs1l*^fl/+^) and En1-Cre; *Hbs1l*^cKO^ embryos. Sections were counterstained with DAPI. (F) Number of cerebellar VZ-derived interneuron precursors (Pax2^+^ cells). Data represent mean + SD. (G) Immunofluorescence using antibodies to Tbr1 (green) on cerebellar sections from E14.5 control (*Hbs1l*^fl/+^) and En1-Cre; *Hbs1l*^cKO^ embryos. Sections were counterstained with DAPI. (H) Immunofluorescence using antibodies to Ki67 (green) on cerebellar sections from E14.5 control (*Hbs1l*^fl/+^) and En1-Cre; *Hbs1l*^cKO^ embryos to identify granule cell precursors in the EGL. Sections were counterstained with DAPI. (I) Number of RL-derived granule cell precursors in the EGL (Ki67^+^ cells). Data represent mean + SD. (J) Immunofluorescence using antibodies to BrdU (green) and Ki67 (red) on cerebellar sections from E14.5 control (*Hbs1l*^fl/+^) and En1-Cre; *Hbs1l*^cKO^ embryos. Embryos were injected with BrdU 24 hours prior to harvest. (J) Number of RL-derived granule cell precursors (BrdU^+^ cells). Data represent mean + SD. (L) Percentage of RL-derived granule cell precursors that exited the cell cycle (BrdU^+^, Ki67^-^ cells). Data represent mean + SD. (M) Percentage of RL-derived granule cell precursors in S-phase (BrdU^+^, Ki67^+^ cells). Embryos were injected with BrdU 30 minutes prior to harvest. Data represent mean + SD. (N) Percentage of RL-derived granule cell precursors in M-phase (pH3^+^, PCNA^+^ cells). Data represent mean + SD. (O) Number of oligodendroglial progenitors (Olig2^+^ cells) from E14.5 control (*Hbs1l*^fl/+^) and En1-Cre; *Hbs1l*^cKO^ cerebella. Scale bars: 20μm (C); 50μm (E, G, H, J). VZ, ventricular zone; NTZ; nuclear transitory zone; EGL, external granule cell layer; RL, rhombic lip. t-tests were corrected for multiple comparisons using Holm-Sidak method (C, D, F, I, K, L, M, N); Student’s t-test (O). ns, not significant; * *P* ≤ 0.05; ** *P* ≤ 0.01; *** *P* ≤ 0.001; **** *P* ≤ 0.0001.

**Figure 2 - figure supplement 2.**
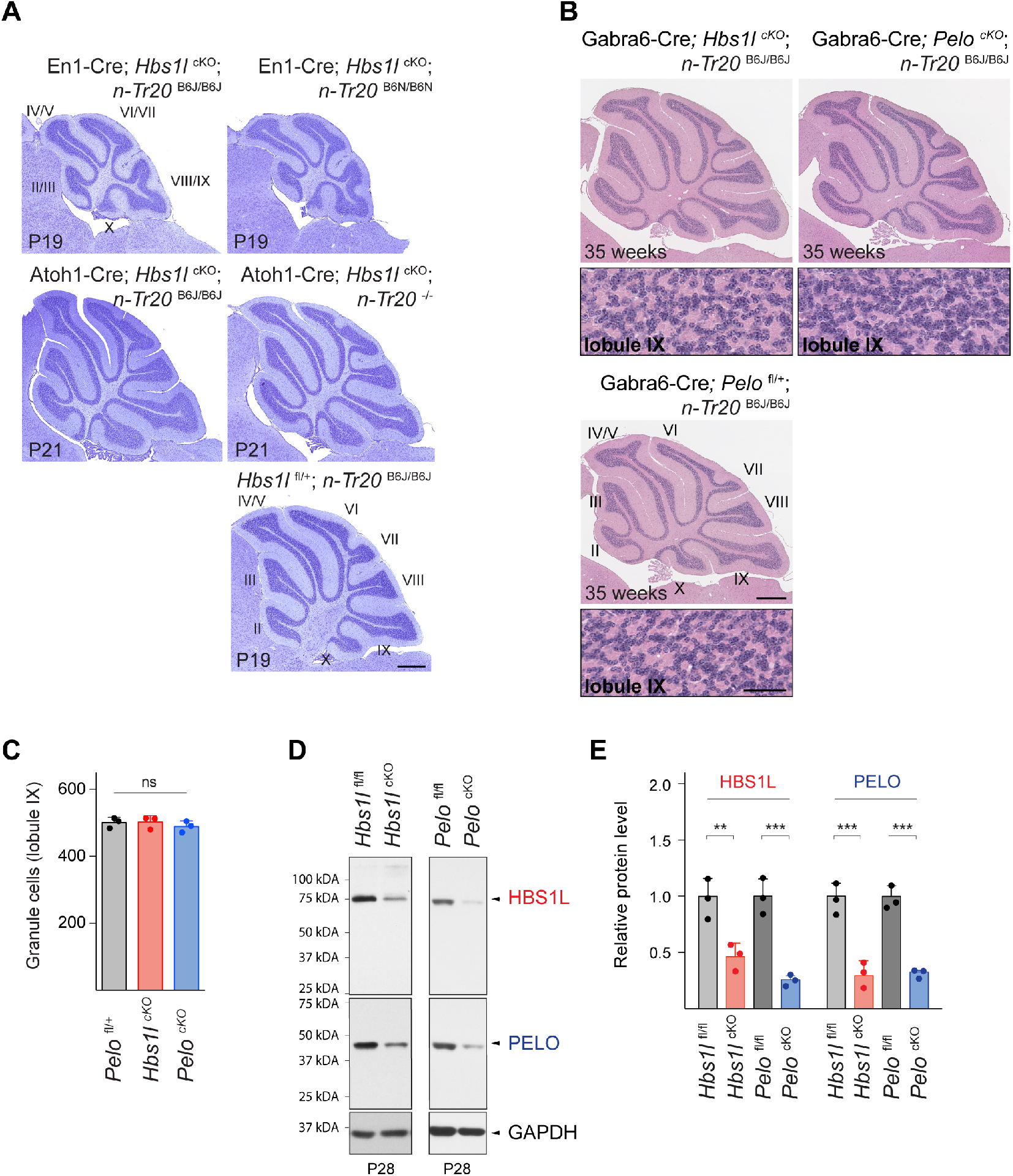
*Hbs1l^-/-^-*mediated cerebellar defects are independent of the B6J-associated mutation in *n-Tr20*. (A) Cresyl violet staining of sagittal sections of En1-Cre; *Hbs1l*^cKO^ cerebellum expressing wild type levels of *n-Tr20* (*n-Tr20*^B6N/B6N^) or reduced levels of *n-Tr20* (*n-Tr20*^B6J/B6J^) and Atoh1-Cre; *Hbs1l*^cKO^ cerebella with reduced levels of *n-Tr20* (*n-Tr20*^B6J/B6J^) or that completely lack *n-Tr20* (*n-Tr20*^-/-^). Cerebellar lobules are indicated by Roman numerals. (B) Sagittal sections from 35-week-old control (Gabra6-Cre; *Pelo*^fl/+^; *n-Tr20*^B6J/B6J^), Gabra6-Cre; *Pelo*^cKO^; *n-Tr20*^B6J/B6J^ and Gabra6-Cre; *Hbs1l*^cKO^; *n-Tr20*^B6J/B6J^ cerebella stained with hematoxylin and eosin. Higher magnification images of lobule IX are shown below. Cerebellar lobules are indicated by Roman numerals. (C) Number of cerebellar granule cells in lobule IX shown in B. Data represent mean + SD. (D) Western blot analysis using cerebellar lysates from P28 control (*Hbs1l*^fl/fl^; *n-Tr20*^B6J/B6J^ or *Pelo*^fl/fl^; *n-Tr20*^B6J/B6J^), Gabra6-Cre; *Pelo*^cKO^; *n-Tr20*^B6J/B6J^ and Gabra6-Cre; *Hbs1l*^cKO^; *n-Tr20*^B6J/B6J^ mice. GAPDH was used an input control. (E) Quantification of HBS1L and PELO protein levels shown in D and were normalized to levels of vinculin. Protein levels are relative to those of controls (*Hbs1l*^fl/fl^ or *Pelo*^fl/fl^) from each cross. Scale bars: 500μm (A); 500μm and 50μm (higher magnification) (B). One-way ANOVA (C) and two-way ANOVA (E) were corrected for multiple comparisons using Tukey method. ns, not significant; ** *P* ≤ 0.01; *** *P* ≤ 0.001.

**Figure 3 - figure supplement 1.**
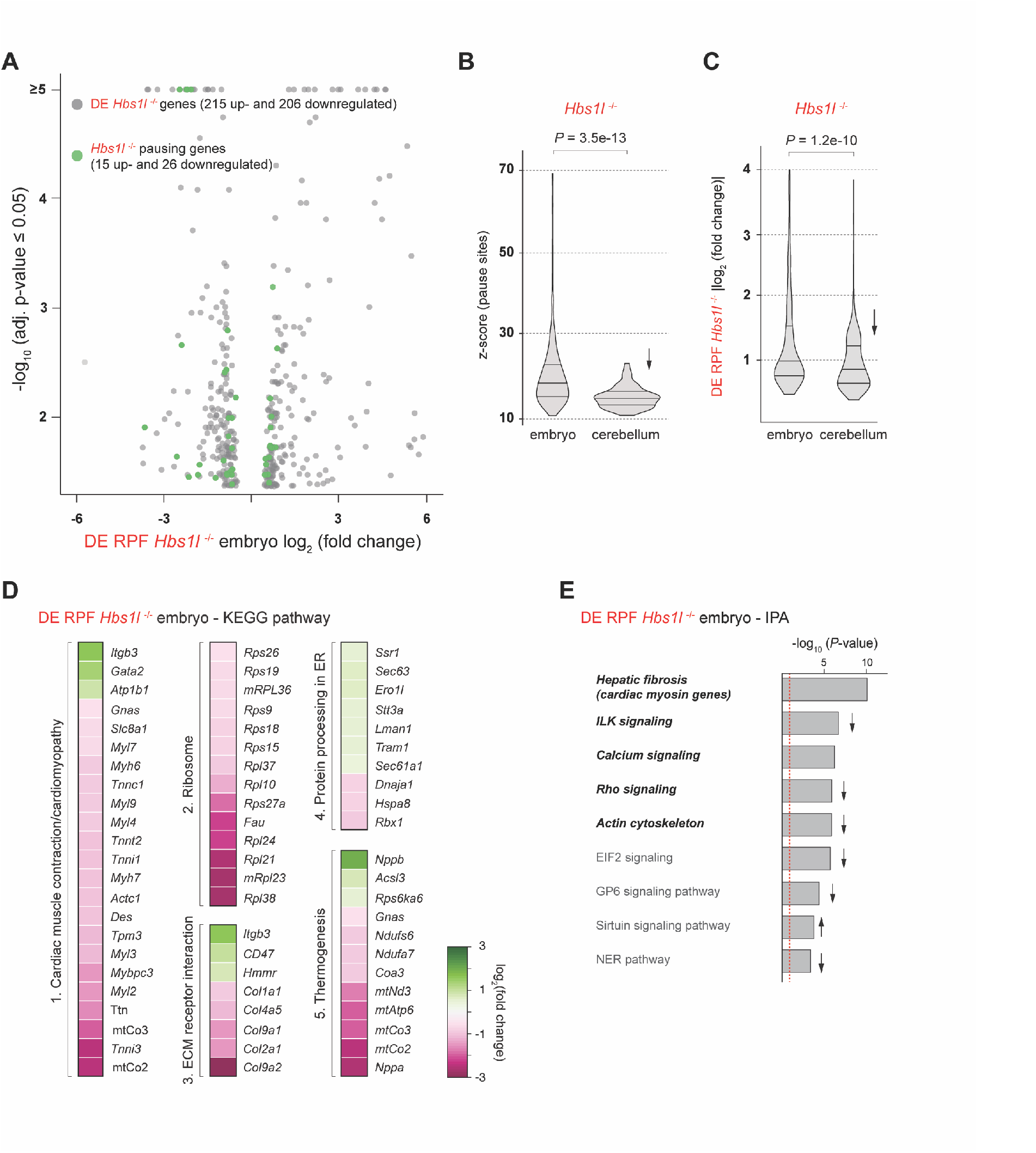
*Hbs1l* deficiency in embryos alters translation of pathways associated with heart function. (A) Analysis of differentially translated genes (DE RPF *Hbs1l*^-/-^, adj. *P* ≤ 0.05) of E8.5 *Hbs1l*^-/-^ embryos (421 genes, grey). Genes that have ribosome pauses specific to *Hbs1l*^-/-^ embryos with significant (adj. *P* ≤ 0.05) changes in translation are shown in green (47 genes). (B) Comparison of z-scores of *Hbs1l*^-/-^-specific pause sites detected in E8.5 *Hbs1l*^-/-^ embryos and P14 Atoh1-Cre; *Hbs1l*^cKO^ cerebella. Downward direction of arrow indicates significant lower z-scores of these pauses in P14 Atoh1-Cre; *Hbs1l*^cKO^ cerebella. (C) Comparison of differentially translated genes (DE RPF *Hbs1l*^-/-^, adj. *P* ≤ 0.05) of E8.5 *Hbs1l*^-/-^ embryos and P14 Atoh1-Cre; *Hbs1l*^cKO^ cerebella. Downward direction of arrow indicates significant lower fold changes in translation in P14 Atoh1-Cre; *Hbs1l*^cKO^ cerebella. (D) KEGG pathway analysis of differentially translated genes (DE RPF *Hbs1l*^-/-^, adj. *P* ≤ 0.05) of E8.5 *Hbs1l*^-/-^ embryos. Top five significantly (*P* ≤ 0.05) enriched pathways are shown and the heatmap indicates relative changes in genes associated with the respective pathway. (E) Ingenuity Pathway Analysis (IPA) of differentially translated genes (DE RPF *Hbs1l*^-/-^) from E8.5 *Hbs1l*^-/-^ embryos. Pathways that are involved in heart, muscle contraction, cardiomyocyte and cytoskeleton function are in italics. The red dashed line indicates the significance threshold (*P* = 0.05). Up- or downward direction of arrow indicates predicated up- or downregulated activity of the pathway, respectively. RPF, ribosome protected fragments. Wilcoxon rank-sum test was used to determine statistical significance (B, C).

**Figure 4 - figure supplement 1.**
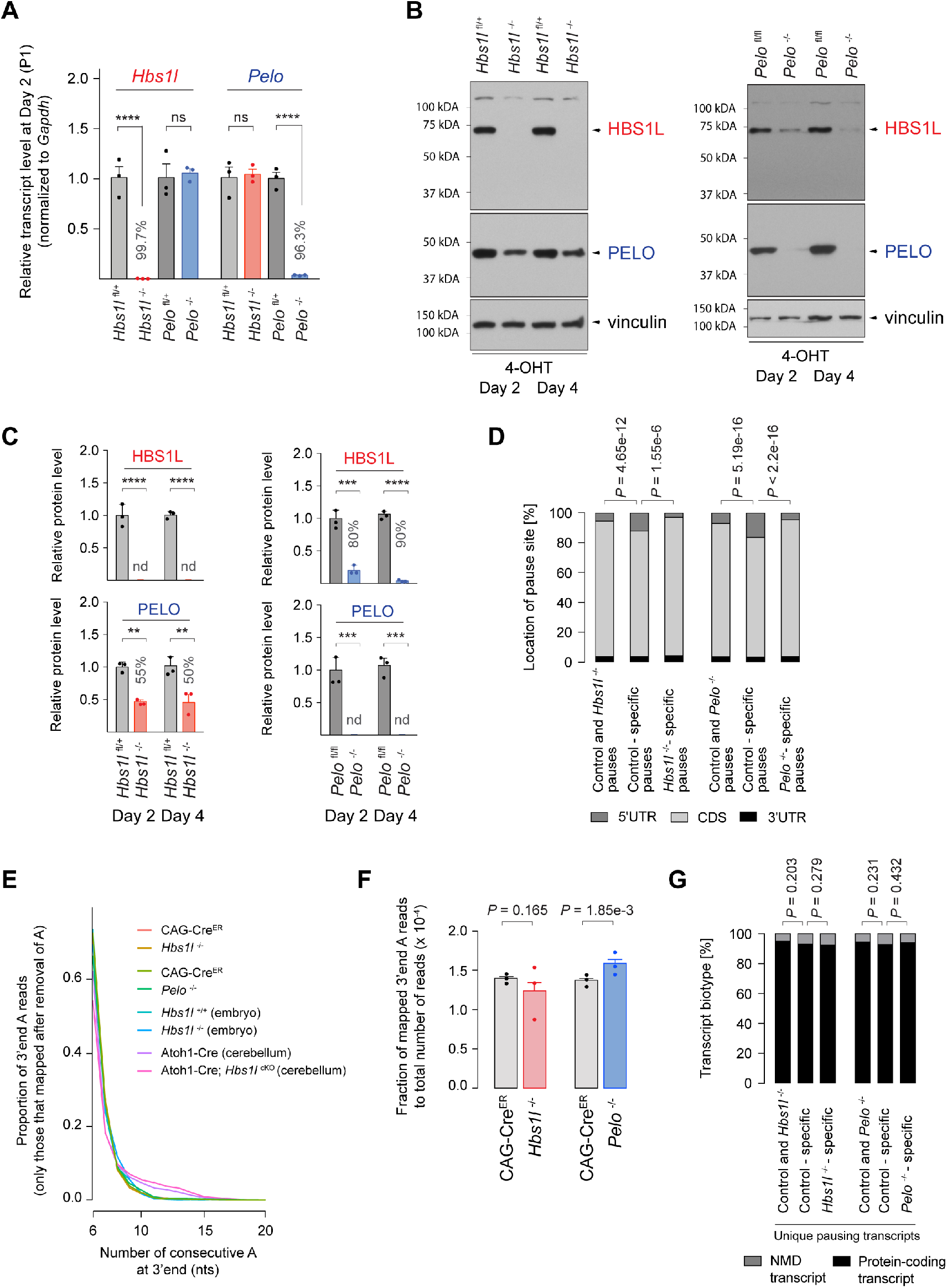
Loss of *Hbs1l* and *Pelo* control protein levels of their respective binding partner. (A) Quantitative RT-PCR analysis of *Hbs1l* and *Pelo* transcripts using cDNA from MEFs of tamoxifen-treated control (*Hbs1l*^fl/+^ or *Pelo*^fl/+^), *Hbs1l*^-/-^ and *Pelo*^-/-^ cells at Day 2 (Passage P1). Data were normalized to *Gapdh* and fold change in expression is relative to those of control (*Hbs1l*^fl/+^ or *Pelo*^fl/+^) cells. Data represent mean + SEM. (B) Western blot analysis of HBS1L and PELO using MEF lysates of tamoxifen-treated control (*Hbs1l*^fl/+^ or *Pelo*^fl/fl^), *Hbs1l*^-/-^ or *Pelo*^-/-^ cells at Day 2 and 4 (Passage P1). Vinculin was used as an input control. (C) Quantification of HBS1L and PELO protein levels shown in B and were normalized to levels of vinculin. Protein levels are relative to those of control (*Hbs1l*^fl/+^ or *Pelo*^fl/fl^) cells at Day 2 (Passage P1). Data represent mean + SD. (D) Location of pause sites. (E) Histogram of A count on the 3’end of ribosome footprints that mapped after removal of the 3’end A’s. (F) Fraction of 3’end A ribosome footprints that mapped after removal of the 3’end A’s relative to the total number of ribosome footprints from tamoxifen treated control (CAG-Cre^ER^) and *Hbs1l*^-/-^, or control (CAG- Cre^ER^) and *Pelo*^-/-^ cells. (G) Percent of unique pausing transcripts that are protein-coding or NMD transcripts. P1, Passage 1; MEFs, primary mouse embryonic fibroblasts; 4-OHT, 4-hyrdoxytamoxifin; PTC, premature termination codon; nd, not detected; A, adenosines; nts, nucleotides; CDS, coding sequence; UTR, untranslated region; NMD, nonsense-mediated decay. Two-way ANOVA was corrected for multiple comparisons using Tukey method (A, C); Fisher’s exact test (D, G); Student’s t-test (F). ns, not significant; ** *P* ≤ 0.01; *** *P* ≤ 0.001; **** *P* ≤ 0.0001.

**Figure 5 - figure supplement 1.**
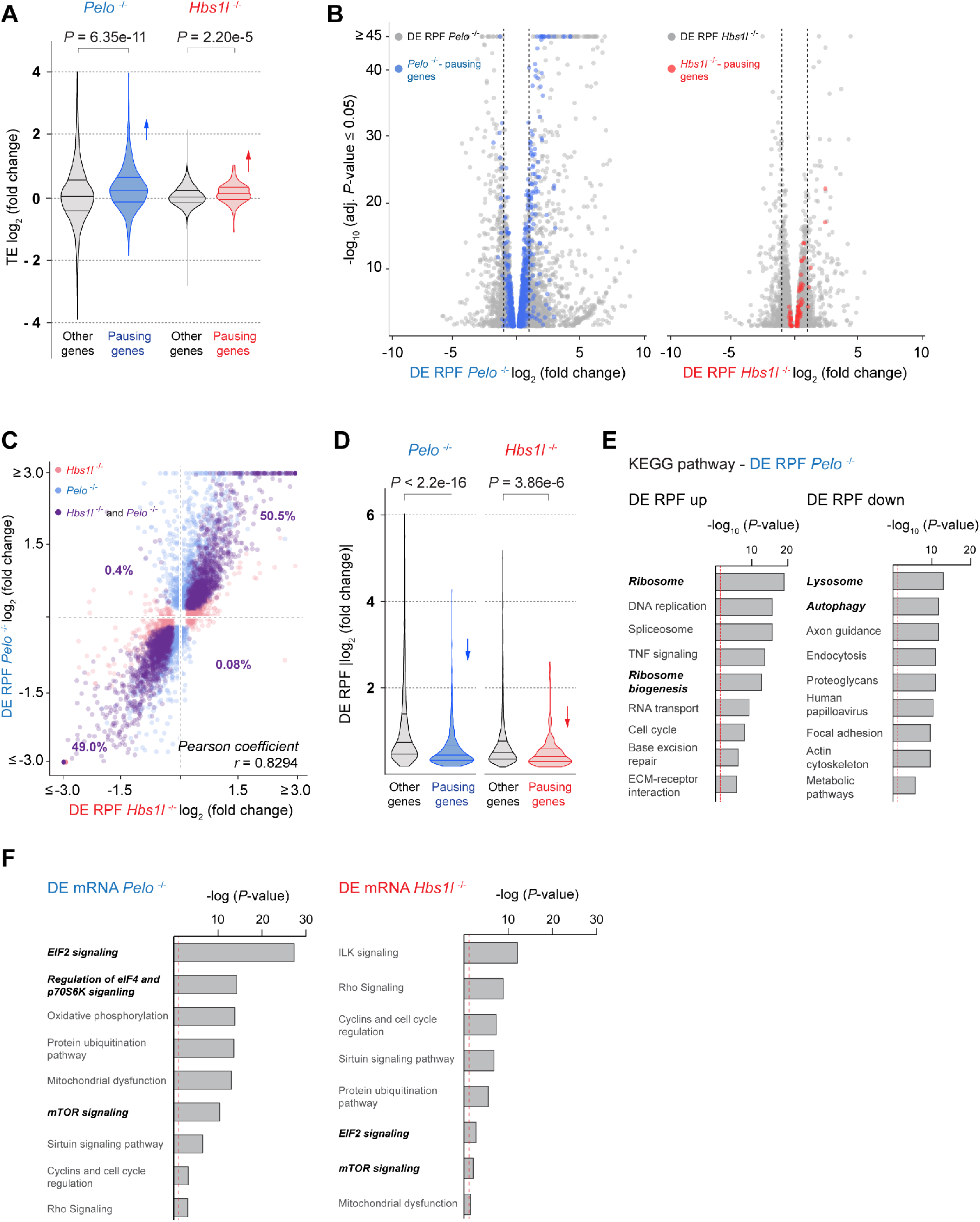
*Pelo*/*Hbs1l* deficiency alters translational gene expression of multiple pathways. (A) Translational efficiency (TE) of *Pelo*^-/-^- and *Hbs1l*^-/-^-pausing genes was compared to the TE of the remaining (‘other’) genes from *Pelo*^-/-^ (blue) and *Hbs1l*^-/-^ (red) MEFs. Upward direction of arrows indicates significant increase in translational efficiency of *Pelo*^-/-^- and *Hbs1l*^-/-^-pausing genes. (B) Analysis of differentially translated genes in *Pelo*^-/-^ (DE RPF *Pelo*^-/-^, adj. *P* ≤ 0.05, left) and *Hbs1l*^-/-^ (DE RPF *Hbs1l*^-/-^, adj. *P* ≤ 0.05, right) MEFs are shown in grey. *Pelo*^-/-^- and *Hbs1l*^-/-^-specific pausing genes (z-score ≥ 10, detected in all 3 biological replicates) that are differentially translated are highlighted. Two-fold changes in expression are indicated by the black dashed lines. (C) Differentially translated genes in *Hbs1l*^-/-^ MEFs (DE RPF *Hbs1l*^-/-^, adj. *P* ≤ 0.05, x-axis, red) were plotted against genes that are differentially translated in *Pelo*^-/-^ MEFs (DE RPF *Pelo*^-/-^, adj. *P* ≤ 0.05, y-axis, blue). Genes those translation was significantly different in both *Hbs1l*^-/-^ and *Pelo*^-/-^ MEFs are shown in purple. (D) Differential translation of genes in *Pelo* (DE RPF *Pelo*^-/-^, adj. *P* ≤ 0.05) and *Hbs1l* (DE RPF *Hbs1l*^-/-^, adj. *P* ≤ 0.05) was compared to that of *Pelo*^-/-^- and *Hbs1l*^-/-^-specific pausing genes. Downward direction of arrows indicates significantly reduced changes in RPFs from the pausing genes compared to non-pausing (‘other’) genes. (E) KEGG pathway analysis of differentially translated (up- and downregulated) genes (DE RPF *Pelo*^-/-^, adj. *P* ≤ 0.05) in *Pelo*^-/-^ MEFs. Top nine of the significantly enriched pathways are shown. The red dashed line indicates the significance threshold (*P* = 0.05). Pathways that are positively (ribosome, ribosome biogenesis) and negatively (lysosome, autophagy) regulated by mTORC1 are in italics. (F) Ingenuity pathway analysis (IPA) of differentially transcribed genes in *Pelo* (DE mRNA *Pelo*^-/-^) and *Hbs1l* (DE mRNA *Hbs1l*^-/-^). EIF2 and mTOR/p70S6K signaling are highlighted in yellow and magenta, respectively. The red dashed line indicates the significance threshold (*P* = 0.05). RPF, ribosome protected fragments. Wilcoxon rank-sum test was used to determine statistical significance (A, D); Pearson coefficient (*r*) was determined to analyze linearity of gene expression changes (C).

**Figure 6 - figure supplement 1.**
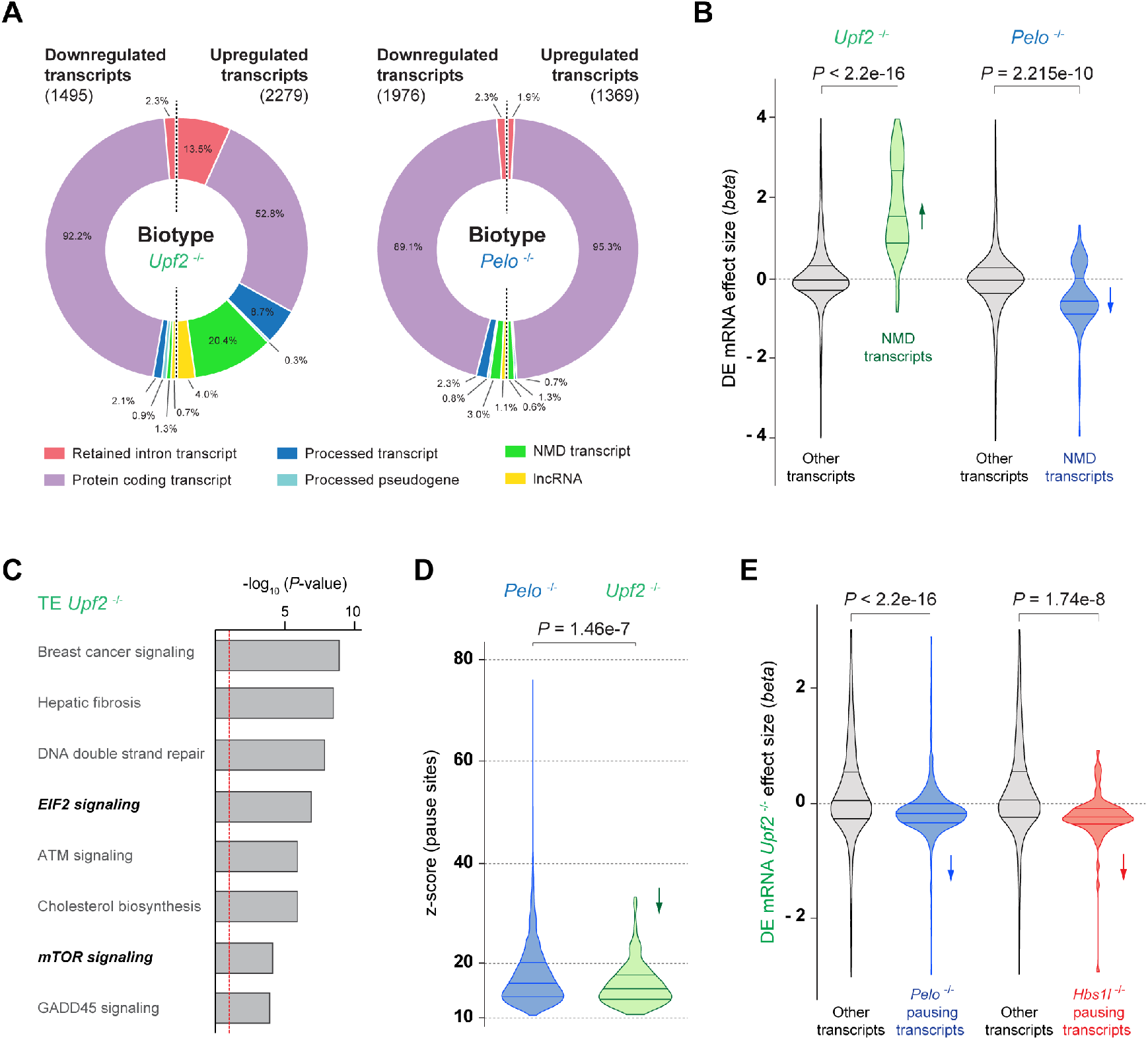
Loss of *Upf2* leads to an increase of NMD targets. (A) Ensembl-annotated transcript biotypes of differentially transcribed mRNAs in *Upf2*^-/-^ (DE mRNA *Upf2*^-/-^, *q*-value ≤ 0.05) and *Pelo*^-/-^ (DE mRNA *Pelo*^-/-^, *q*-value ≤ 0.05) MEFs. (B) Transcriptional expression of NMD transcripts with significant (*q*-value ≤ 0.05) changes in expression in *Upf2*^-/-^ (DE mRNA *Upf2*^-/-^) and *Pelo*^-/-^ (DE mRNA *Pelo*^-/-^) was compared to expression of the remaining (‘other’) transcripts. The effect size (*beta*) is analogous to the natural log fold change in expression. Up- or downward direction of arrows indicates significant increase or decrease in expression of NMD transcripts in *Upf2*^-/-^ or *Pelo*^-/-^ MEFs. (C) Ingenuity pathway analysis (IPA) of genes with differential translational efficiency (TE) in *Upf2*^-/-^ MEFs (TE *Upf2*^-/-^). EIF2 and mTOR signaling are in italics. The red dashed line indicates the significance threshold (*P* = 0.05). (D) Comparison of z-scores (‘pause score’) of specific pauses observed in *Pelo*^-/-^ (blue) and *Upf2*^-/-^ (green) MEFs. Downward direction of arrow indicates significantly lower pause scores specific to *Upf2*^-/-^ compared to *Pelo*^-/-^ MEFs. (E) Transcriptional expression of either *Pelo*- or *Hbs1l*-specific pausing transcripts (unique protein-coding pausing transcripts) was compared to the remaining (‘other’) transcripts in *Upf2*^-/-^ (DE mRNA *Upf2*^-/-^) MEFs. The effect size (*beta*) is analogous to the natural log fold change in expression. Downward direction of arrows indicates a significant decrease in expression of *Pelo*- or *Hbs1l*-specific pausing transcripts in *Upf2*^-/-^ MEFs. NMD, nonsense-mediated transcript; lncRNA, long noncoding RNA. Wilcoxon rank-sum test was used to determine statistical significance (B, D, E).

**Figure 7 - figure supplement 1.**
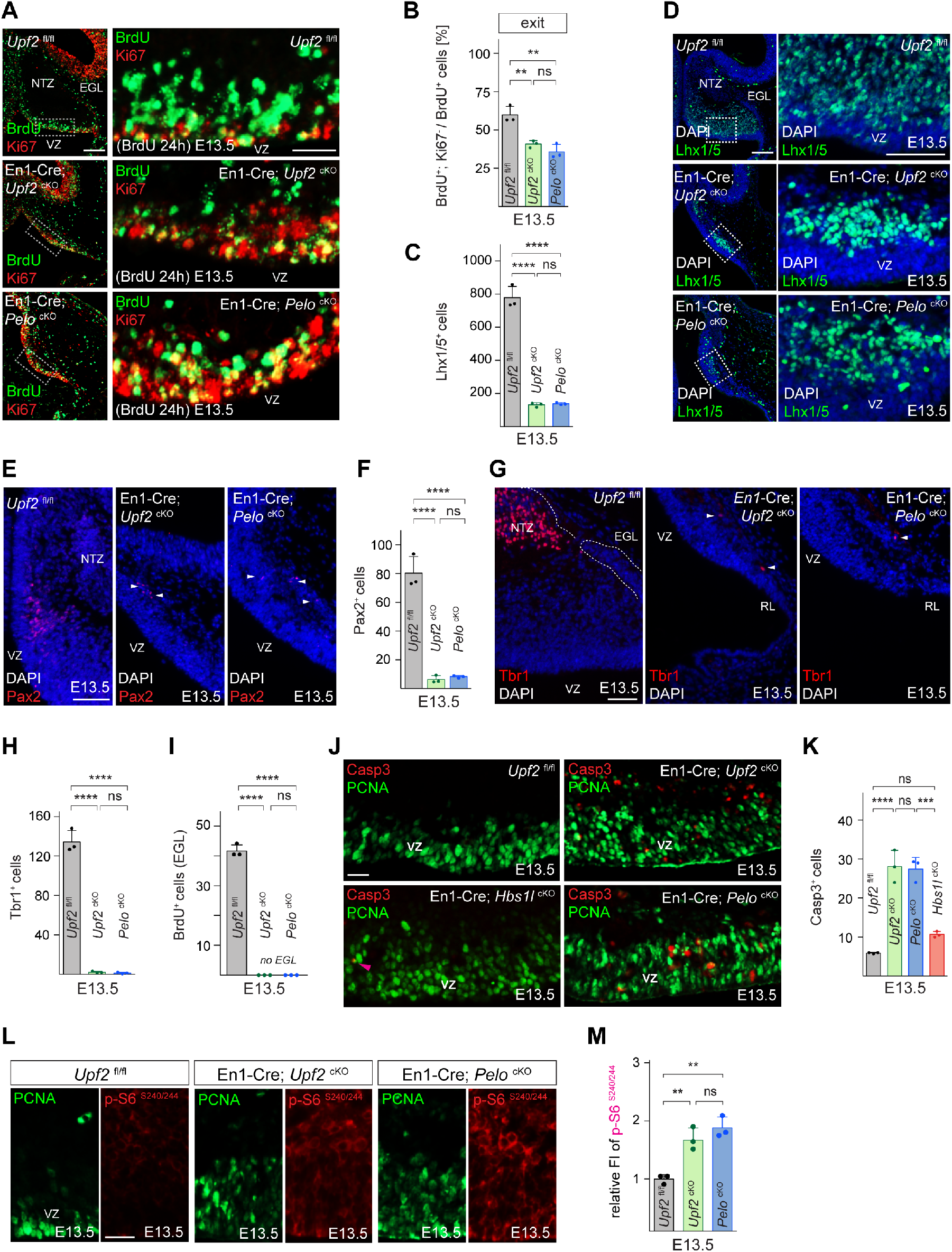
*Upf2* and *Pelo* are required for the development of multiple cerebellar linages. (A) Immunofluorescence using antibodies to BrdU (green) and Ki67 (red) on cerebellar sections from E13.5 control (*Upf2*^fl/fl^), En1-Cre; *Upf2*^cKO^ and En1-Cre; *Pelo*^cKO^ embryos to determine the fraction of cells in VZ that exited the cell cycle. Embryos were injected with BrdU 24 hours prior to harvest. (B) Percentage of cerebellar VZ-progenitors that exited the cell cycle (BrdU^+^, Ki67^-^ cells). Data represent mean + SD. (C) Number of cerebellar Purkinje cell precursors (Lhx1/5^+^ cells). Data represent mean + SD. (D) Immunofluorescence using antibodies to Lhx1/5 on cerebellar sections from E13.5 control (*Upf2*^fl/fl^), En1-Cre; *Upf2*^cKO^ and En1-Cre; *Pelo*^cKO^ embryos. Sections were counterstained with DAPI and higher magnification images of boxed areas are shown. (E) Immunofluorescence using antibodies to Pax2 (red) on cerebellar sections from E13.5 control (*Upf2*^fl/fl^), En1-Cre; *Upf2*^cKO^ and En1-Cre; *Pelo*^cKO^ embryos. Sections were counterstained with DAPI. Arrowheads denote occasional Pax2^+^ cells. (F) Number of cerebellar VZ-derived interneuron precursors (Pax2^+^ cells). Data represent mean + SD. (G) Immunofluorescence using antibodies to Tbr1 (red) on cerebellar sections from E13.5 control (*Upf2*^fl/fl^), En1-Cre; *Upf2*^cKO^ and En1-Cre; *Pelo*^cKO^ embryos. Sections were counterstained with DAPI. Arrowheads denote occasional Tbr1^+^ cells. (H) Number of cerebellar RL-derived Tbr1^+^ deep neurons. Data represent mean + SD. (I) Quantification of newborn granule cell precursors (BrdU^+^ cells) in E13.5 control (*Upf2*^fl/fl^), En1-Cre; *Upf2*^cKO^ and En1-Cre; *Pelo*^cKO^ EGL. Data represent mean + SD. Embryos were injected with BrdU 24 hours prior to harvest. (K) Immunofluorescence using antibodies to cleaved caspase 3 (Casp3, red) on cerebellar sections from E13.5 control (*Upf2*^fl/fl^), En1-Cre; *Upf2*^cKO^, En1-Cre; *Pelo*^cKO^ and En1-Cre; *Hbs1l*^cKO^ embryos. Sections were counterstained with DAPI. Note: Because the number of apoptotic cells was noticeably higher in En1-Cre; *Upf2*^cKO^ and En1-Cre; *Pelo*^cKO^ cerebella, arrowheads (magenta) to indicate these cells were not included. (J) Number of Casp3^+^ cells. Data represent mean + SD. (L) Immunofluorescence on sections of E13.5 control (*Upf2*^fl/fl^), En1-Cre; *Upf2*^cKO^ and En1-Cre; *Pelo*^cKO^ cerebellum with antibodies to p-S6^S240/244^ (red) and PCNA (green). (M) Relative fluorescence intensity (FI) of p-S6^S240/244^ of control (*Upf2*^fl/fl^), En1-Cre; *Upf2*^cKO^ and En1-Cre; *Pelo*^cKO^ embryos. Levels of p-S6^S240/244^ are relative to those of control (*Upf2*^fl/fl^). Data represent mean + SD. Scale bars: 100μm and 20μm (higher magnification) (A); 100μm and 50μm (higher magnification) (D); 50μm (E, G); 20μm (J); 25μm (L). VZ, ventricular zone; RL, rhombic lip; NTZ; nuclear transitory zone; EGL, external granule cell layer. One-way ANOVA was corrected for multiple comparisons using Tukey method (B, C, F, H, I, K, M). ns, not significant; ** *P* ≤ 0.01; *** *P* ≤ 0.001; **** *P* ≤ 0.0001.

## Notes

### Competing Interest Statement

The authors have declared no competing interest.

## References

Abreua R de S, Penalvaa LO, Marcotteb EM, Vogel C. 2009. Global signatures of protein and mRNA expression levels. Energy Convers Manag 5:1512–1526. doi:10.1039/b908315d

Adham IM, Sallam MA, Steding G, Korabiowska M, Brinck U, Hoyer-Fender S, Oh C, Engel W. 2003. Disruption of the Pelota Gene Causes Early Embryonic Lethality and Defects in Cell Cycle Progression. Mol Cell Biol 23:1470–1476. doi:10.1128/mcb.23.4.1470-1476.2003

Arpat AB, Liechti A, de Matos M, Dreos R, Janich P, Gatfield D. 2020. Transcriptome-wide sites of collided ribosomes reveal principles of translational pausing. Genome Res 7:985– 999. doi:10.1101/710061

Arribere JA, Fire AZ. 2018. Nonsense mRNA suppression via nonstop decay. Elife 7:1–23. doi:10.7554/eLife.33292

Blair JD, Hockemeyer D, Doudna JA, Bateup HS, Floor SN. 2017. Widespread translational remodeling during human neuronal differentiation. Cell Rep 21:2005–2016. doi:10.1016/j.celrep.2017.10.095

Blanco S, Bandiera R, Popis M, Hussain S, Lombard P, Aleksic J, Sajini A, Tanna H, Cortés-Garrido R, Gkatza N, Dietmann S, Frye M. 2016. Stem cell function and stress response are controlled by protein synthesis. Nature 534:335–340. doi:10.1038/nature18282

Brandman O, Hegde RS. 2016. Ribosome-associated Protein Quality Control. Nat Struct Mol Biol 23:7–15. doi:10.1038/nsmb.3147

Bray NL, Pimentel H, Melsted P, Lior Pachter. 2016. Near-optimal probabilistic RNA-seq quantification. Nat Biotechnol 34:525–527. doi:10.1038/nbt.3519

Britz O, Zhang J, Grossmann KS, Dyck J, Kim JC, Dymecki S, Gosgnach S, Goulding M. 2015. A genetically defined asymmetry underlies the inhibitory control of flexor–extensor locomotor movements. Elife 4:1–22. doi:10.7554/eLife.04718

Brule CE, Grayhack EJ. 2017. Synonymous codons: choose wisely for expression. Trends Genet 33:283–297. doi:10.1016/j.tig.2017.02.001

Brunkard JO, Baker B. 2018. A two-headed monster to avert disaster: hbs1/ski7 is alternatively spliced to build eukaryotic rna surveillance complexes. Front Plant Sci 9:1–17. doi:10.3389/fpls.2018.01333

Buhr F, Jha S, Thommen M, Mittelstaet J, Kutz F, Schwalbe H, Rodnina M V, Komar AA. 2016. Synonymous codons direct cotranslational folding toward different protein conformations. Mol Cell 61:341–351. doi:10.1016/j.molcel.2016.01.008

Buszczak M, Signer RAJ, Morrison SJ. 2014. Cellular differences in protein synthesis regulate tissue homeostasis. Cell 159:242–251. doi:doi:10.1016/j.cell.2014.09.016.

Carnevalli LS, Pereira CM, Longo BM, Jaqueta CB, Avedissian M, Mello LEAM, Castilho BA. 2004. Phosphorylation of translation initiation factor eIF2α in the brain during pilocarpine-induced status epilepticus in mice. Neurosci Lett 357:191–194. doi:10.1016/j.neulet.2003.12.093

Castelo-Szekely V, Arpat AB, Janich P, Gatfield D. 2017. Translational contributions to tissue specificity in rhythmic and constitutive gene expression. Genome Biol 18:1–17. doi:10.1186/s13059-017-1222-2

Choe Y-J, Park S-H, Hassemer T, Körner R, Vincenz-Donnelly L, Hayer-Hartl M, Hartl FU. 2016. Failure of RQC machinery causes protein aggregation and proteotoxic stress. Nature 531:191–195. doi:10.1038/nature16973

Chu J, Hong NA, Masuda CA, Jenkins B V., Nelms KA, Goodnow CC, Glynne RJ, Wu H, Masliah E, Joazeiro CAP, Kay SA. 2009. A mouse forward genetics screen identifies LISTERIN as an E3 ubiquitin ligase involved in neurodegeneration. Proc Natl Acad Sci U S A 106:2097–2103. doi:10.1073/pnas.0812819106

Chung S-H, Kim C-T, Jung Y-H, Lee N-S, Jeong Y-G. 2010. Early cerebellar granule cell migration in the mouse embryonic development. Anat Cell Biol 43:86. doi:10.5115/acb.2010.43.1.86

Chung SH, Guo F, Jiang P, Pleasure DE, Deng W. 2013. Olig2/Plp-positive progenitor cells give rise to bergmann glia in the cerebellum. Cell Death Dis 4:1–13. doi:10.1038/cddis.2013.74

Collart MA, Weiss B. 2019. Ribosome pausing, a dangerous necessity for co-translational events. Nucleic Acids Res 48:1043–1055. doi:10.1093/nar/gkz763

Cui M, Wang Z, Bassel-Duby R, Olson EN. 2018. Genetic and epigenetic regulation of cardiomyocytes in development, regeneration and disease. Dev 145. doi:10.1242/dev.171983

D’Orazio KN, Wu CCC, Sinha N, Loll-Krippleber R, Brown GW, Green R. 2019. The endonuclease cue2 cleaves mRNAs at stalled ribosomes during no go decay. Elife 8:1–27. doi:10.7554/eLife.49117.001

Diament A, Feldman A, Schochet E, Kupiec M, Arava Y, Tuller T. 2018. The extent of ribosome queuing in budding yeast. PLoS Comput Biol 14:1–21. doi:10.1371/journal.pcbi.1005951

Doma MK, Parker R. 2006. Endonucleolytic cleavage of eukaryotic mRNAs with stalls in translation elongation. Nature 440:561–4. doi:10.1038/nature04530

Dong X, Kwan KM. 2020. Yin Yang 1 is critical for mid-hindbrain neuroepithelium development and involved in cerebellar agenesis. Mol Brain 13:1–18. doi:10.1186/s13041-020-00643-z

Drummond DA, Wilke CO. 2008. Mistranslation-induced protein misfolding as a dominant constraint on coding-sequence evolution. Cell 134:341–352. doi:10.1016/j.cell.2008.05.042

Durinck S, Moreau Y, Kasprzyk A, Davis S, De Moor B, Brazma A, Huber W. 2005. BioMart and Bioconductor: A powerful link between biological databases and microarray data analysis. Bioinformatics 21:3439–3440. doi:10.1093/bioinformatics/bti525

Eberle AB, Lykke-Andersen S, Mühlemann O, Jensen TH. 2009. SMG6 promotes endonucleolytic cleavage of nonsense mRNA in human cells. Nat Struct Mol Biol 16:49–55. doi:10.1038/nsmb.1530

Fink AJ, Englund C, Daza RAM, Pham D, Lau C, Nivison M, Kowalczyk T, Hevner RF. 2006. Development of the Deep Cerebellar Nuclei : Transcription Factors and Cell Migration from the Rhombic Lip. J Neurosci 26:3066–3076. doi:10.1523/JNEUROSCI.5203-05.2006

Fujii K, Shi Z, Zhulyn O, Denans N, Barna M. 2017. Pervasive translational regulation of the cell signalling circuitry underlies mammalian development. Nat Commun 8:1–13. doi:10.1038/ncomms14443

Fünfschilling U, Reichardt LF. 2002. Cre-mediated recombination in rhombic lip derivatives. Genesis 33:160–169. doi:10.1002/gene.10104

Gabut M, Bourdelais F, Durand S. 2020. Ribosome and Translational Control in Stem Cells. Cells 9:497. doi:10.3390/cells9020497

Ge SX, Jung D, Jung D, Yao R. 2020. ShinyGO: A graphical gene-set enrichment tool for animals and plants. Bioinformatics 36:2628–2629. doi:10.1093/bioinformatics/btz931

Glover ML, Burroughs AM, Monem PC, Egelhofer TA, Pule MN, Aravind L, Arribere JA. 2020. NONU-1 encodes a conserved endonuclease required for mRNA translation surveillance. Cell Rep 30:4321–4331. doi:10.1016/j.celrep.2020.03.023

Gonzalez C, Sims JS, Hornstein N, Mela A, Garcia F, Lei L, Gass DA, Amendolara B, Bruce JN, Canoll P, Sims PA. 2014. Ribosome profiling reveals a cell-type-specific translational landscape in brain tumors. J Neurosci 34:10924–10936. doi:10.1523/JNEUROSCI.0084-14.2014

Götz M, Huttner WB. 2005. The cell biology of neurogenesis. Nat Rev Mol Cell Biol. doi:https://doi.org/10.1038/nrm1739

Guo Q, Li K, Sunmonu NA, Li JYH. 2010. Fgf8b-containing spliceforms, but not Fgf8a, are essential for Fgf8 function during development of the midbrain and cerebellum. Dev Biol 338:183–192. doi:10.1016/j.ydbio.2009.11.034

Guydosh NR, Green R. 2017. Translation of poly(A) tails leads to precise mRNA cleavage. RNA 23:749–761. doi:10.1261/rna.060418.116

Guydosh NR, Green R. 2014. Dom34 rescues ribosomes in 3′ untranslated regions. Cell 156:950–962. doi:10.1016/j.cell.2014.02.006

Guydosh NR, Kimmig P, Walter P, Green R. 2017. Regulated Ire1-dependent mRNA decay requires no-go mRNA degradation to maintain endoplasmic reticulum homeostasis in S. Pombe. Elife 6:e29216. doi:10.7554/eLife.29216

Han P, Shichino Y, Schneider-Poetsch T, Mito M, Hashimoto S, Udagawa T, Kohno K, Yoshida M, Mishima Y, Inada T, Iwasaki S. 2020. Genome-wide Survey of Ribosome Collision. Cell Rep 31:107610. doi:10.1016/j.celrep.2020.107610

Hartman NW, Lin T V., Zhang L, Paquelet GE, Feliciano DM, Bordey A. 2013. MTORC1 Targets the Translational Repressor 4E-BP2, but Not S6 Kinase 1/2, to Regulate Neural Stem Cell Self-Renewal InVivo. Cell Rep 5:433–444. doi:10.1016/j.celrep.2013.09.017

Hashimoto Y, Takahashi M, Sakota E, Nakamura Y. 2017. Nonstop-mRNA decay machinery is involved in the clearance of mRNA 5′-fragments produced by RNAi and NMD in Drosophila melanogaster cells. Biochem Biophys Res Commun 484:1–7. doi:10.1016/j.bbrc.2017.01.092

He F, Li X, Spatrick P, Casillo R, Dong S, Jacobson A. 2003. Genome-Wide Analysis of mRNAs Regulated by the Nonsense-Mediated and 5′ to 3′ mRNA Decay Pathways in Yeast. Mol Cell 12:1439–1452. doi:10.1016/S1097-2765(03)00446-5

Hilal T, Yamamoto H, Loerke J, Bürger J, Mielke T, Spahn CMT. 2016. Structural insights into ribosomal rescue by Dom34 and Hbs1 at near-atomic resolution. Nat Commun 7:1–8. doi:10.1038/ncomms13521

Hu W, Sweet TJ, Chamnongpol S, Baker KE, Coller J. 2009. Co-translational mRNA decay in Saccharomyces cerevisiae. Nature 461:225–229. doi:10.1038/nature08265

Huang L, Shum E, Jones S, Lou C-H, Dumdie J, Kim H, Roberts A, Espinoza J, Skarbrevik D, Phan M, Cook-Andersen H, Swerdlow N, Gecz J, Wilkinson M. 2017. A Upf3b-mutant mouse model with behavioral and neurogenesis defects. Mol Psychiatry 23:1773–1786. doi:10.1038/mp.2017.173

Ibrahim F, Maragkakis M, Alexiou P, Mourelatos Z. 2018. Ribothrypsis, a novel process of canonical mRNA decay, mediates ribosome-phased mRNA endonucleolysis. Nat Struct Mol Biol 25:302–310. doi:10.1038/s41594-018-0042-8

Ingolia NT, Brar GA, Rouskin S, McGeachy AM, Weissman JS. 2012. The ribosome profiling strategy for monitoring translation in vivo by deep sequencing of ribosome-protected mRNA fragments. Nat Protoc 7:1534–1550. doi:10.1038/nprot.2012.086

Ishimura R, Nagy G, Dotu I, Chuang JH, Ackerman SL. 2016. Activation of GCN2 kinase by ribosome stalling links translation elongation with translation initiation. Elife 5:1–22. doi:10.7554/eLife.14295

Ishimura R, Nagy G, Dotu I, Zhou H, Yang XL, Schimmel P, Senju S, Nishimura Y, Chuang JH, Ackerman SL. 2014. Ribosome stalling induced by mutation of a CNS-specific tRNA causes neurodegeneration. Science (80-) 345:455–459. doi:10.1126/science.1249749

Jaffrey S, Wilkinson MF. 2018. Nonsense-mediated RNA decay in the brain: Emerging modulator of neural development and disease. Nat Rev Neurosci.

Jamar NH, Kritsiligkou P, Grant CM. 2018. Loss of mRNA surveillance pathways results in widespread protein aggregation. Sci Rep 8:1–10. doi:10.1038/s41598-018-22183-2

Jiang X, Feng S, Chen Y, Feng Y, Deng H. 2016. Proteomic analysis of mTOR inhibition-mediated phosphorylation changes in ribosomal proteins and eukaryotic translation initiation factors. Protein Cell 7:533–537. doi:10.1007/s13238-016-0279-0

Johnson JL, Stoica L, Liu Y, Zhu PJ, Bhattacharya A, Buffington SA, Huq R, Eissa NT, Larsson O, Porse BT, Domingo D, Nawaz U, Carroll R, Jolly L, Scerri TS, Kim HG, Brignell A, Coleman MJ, Braden R, Kini U, Jackson V, Baxter A, Bahlo M, Scheffer IE, Amor DJ, Hildebrand MS, Bonnen PE, Beeton C, Gecz J, Morgan AT, Costa-Mattioli M. 2019. Inhibition of Upf2-Dependent Nonsense-Mediated Decay Leads to Behavioral and Neurophysiological Abnormalities by Activating the Immune Response. Neuron 104:665–679.e8. doi:10.1016/j.neuron.2019.08.027

Ju J, Liu Q, Zhang Y, Liu Y, Jiang M, Zhang L, He X, Peng C, Zheng T, Lu QR, Li H. 2016. Olig2 regulates Purkinje cell generation in the early developing mouse cerebellum. Sci Rep 6:1–11. doi:10.1038/srep30711

Juszkiewicz S, Speldewinde SH, Wan L, Svejstrup JQ, Hegde RS. 2020a. The ASC-1 complex disassembles collided ribosomes. Mol Cell 79:603–614. doi:10.1016/j.molcel.2020.06.006

Juszkiewicz S, Speldewinde SH, Wan L, Svejstrup JQ, Hegde RS. 2020b. The ASC-1 complex disassembles collided ribosomes. Mol Cell 79:603–614. doi:10.1016/j.molcel.2020.06.006

Kalisiak K, Kuliński TM, Tomecki R, Cysewski D, Pietras Z, Chlebowski A, Kowalska K, Dziembowski A. 2017. A short splicing isoform of HBS1L links the cytoplasmic exosome and SKI complexes in humans. Nucleic Acids Res 45:2068–2080. doi:10.1093/nar/gkw862

Kapur M, Ganguly A, Nagy G, Adamson SI, Chuang JH, Frankel WN, Ackerman SL. 2020. Expression of the neuronal tRNA n-Tr20 regulates synaptic transmission and seizure susceptibility. Neuron 108:1–16. doi:10.1016/j.neuron.2020.07.023

Karousis ED, Mühlemann O. 2019. Nonsense-mediated mRNA decay begins where translation ends. Cold Spring Harb Perspect Biol 11:1–18. doi:10.1101/cshperspect.a032862

Kim D, Paggi JM, Park C, Bennett C, Salzberg SL. 2019. Graph-based genome alignment and genotyping with HISAT2 and HISAT-genotype. Nat Biotechnol 37:907–915. doi:10.1038/s41587-019-0201-4

Kim J, Kundu M, Viollet B, Guan K-L. 2011. AMPK and mTOR regulate autophagy through direct phosphorylation of Ulk1. Nat Cell Biol. doi:10.1038/ncb2152

Kimmel RA, Turnbull DH, Blanquet V, Wurst W, Loomis CA, Joyner AL. 2000. Two lineage boundaries coordinate vertebrate apical ectodermal ridge formation. Genes Dev 14:1377– 1389. doi:10.1101/gad.14.11.1377

Knoepfler PS, Cheng PF, Eisenman RN. 2002. N-myc is essential during neurogenesis for the rapid expansion of progenitor cell populations and the inhibition of neuronal differentiation. Genes Dev 16:2699–2712. doi:10.1101/gad.1021202

Kong J, Lasko P. 2012. Translational control in cellular and developmental processes. Nat Rev Genet 13:383–394. doi:10.1038/nrg3184

Kristensen AR, Gsponer J, Foster LJ. 2013. Protein synthesis rate is the predominant regulator of protein expression during differentiation. Mol Syst Biol 9:1–12. doi:10.1038/msb.2013.47

Langmead B, Salzberg SL. 2012. Fast gapped-read alignment with Bowtie 2. Nat Methods 9:357–359. doi:10.1038/nmeth.1923

Lauria F, Tebaldi T, Bernabò P, Groen EJN, Gillingwater TH, Viero G. 2018. riboWaltz: Optimization of ribosome P-site positioning in ribosome profiling data. PLoS Comput Biol 14:1–20. doi:10.1371/journal.pcbi.1006169

Legué E, Gottshall JL, Jaumouillé E, Roselló-Díez A, Shi W, Barraza LH, Washington S, Grant RL, Joyner AL. 2016. Differential timing of granule cell production during cerebellum development underlies generation of the foliation pattern. Neural Dev 11:1–14. doi:10.1186/s13064-016-0072-z

Lelivelt MJ, Culbertson MR. 1999. Yeast Upf Proteins Required for RNA Surveillance Affect Global Expression of the Yeast Transcriptome. Mol Cell Biol 19:6710–6719. doi:10.1128/mcb.19.10.6710

Leto K, Rolando C, Rossi F. 2012. The Genesis of Cerebellar GABAergic Neurons: Fate Potential and Specification Mechanisms. Front Neuroanat 6:1–10. doi:10.3389/fnana.2012.00006

Li JYH, Lao Z, Joyner AL. 2002. Changing requirements for Gbx2 in development of the cerebellum and maintenance of the Mid/hindbrain organizer. Neuron 36:31–43. doi:10.1016/S0896-6273(02)00935-2

Li T, Shi Y, Wang P, Guachalla LM, Sun B, Joerss T, Chen Y, Groth M, Krueger A, Platzer M, Yang Y, Rudolph KL, Wang Z. 2015. Smg6/Est1 licenses embryonic stem cell differentiation via nonsense-mediated mRNA decay . EMBO J 34:1630–1647. doi:10.15252/embj.201489947

Li W, Wang W, Uren PJ, Penalva LOF, Smith AD. 2017. Riborex: Fast and flexible identification of differential translation from Ribo-seq data. Bioinformatics 33:1735–1737. doi:10.1093/bioinformatics/btx047

Liakath-Ali K, Mills EW, Sequeira I, Lichtenberger BM, Pisco AO, Sipilä KH, Mishra A, Yoshikawa H, Wu CCC, Ly T, Lamond AI, Adham IM, Green R, Watt FM. 2018. An evolutionarily conserved ribosome-rescue pathway maintains epidermal homeostasis. Nature 556:376–380. doi:10.1038/s41586-018-0032-3

Liao Y, Smyth GK, Shi W. 2014. FeatureCounts: An efficient general purpose program for assigning sequence reads to genomic features. Bioinformatics 30:923–930. doi:10.1093/bioinformatics/btt656

Licausi F, Hartman NW. 2018. Role of mTOR complexes in neurogenesis. Int J Mol Sci 19. doi:10.3390/ijms19051544

Liu R, Iadevaia V, Averous J, Taylor PM, Zhang Z, Proud CG. 2014. Impairing the production of ribosomal RNA activates mammalian target of rapamycin complex 1 signalling and downstream translation factors. Nucleic Acids Res 42:5083–5096. doi:10.1093/nar/gku130

Liu Y, Beyer A, Aebersold R. 2016. On the dependency of cellular protein levels on mRNA abundance. Cell 165:535–550. doi:10.1016/j.cell.2016.03.014

Livak KJ, Schmittgen TD. 2001. Analysis of relative gene expression data using real-time quantitative PCR and the 2-ΔΔCT method. Methods 25:402–408. doi:10.1006/meth.2001.1262

Lorenz A, Deutschmann M, Ahlfeld J, Prix C, Koch A, Smits R, Fodde R, Kretzschmar HA, Schuller U. 2011. Severe Alterations of Cerebellar Cortical Development after Constitutive Activation of Wnt Signaling in Granule Neuron Precursors. Mol Cell Biol 31:3326–3338. doi:10.1128/mcb.05718-11

Love MI, Huber W, Anders S. 2014. Moderated estimation of fold change and dispersion for RNA-seq data with DESeq2. Genome Biol 15:1–21. doi:10.1186/s13059-014-0550-8

Ma M, Wu W, Li Q, Li J, Sheng Z, Shi J, Zhang M, Yang H, Wang Z, Sun R, Fei J. 2015. N-myc is a key switch regulating the proliferation cycle of postnatal cerebellar granule cell progenitors. Sci Rep 5:1–13. doi:10.1038/srep12740

Magri L, Cambiaghi M, Cominelli M, Alfaro-Cervello C, Cursi M, Pala M, Bulfone A, Garca-Verdugo JM, Leocani L, Minicucci F, Poliani PL, Galli R. 2011. Sustained activation of mTOR pathway in embryonic neural stem cells leads to development of tuberous sclerosis complex-associated lesions. Cell Stem Cell 9:447–462. doi:10.1016/j.stem.2011.09.008

Maier T, Güell M, Serrano L. 2009. Correlation of mRNA and protein in complex biological samples. FEBS Lett 583:3966–3973. doi:10.1016/j.febslet.2009.10.036

Marshall AN, Han J, Kim M, Van Hoof A. 2018. Conservation of mRNA quality control factor Ski7 and its diversification through changes in alternative splicing and gene duplication. Proc Natl Acad Sci U S A 115:E6808–E6816. doi:10.1073/pnas.1801997115

Martin PB, Kigoshi-Tansho Y, Sher RB, Ravenscroft G, Stauffer JE, Kumar R, Yonashiro R, Müller T, Griffith C, Allen W, Pehlivan D, Haral T, Zenker M, Howting D, Schanze D, Faqeih EA, Almontashiri NAM, Maroofian R, Houlden H, Mazaheri N, Galehdari H, Douglas G, Posey JE, Ryan M, Lupski JR, Laing NG, Joazeiro CAP, Cox GA. 2020. NEMF mutations that impair ribosome-associated quality control are associated with neuromuscular disease. Nat Commun 11:1–12. doi:10.1038/s41467-020-18327-6

Mayer C, Grummt I. 2006. Ribosome biogenesis and cell growth: mTOR coordinates transcription by all three classes of nuclear RNA polymerases. Oncogene 25:6384–6391. doi:10.1038/sj.onc.1209883

Mendell JT, Sharifi NA, Meyers JL, Martinez-Murillo F, Dietz HC. 2004. Nonsense surveillance regulates expression of diverse classes of mammalian transcripts and mutes genomic noise. Nat Genet 36:1073–1078. doi:10.1038/ng1429

Meng D, Frank AR, Jewell JL. 2018. mTOR signaling in stem and progenitor cells. Development 145. doi:10.1242/dev.152595

Meydan S, Guydosh NR. 2020. Disome and trisome profiling reveal genome-wide targets of ribosome quality control. Mol Cell 79:588–602. doi:10.1016/j.molcel.2020.06.010

Mills EW, Green R, Ingolia NT. 2017. Slowed decay of mRNAs enhances platelet specific translation. Blood 129:e38–e48. doi:10.1182/blood-2016-08-736108

Mills EW, Wangen J, Green R, Ingolia NT. 2016. Dynamic Regulation of a Ribosome Rescue Pathway in Erythroid Cells and Platelets. Cell Rep 17:1–10. doi:10.1016/j.celrep.2016.08.088

Molkentin JD, Lin Q, Duncan SA, Olson EN. 1997. Requirement of the transcription factor GATA4 for heart tube formation and ventral morphogenesis. Genes Dev 11:1061–1072. doi:10.1101/gad.11.8.1061

Muñoz-Martín N, Sierra R, Schimmang T, Del Campo CV, Torres M. 2019. Myc is dispensable for cardiomyocyte development but rescues Mycn-deficient hearts through functional replacement and cell competition. Dev 146:1–7. doi:10.1242/dev.170753

Nagy A, Gertsenstein M, Vintersten K, Behringer R. 2014. Manipulating the mouse embryo - A laboraty manual. Cold Spring Harb Lab Press 4th.

Notaras M, Allen M, Longo F, Volk N, Toth M, Jeon NL, Klann E, Colak D. 2019. UPF2 leads to degradation of dendritically targeted mRNAs to regulate synaptic plasticity and cognitive function. Mol Psychiatry. doi:10.1038/s41380-019-0547-5

O’Connell AE, Gerashchenko M V., O’Donohue MF, Rosen SM, Huntzinger E, Gleeson D, Galli A, Ryder E, Cao S, Murphy Q, Kazerounian S, Morton SU, Schmitz-Abe K, Gladyshev VN, Gleizes PE, Séraphin B, Agrawal PB. 2019. Mammalian Hbs1L deficiency causes congenital anomalies and developmental delay associated with Pelota depletion and 80S monosome accumulation. PLoS Genet 15:1–25. doi:10.1371/journal.pgen.1007917

Pan N, Jahan I, Lee JE, Fritzsch B. 2009. Defects in the cerebella of conditional Neurod1 null mice correlate with effective Tg(Atoh1-cre) recombination and granule cell requirements for Neurod1 for differentiation. Cell Tissue Res 337:407–428. doi:10.1007/s00441-009-0826-6

Pimentel H, Bray NL, Puente S, Melsted P, Pachter L. 2017. Differential analysis of RNA-seq incorporating quantification uncertainty. Nat Methods 14:687–690. doi:10.1038/nmeth.4324

Pisareva VP, Skabkin MA, Hellen CUT, Pestova T V., Pisarev A V. 2011. Dissociation by Pelota, Hbs1 and ABCE1 of mammalian vacant 80S ribosomes and stalled elongation complexes. EMBO J 30:1804–1817. doi:10.1038/emboj.2011.93

Puertollano R. 2014. mTOR and lysosome regulation. F1000Prime Rep 6:1–7. doi:10.12703/P6-52

Qiu Z, Cang Y, Goff SP. 2010. Abl family tyrosine kinases are essential for basement membrane integrity and cortical lamination in the cerebellum. J Neurosci 30:14430–14439. doi:10.1523/JNEUROSCI.2861-10.2010

Rabanal-Ruiz Y, Korolchuk VI. 2018. mTORC1 and nutrient homeostasis: The central role of the lysosome. Int J Mol Sci 19. doi:10.3390/ijms19030818

Rainer J, Gatto L, Weichenberger CX. 2019. Ensembldb: An R package to create and use Ensembl-based annotation resources. Bioinformatics 35:3151–3153. doi:10.1093/bioinformatics/btz031

Roczniak-Ferguson A, Petit CS, Froehlich F, Qian S, Ky J, Angarola B, Walther TC, Ferguson SM. 2012. The Transcription Factor TFEB Links mTORC1 Signaling to Transcriptional Control of Lysosome Homeostasis. Sci Signal. doi:10.1126/scisignal.2002790

Rodrigues DC, Mufteev M, Weatheritt RJ, Djuric U, Ha KCH, Ross PJ, Wei W, Piekna A, Sartori MA, Byres L, Mok RSF, Zaslavsky K, Pasceri P, Diamandis P, Morris Q, Blencowe BJ, Ellis J. 2020. Shifts in Ribosome Engagement Impact Key Gene Sets in Neurodevelopment and Ubiquitination in Rett Syndrome. Cell Rep 30:4179–4196.e11. doi:10.1016/j.celrep.2020.02.107

Sapir T, Geiman EJ, Wang Z, Velasquez T, Mitsui S, Yoshihara Y, Frank E, Alvarez FJ, Goulding M. 2004. Pax6 and Engrailed 1 Regulate Two Distinct Aspects of Renshaw Cell Development. J Neurosci 24:1255–1264. doi:10.1523/JNEUROSCI.3187-03.2004

Schuller AP, Green R. 2019. Roadblocks and resolutions in eukaryotic translation. Nat Rev Mol Cell Biol 19:526–541. doi:10.1038/s41580-018-0011-4

Serin G, Gersappe A, Black JD, Aronoff R, Maquat LE. 2001. Identification and Characterization of Human Orthologues to Saccharomyces cerevisiae Upf2 Protein and Upf3 Protein (Caenorhabditis elegans SMG-4). Mol Cell Biol 21:209–223. doi:10.1128/mcb.21.1.209-223.2001

Sgaier SK, Lao Z, Villanueva MP, Berenshteyn F, Stephen D, Turnbull RK, Joyner AL. 2007. Genetic subdivision of the tectum and cerebellum into functionally related regions based on differential sensitivity to engrailed proteins. Development. doi:10.1242/dev.000620

Shoemaker CJ, Green R. 2012. Translation drives mRNA quality control. Nat Struct Mol Biol 19:594–601. doi:10.1038/nsmb.2301

Shoemaker CJ, Green R. 2011. Kinetic analysis reveals the ordered coupling of translation termination and ribosome recycling in yeast. Proc Natl Acad Sci U S A 108:E1392–8. doi:10.1073/pnas.1113956108

Signer RAJ, Magee JA, Salic A, Morrison SJ. 2014. Haematopoietic stem cells require a highly regulated protein synthesis rate. Nature 508:49–54. doi:10.1038/nature13035

Simms CL, Hudson BH, Mosior JW, Rangwala AS, Zaher HS. 2014. An Active Role for the Ribosome in Determining the Fate of Oxidized mRNA. Cell Rep 9:1256–1264. doi:10.1016/j.celrep.2014.10.042

Simms CL, Thomas EN, Zaher HS. 2017. Ribosome-based quality control of mRNA and nascent peptides. Wiley Interdiscip Rev RNA. doi:10.1002/wrna.1366

Skarnes WC, Rosen B, West AP, Koutsourakis M, Bushell W, Iyer V, Mujica AO, Thomas M, Harrow J, Cox T, Jackson D, Severin J, Biggs P, Fu J, Nefedov M, De Jong PJ, Stewart AF, Bradley A. 2011. A conditional knockout resource for the genome-wide study of mouse gene function. Nature 474:337–344. doi:10.1038/nature10163

Spencer PS, Siller E, Anderson JF, Barral JM. 2012. Silent substitutions predictably alter translation elongation rates and protein folding efficiencies. J Mol Biol 422:328–335. doi:10.1016/j.jmb.2012.06.010

Steimle JD, Moskowitz IP. 2017. TBX5: A Key Regulator of Heart Development. Curr Top Dev Biol 122:195–221. doi:10.1016/bs.ctdb.2016.08.008

Sudmant PH, Lee H, Dominguez D, Heiman M, Burge CB. 2018. Widespread accumulation of ribosome-associated isolated 3′ UTRs in neuronal cell populations of the aging brain. Cell Rep 25:2447–2456. doi:10.1016/j.celrep.2018.10.094

Tahmasebi S, Amiri M, Sonenberg N. 2019. Translational control in stem cells. Front Genet 10:1–9. doi:10.3389/fgene.2018.00709

Tatsumi K, Isonishi A, Yamasaki M, Kawabe Y, Morita-Takemura S, Nakahara K, Terada Y, Shinjo T, Okuda H, Tanaka T, Wanaka A. 2018. Olig2-Lineage astrocytes: A distinct subtype of astrocytes that differs from GFAP astrocytes. Front Neuroanat 12:1–17. doi:10.3389/fnana.2018.00008

Terrey M, Adamson SI, Gibson AL, Deng T, Ishimura R, Chuang JH, Ackerman SL. 2020. GTPBP1 resolves paused ribosomes to maintain neuronal homeostasis. Elife 9:1–22. doi:10.7554/eLife.62731

Thommen M, Holtkamp W, Rodnina M V. 2017. Co-translational protein folding: progress and methods. Curr Opin Struct Biol 42:83–89. doi:10.1016/j.sbi.2016.11.020

Tripathi PP, Di Giovannantonio LG, Viegi A, Wurst W, Simeone A, Bozzi Y. 2008. Serotonin hyperinnervation abolishes seizure susceptibility in Otx2 conditional mutant mice. J Neurosci 28:9271–9276. doi:10.1523/JNEUROSCI.2208-08.2008

Vong KI, Leung CKY, Behringer RR, Kwan KM. 2015. Sox9 is critical for suppression of neurogenesis but not initiation of gliogenesis in the cerebellum. Mol Brain 8:0–17. doi:10.1186/s13041-015-0115-0

Way SW, Mckenna J, Mietzsch U, Reith RM, Wu HCJ, Gambello MJ. 2009. Loss of Tsc2 in radial glia models the brain pathology of tuberous sclerosis complex in the mouse. Hum Mol Genet 18:1252–1265. doi:10.1093/hmg/ddp025

Weischenfeldt J, Damgaard I, Bryder D, Theilgaard-Mönch K, Thoren LA, Nielsen FC, Jacobsen SEW, Nerlov C, Porse BT. 2008. NMD is essential for hematopoietic stem and progenitor cells and for eliminating by-products of programmed DNA rearrangements. Genes Dev 22:1381–1396. doi:10.1101/gad.468808

Wey A, Cerdeno VM, Pleasure D, Knoepfler PS. 2010. c- and N-myc regulate neural precursor cell fate, cell cycle, and metabolism to direct cerebellar development. Cerebellum 9:537– 547. doi:10.1007/s12311-010-0190-9

Wizeman JW, Guo Q, Wilion EM, Li JYH. 2019. Specification of diverse cell types during early neurogenesis of the mouse cerebellum. Elife 8:1–24. doi:10.7554/eLife.42388

Wojcinski A, Morabito M, Lawton AK, Stephen DN, Joyner AL. 2019. Genetic deletion of genes in the cerebellar rhombic lip lineage can stimulate compensation through adaptive reprogramming of ventricular zone-derived progenitors. Neural Dev 14:1–15. doi:10.1186/s13064-019-0128-y

Wolf AS, Grayhack EJ. 2015. Asc1, homolog of human RACK1, prevents frameshifting in yeast by ribosomes stalled at CGA codon repeats. RNA 21:935–945. doi:10.1261/rna.049080.114

Wurst W, Auerbach AB, Joyner AL. 1994. Multiple developmental defects in Engrailed-1 mutant mice: An early mid-hindbrain deletion and patterning defects in forelimbs and sternum. Development 120:2065–2075.

Xiang X, Zhao J, Xu G, Li Y, Zhang W. 2011. mTOR and the differentiation of mesenchymal stem cells. Acta Biochim Biophys Sin (Shanghai) 43:501–510. doi:10.1093/abbs/gmr041

Yamashita R, Suzuki Y, Takeuchi N, Wakaguri H, Ueda T, Sugano S, Nakai K. 2008. Comprehensive detection of human terminal oligo-pyrimidine (TOP) genes and analysis of their characteristics. Nucleic Acids Res 36:3707–3715. doi:10.1093/nar/gkn248

Yonashiro R, Tahara EB, Bengtson MH, Khokhrina M, Lorenz H, Chen KC, Kigoshi-Tansho Y, Savas JN, Yates JR, Kay SA, Craig EA, Mogk A, Bukau B, Joazeiro CAP. 2016. The Rqc2/Tae2 subunit of the ribosome-associated quality control (RQC) complex marks ribosome-stalled nascent polypeptide chains for aggregation. Elife 5:1–16. doi:10.7554/eLife.11794

Young DJ, Guydosh NR, Zhang F, Hinnebusch AG, Green R. 2015. Rli1/ABCE1 Recycles Terminating Ribosomes and Controls Translation Reinitiation in 3′UTRs In Vivo. Cell 162:872–884. doi:10.1016/j.cell.2015.07.041

Yu CH, Dang Y, Zhou Z, Wu C, Zhao F, Sachs MS, Liu Y. 2015. Codon usage influences the local rate of translation elongation to regulate co-translational protein folding. Mol Cell 59:744–754. doi:10.1016/j.molcel.2015.07.018

Zhang Y, Chen K, Sloan SA, Bennett ML, Scholze AR, O’Keeffe S, Phatnani HP, Guarnieri P, Caneda C, Ruderisch N, Deng S, Liddelow SA, Zhang C, Daneman R, Maniatis T, Barres BA, Wu JQ. 2014. An RNA-sequencing transcriptome and splicing database of glia, neurons, and vascular cells of the cerebral cortex. J Neurosci 34:11929–11947. doi:10.1523/JNEUROSCI.1860-14.2014

